# Multilayer MEG source modelling enables depth-resolved laminar inference in humans

**DOI:** 10.1101/2025.05.28.656642

**Authors:** Maciej J. Szul, Matteo Maspoli, Danila Shelepenkov, Ishita Agarwal, Quentin Moreau, Solène Gailhard, Maxime Ferez, Carolina Fernandez Pujol, Yunkai Zhu, Bassem Hiba, Sebastien Daligault, Franck Lamberton, Denis Schwartz, Jérémie Mattout, Mathilde Bonnefond, Andrew R. Dykstra, Sven Bestmann, Gareth R. Barnes, James J. Bonaiuto

## Abstract

Neural dynamics at the laminar level are critical for cortical computation. However, in humans, non-invasive methods to probe such dynamics have been limited to coarse distinctions between deep and superficial layers. Here, we present a multilayer magnetoencephalography source reconstruction framework and evaluate the conditions under which depth-resolved laminar inference may be feasible. Using simulations, we systematically assess the limits of magnetoencephalography depth resolution, showing that laminar discrimination depends on sufficiently high signal-to-noise ratio, precise co-registration, and accurate specification of cortical column orientation. We demonstrate that regional variations in cortical anatomy influence reconstruction fidelity, with lead-field separability emerging as a key determinant. We then apply this framework to empirical data from three independent datasets and find laminar activation patterns that align with canonical feedforward and feedback motifs in visual and sensorimotor circuits, supporting the plausibility of laminar inference under favorable conditions and offering opportunities to bridge invasive electrophysiology and human neuroimaging.

## INTRODUCTION

The morphology, connectivity, and computations of cortical laminae have been central topics in neuroscience for over a century^1^. Invasive laminar electrophysiology has provided fundamental insights into sensory and sensorimotor computations - e.g., laminar specific receptive fields^2^ and the directional flow of information in feed-forward and feedback pathways^3^ - that would be otherwise inaccessible. In cognitive neuroscience, canonical microcircuit models and other laminar interaction frameworks have generated mechanistic hypotheses about lamina-specific contributions to perception, cognition, and conscious processing^4–8^. Clinically, assessment of laminar dynamics has improved our understanding of epileptogenic propagation^9^ and holds promise for refining our knowledge of disorders such as Parkinson’s disease^10^, but these investigations have been mostly limited to invasive methods.

Invasive electrophysiological recordings are widely used in animal models, whereas they are restricted to specific clinical populations in humans^11^. Traditional invasive recordings provide direct access to neuronal events, capturing single-unit and local population activity with high spatial and temporal resolution, though newer technologies such as Neuropixels now extend this access across entire cortical columns and brain structures^12^. However, their use requires surgical implantation, necessitating ethical considerations in animal research, and medical justification in human studies. Furthermore, although invasive recordings enable laminar-level recordings in cortex, they remain extracellular and thus are still subject to an inverse problem^13^. This limitation has motivated recent efforts to develop magnetrodes, which aim to directly measure intracellular currents^14^. Nevertheless, these invasive methods are not feasible for broader applications in cognitive neuroscience. In contrast, for both practical and ethical considerations, non-invasive methods like electroencephalography (EEG), magnetoencephalography (MEG), and functional magnetic resonance imaging (fMRI) dominate human neuroscience^15,16^. Their whole brain coverage enables the study of network-level processing, but each modality has inherent limitations. EEG and MEG provide high temporal resolution, but suffer from an inverse problem and are thought to lack the spatial specificity needed for laminar inference^16,17^, whereas fMRI has excellent spatial, but poor temporal resolution^15^. Recent advances in high resolution fMRI have enabled laminar-level analysis, but its inherently limited temporal resolution precludes the study of laminar dynamics^18^.

MEG has traditionally been considered incapable of resolving sources with a precision at or below the thickness of the cortex^17^. However, this assumption has been challenged by studies indicating that there is no fundamental theoretical barrier to achieving high spatial resolution with MEG^19,20^; the primary limitation is the practical issue of signal-to-noise ratio (SNR). With sufficiently high SNR, MEG can, in principle, resolve sources with much finer precision than traditionally assumed^19,20^. Achieving high signal quality in actual recordings requires minimizing co-registration errors, between-session head position variability, and within-session movement, all of which can be addressed using customized head-casts or 3D-printed helmets that stabilize the head relative to the sensor array, an approach called high precision MEG (hpMEG)^21–23^. hpMEG spurred the development of early laminar MEG approaches, which involved Bayesian model comparison between separate forward models for two cortical depths: deep and superficial, or analysis of source signals using a forward model combining surfaces for the two layers^19,20,24^. These approaches have been used to successfully detect laminar differences predicted by theoretical models and invasive recordings, such as lamina-specific spectral activity^24^ and distinct laminar signatures of beta bursts^25^. However, previous studies have been limited to coarse deep versus superficial distinctions.

Here, we extend these approaches by introducing a multilayer hpMEG source reconstruction framework that moves beyond binary deep-versus-superficial comparisons toward depth-resolved inference across the cortical column. We expand the source space to include an arbitrary number of intermediate layers. We refer to the equidistant depth surfaces used in our forward models as layers (denoted with Arabic numerals, i.e. 1-11), while we use laminae (denoted with Roman numerals) to refer to cytoarchitectonically defined cortical layers (i.e. I-VI). We first use simulations to systematically characterize the conditions under which laminar inference is feasible, quantifying the influence of SNR, co-registration accuracy, dipole orientation, and anatomical variability across the cortex. This extends prior work by moving from two-layer model comparison^20^ to a finer-grained depth-resolved framework and by explicitly identifying the anatomical and data-quality constraints that govern inference accuracy.

Importantly, we distinguish between theoretical feasibility and empirical validation. While simulations allow assessment of laminar inference accuracy under controlled conditions, empirical data lack ground-truth laminar labels. Accordingly, our empirical analyses focus on whether multilayer hpMEG recovers expected laminar-family activation patterns in well-characterized event-related fields (ERFs). In the visual cortex, hpMEG recovers a feedforward activation sequence consistent with laminar models of sensory processing. In the somatosensory and motor cortices, analysis of two independent datasets reveals broadly consistent deep-superficial activation patterns characteristic of motor execution. These results demonstrate that multilayer hpMEG extends non-invasive laminar inference beyond coarse depth distinctions under favorable conditions, while highlighting the constraints that currently limit precise identification of individual laminae *in vivo*. This positions hpMEG as a powerful tool for interrogating neural circuits at a laminar level, offering new opportunities to study how computations within and across cortical laminae shape network-level interactions underlying cognition and behavior^6^.

## RESULTS

### Overview of the multilayer hpMEG framework

We developed a multilayer high-precision MEG (hpMEG) framework for depth-resolved laminar inference that extends previous approaches based on binary deep-versus-superficial comparisons. The framework combines anatomically informed multilayer cortical surface models, Bayesian source reconstruction, and hierarchical model comparison to estimate the cortical depth most consistent with the measured MEG signals (Figure 1). Subject-specific cortical surfaces spanning the full cortical thickness were generated from MRI and used to construct forward models representing different depths within the cortical column. Source reconstruction was then performed separately for each depth model, and model evidence was compared to infer the most probable laminar origin of the activity.

**Figure 1.**
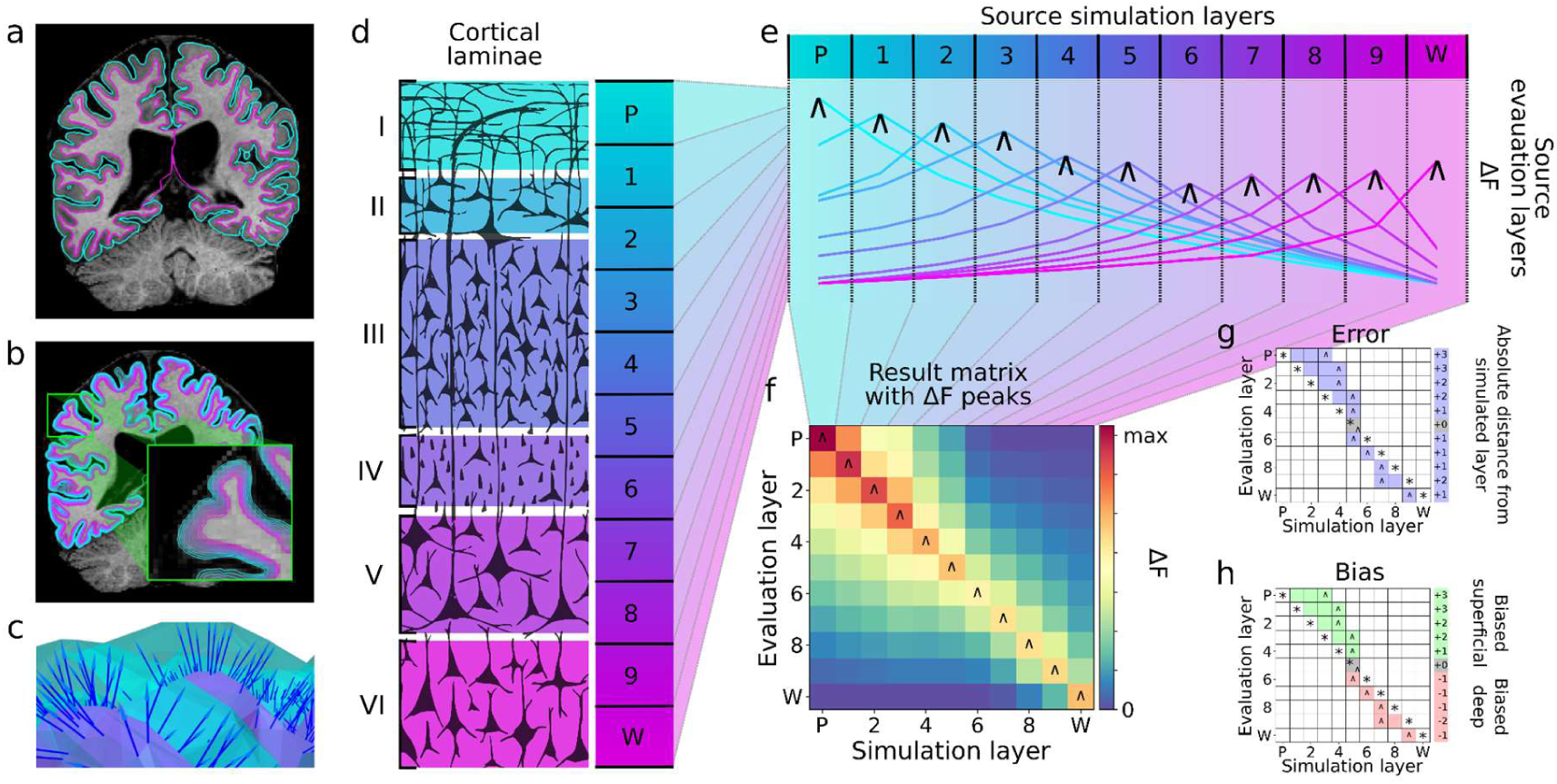
Overview of the simulation strategy and analysis. **a** Pial and white matter boundaries surfaces are extracted from anatomical magnetic resonance imaging (MRI) volumes. **b** Intermediate equidistant surfaces are generated between the pial and white matter surfaces (labeled as superficial (S) and deep (D) respectively). **c** Surfaces are downsampled together, maintaining vertex correspondence across layers. Dipole orientations are constrained using vectors linking corresponding vertices (link vectors; represented by blue lines). **d** The thickness of cortical laminae varies across the cortical depth, which is evenly sampled by the equidistant source surface layers. **e** Each colored line represents the model evidence (relative to the worst model, ΔF) over source layer models, for a signal simulated at a particular layer (the simulated layer is indicated by the line color). The source layer model with the maximal ΔF is indicated by a caret. **f** Result matrix summarizing ΔF across simulated source locations, with peak relative model evidence marked with a caret. **g** Error is calculated from the result matrix as the absolute distance in mm or layers from the simulated source (asterisk) to the peak ΔF (caret). **h** Bias is calculated as the relative position of the peak ΔF (caret) to the simulated source (asterisk) in layers or mm.

We first evaluated the feasibility and limitations of this approach using simulations that systematically varied signal-to-noise ratio, co-registration accuracy, cortical column orientation, and anatomical characteristics. Additional robustness analyses examined the effects of realistic noise structure, concurrent sources, and forward-model mismatch. We then applied the framework to empirical visual and sensorimotor event-related fields (ERFs) to determine whether the recovered laminar activation sequences were consistent with established physiological models of cortical processing. The overall workflow is summarized in Figure 1.

### SNR and co-registration constraints on laminar inference

To determine the SNR required for accurate laminar reconstruction across multiple cortical depths, we selected 100 random cortical locations (vertices on the pial surface). At each location, we ran 11 simulations, each simulating a source signal with a Gaussian temporal profile at the corresponding vertex on one of 11 surface meshes ranging from the white matter (deepest) to pial (most superficial) surface (see Supplementary Fig. 1 for simulated time-course). For each simulation, we ran source reconstruction (MSP, with the prior set to the simulated cortical location) within a 50 ms central window 11 times, each time using a different surface mesh to construct the forward model, and compared the resulting model evidence, approximated by free energy. In the absence of noise (e.g. sensor noise, co-registration error etc), the model based on the surface corresponding to the depth of the simulated signal will have greater model evidence, or free energy (i.e. if a signal is simulated on the pial surface, the model based on the pial surface should have the greatest free energy). This set of simulations was repeated at each selected cortical location with different levels of simulated sensor-level SNR, varying from -∞ (pure noise) to 5 dB.

For each location, this yielded a matrix of model evidence (free energy) with a column for simulation at each cortical depth, and a row for each forward model. For each simulated cortical depth (matrix column), we transformed free energy values for each forward model by subtracting the lowest value, thus representing model evidence relative to the worst model. For each matrix, we identified the model with the peak free energy for each simulated cortical depth, and generated a probability density of these peaks for each SNR level. We also computed a measure of diagonal dominance, or the ratio of diagonal values to total values (perfect reconstruction would yield a purely diagonal matrix), and compared this to a null distribution obtained by shuffling the matrices. At low levels of SNR (< -50 dB), there was no pattern in the mean free energy (averaged over all simulated locations) over simulated cortical depths (Figure 2a). However, the probability density of peak model evidence showed a bias toward either the most superficial or the deepest surface model (Figure 2e). This likely reflects the greater separability of their lead fields at the sensor level: superficial and deep sources produce more distinct field patterns than intermediate depths, making them preferentially selected under noisy conditions. Consistent with this interpretation, lead-field dissimilarity exhibited a U-shaped profile across cortical depth, with the most superficial and deepest layers showing the greatest separability from the remaining layers (Supplementary Fig. 2). At -50 dB, a step change in the mean free energy was observed, with the greatest mean free energy for roughly the surface corresponding to the simulated depth (Figure 2b), and a diagonal dominance score greater than expected by chance (standardized difference = 3.22, *p* = 0.001; Supplementary Fig. 3a). A corresponding gradient in the probability density of the peak free energy model was observed, with a greater probability for superficial surface model peaks for superficial simulations, and for deep surface model peaks for deep simulations (Figure 2f). At higher levels of SNR (≥- 35 dB), both the mean free energy (Figure 2c,d) and the probability density (Figure 2g,h) exhibited significant diagonality (all standardized differences > 3.5, all *p* < 0.001; Supplementary Fig. 3a), indicating near perfect inference of the depth of the simulated signal.

**Figure 2.**
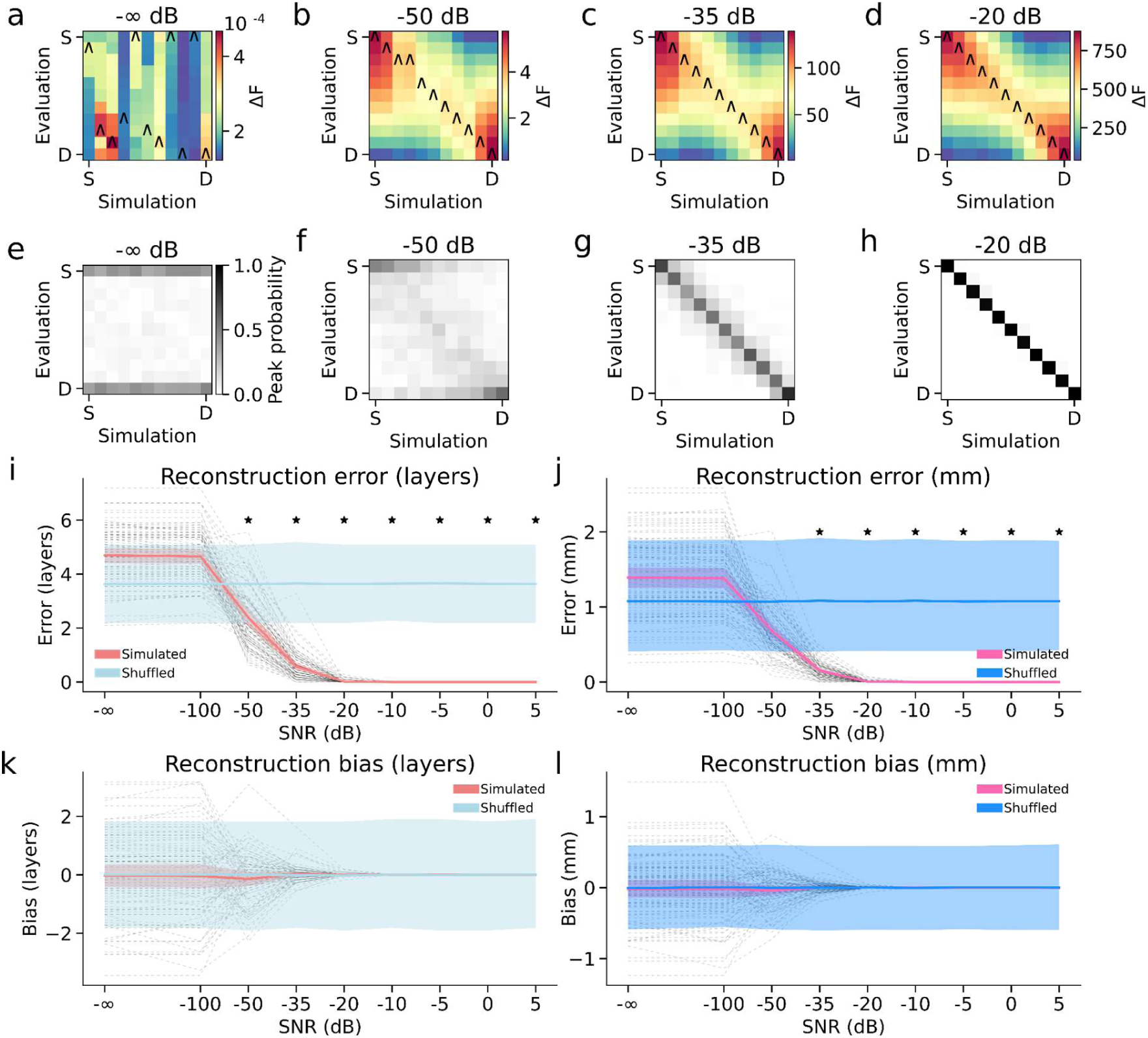
Laminar source-reconstruction accuracy as a function of sensor-level SNR. **a-d** Mean model evidence (free energy, ΔF) matrices (relative to the worst model; S = superficial surface layer, D = deep surface layer) for simulated signals placed on 11 cortical surfaces at four SNR levels (-∞ dB, - 50 dB, -35 dB, -20 dB). Accurate laminar inference begins to emerge around -35 dB, with near-perfect identification at -20 dB. Caret markers denote the forward model with peak free energy for each simulated depth. **e-h** Probability density of each forward model being the maximum-evidence solution, aggregated across cortical locations: a diagonal pattern appears from -35 dB, indicating accurate depth localization. **i, j** Laminar reconstruction error (in layers and mm) falls below chance at -35 dB and improves at higher SNR; shuffled controls show ∼1 mm error due to chance matches between simulated and evaluated layers. **k, l** Laminar reconstruction bias (in surface layers and millimeters), again comparing simulated and shuffled data. Shaded bands represent 95% confidence intervals, and asterisks mark SNR levels at which observed error or bias differed significantly from chance (*p* < 0.05).

At each cortical location, we computed bias as the mean distance, over the 11 simulations, between the simulated source and the surface model with the highest free energy (in terms of surfaces and millimeters), and error as the mean absolute distance between the simulated surface and the best surface models (also in surfaces and millimeters). For each metric, we compared the observed values across simulated SNR levels with the null distribution obtained by shuffling the surface model free energy for each simulated depth. At low SNR levels (−50 dB and below), both error and bias were at chance levels (*p* > 0.05 and *p* > 0.8, respectively). A significant reduction in error emerged at -35 dB (standardized difference = -4.17, *p* < 0.001), with further minimal error at -20 dB and above (all standardized differences > 4.8, all *p* < 0.001; Figure 2i,j). Bias was not significant at any SNR level (all *p* > 0.8), though extreme deep or superficial biases were observed at some locations below -50 dB (Figure 2k,l). These results were nearly identical when error and bias were computed in millimeters rather than surface layers (Figure 2k,l). Consistent with previous simulation results^20^, inference was biased at low SNR levels when using the empirical Bayesian beamformer algorithm (EBB; Supplementary Fig. 4)^45^ rather than MSP with a spatial prior, as used in this study. We also examined error separately for each simulated surface layer and found no differences between layers (Supplementary Fig. 5a). Laminar inference at a coarse scale (deep versus superficial) is therefore possible in principle at -50 to -35 dB SNR, with fine-scale inference possible at -20 dB and above.

Previous work suggests that accurate laminar inference is feasible with up to 2 mm of co-registration error^19,20^. To evaluate whether this holds in our multilayer framework, we ran another set of similar simulations at the same cortical locations, this time fixing the SNR at 0 dB, and varying the simulated co-registration error from 0 to 5 mm. At levels of co-registration less than or equal to 2 mm, the surface model with the highest mean free energy corresponded to the simulated cortical depth (Figure 3a-c), and the peak free energy model probability density exhibited strong diagonality (all standardized differences > 4, all *p* < 0.001: Figure 3e-f; Supplementary Fig. 3b), indicating accurate inference across the range of simulated depths. At higher levels of co-registration error (3-5 mm), the mean free energy still exhibited significant diagonality (all standardized differences > 3.9, all *p* < 0.001; Figure 3d; Supplementary Fig. 3b), but the probability density only indicated a gradient in free energy peaks in the deepest and most superficial surfaces (Figure 3h).

**Figure 3.**
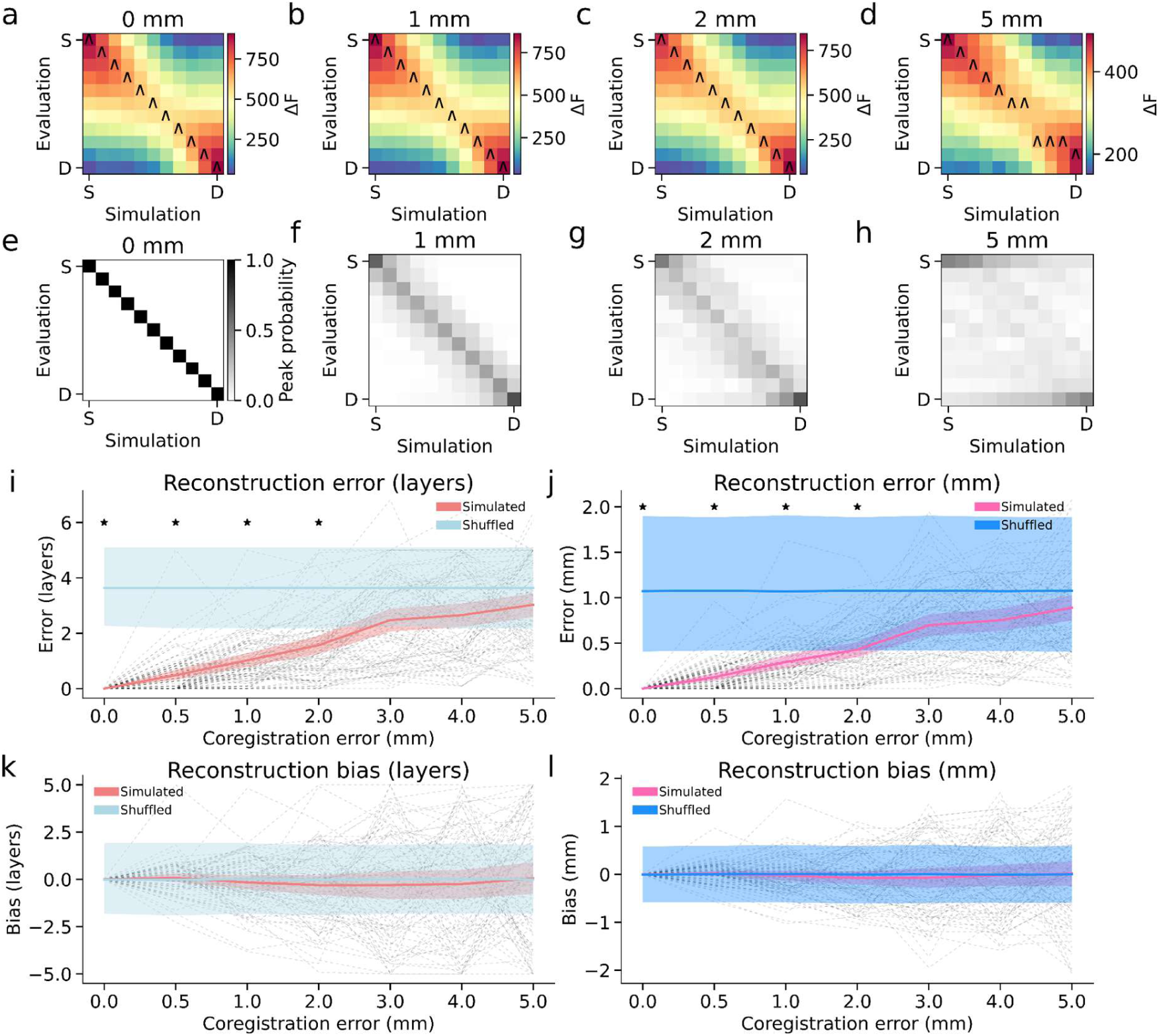
Laminar source-reconstruction accuracy as a function of co-registration error at fixed 0 dB SNR. **a-d** Mean free energy (ΔF) matrices (relative to the worst model; S = superficial surface layer, D = deep surface layer) for simulated signals placed on 11 cortical surfaces at 0, 1, 2, and 5 mm co-registration error. Carets mark the best-fitting forward model for each simulated depth. **e-h** Probability density of each surface model being the maximum-evidence solution, aggregated across cortical locations. **i, j** Laminar reconstruction error (in surface layers and millimeters) for simulated versus shuffled data at varying registration errors. **k, l** Corresponding bias (in surface layers and millimeters). Shaded bands denote 95% confidence intervals, and asterisks mark significance at p < 0.05.

We computed bias and error for the co-registration error simulations using the same approach as in the SNR simulations and compared the observed values against the null distribution obtained by shuffling. Error was lowest at 0 mm (standardized difference = -4.84, *p* < 0.001) and progressively increased with greater co-registration error, though it remained significantly lower than chance levels up to 2 mm (standardized difference = -2.77, *p* = 0.002; Figure 3i,j). At 3 mm and above, error was at chance level (all *p* > 0.06). There was no significant bias at any level of co-registration error (all *p* > 0.7). These results were nearly identical when error and bias were computed in millimeters rather than surface layers (Figure 3k,l). When split by simulated surface layer, the accuracy of laminar inference for simulated deep sources was less affected by co-registration error than superficial sources (Supplementary Fig. 5b). Accurate laminar inference therefore requires a co-registration error of 2 mm or less, though a coarse deep-versus-superficial distinction may be possible in some locations at errors up to 5 mm, provided there is sufficient SNR.

Given that cortical laminae vary in thickness both within and across regions^38^, we assessed laminar reconstruction accuracy in terms of cytoarchitectonic laminae rather than fixed-depth surfaces. Specifically, surface layers that mapped to the same lamina were treated as equivalent (e.g., errors between adjacent surface layers were not considered errors if both corresponded to lamina VI). To achieve this, we re-analyzed the effects of SNR and co-registration error using the BigBrain atlas^40^, which we mapped to the subject’s anatomy via FreeSurfer’s surface-based registration through fsaverage space^41^. This allowed us to estimate proportional laminar thickness at each simulated cortical location (Figure 4a-f), assign surface layers to their corresponding laminae, and evaluate reconstruction accuracy in terms of cytoarchitectonic rather than geometric depth.

**Figure 4.**
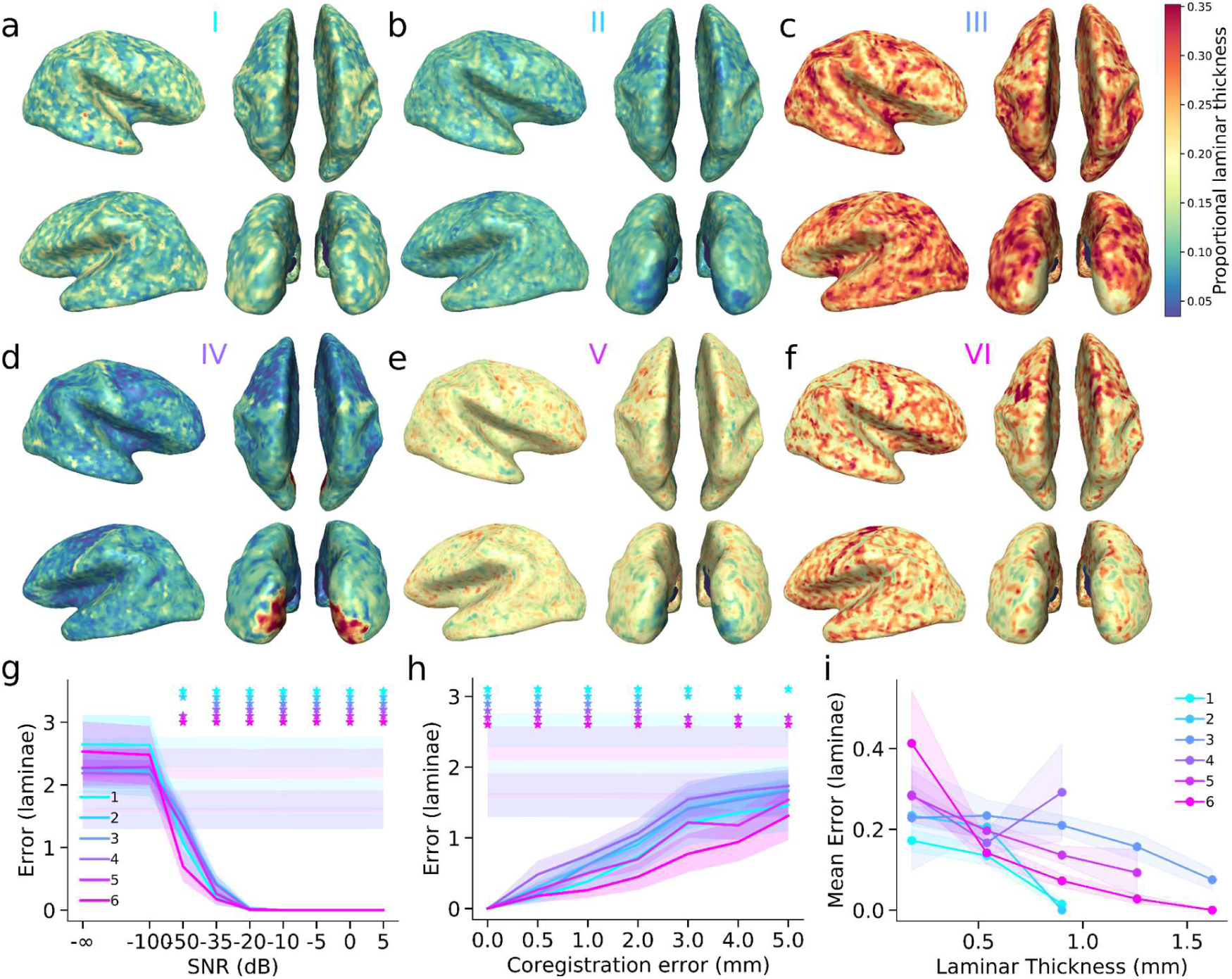
Laminar source-reconstruction accuracy as a function of SNR, co-registration error, and cortical thickness. **a-f** Cortical surfaces showing proportional laminar thickness across the brain, derived from the BigBrain atlas and mapped onto the surface models (representing laminae I-VI respectively) used in the simulations. These maps illustrate the variability of laminar thickness across cortical regions, which influences inference accuracy. Proportional laminar boundaries were subsequently scaled by local cortical thickness to compute laminar depths in millimeters for reconstruction-error analyses. **g** Mean laminar reconstruction error (in laminae) across SNR levels, computed as the absolute difference between the simulated and inferred lamina. Lines indicate the mean error for each lamina, with shaded bands representing 95% confidence intervals. Lighter shaded bands represent shuffled null distributions. Asterisks mark SNR levels where reconstruction error was significantly lower than chance. **h** Mean laminar reconstruction error as a function of co-registration error. **i** Relationship between laminar thickness and reconstruction error. Each line represents a different lamina (I-VI), showing how increased laminar thickness is associated with reduced mean reconstruction error (the shaded regions denote SEM).

We re-analyzed both SNR and co-registration error simulations under this framework. Error was computed as the absolute difference between the simulated and inferred lamina, and significance was assessed using permutation tests (*N* = 10,000) against a shuffled null distribution. At very low SNR levels (−100 dB and below), reconstruction error was indistinguishable from chance for all laminae (all *p* > 0.8). At -50 dB, inference remained at chance for middle laminae (laminae III and IV; both *p* > 0.07) but was significantly better than chance for superficial and deep laminae (all *p* < 0.001). By -35 dB, reconstruction accuracy improved across all laminae (all *p* < 0.001), with standardized differences ranging from -11.36 to -23.15 (Figure 4g). The most pronounced improvements were observed at -20 dB and above, where errors reached minimal levels (all standardized differences < -14.57, all *p* < 0.001). These results confirm that true laminar inference is possible at sufficiently high SNR, even when accounting for regional variability in cortical laminae thickness.

Co-registration errors had a similar impact, with small misalignment (≤ 2 mm) having minimal effects on reconstruction accuracy across all laminae (all *p* < 0.001), while increasing error progressively degraded performance. At 2 mm, reconstruction accuracy remained high (all standardized differences < -4.45, all *p* < 0.001), but at 3 and 4 mm, performance declined sharply, particularly for middle laminae (laminae III and IV; both *p* > 0.4; Figure 4h). At 5 mm, only the most superficial (lamina I) and deepest laminae (laminae V and VI) remained significantly different from chance (lamina I: standardized difference = -8.51, *p* < 0.001; laminar V: standardized difference = -2.10, *p* = 0.018; lamina VI: standardized difference = -8.45, *p* < 0.001). These findings suggest that fine-scale laminar inference is robust to small co-registration errors but becomes unreliable at misalignment exceeding ∼3 mm, particularly for middle-layer sources. At errors of 5 mm, only deep and superficial sources retain some discriminability, indicating that accurate reconstruction at this scale requires co-registration precision within at least approximately 2 mm.

Finally, we assessed how laminar reconstruction accuracy depends on laminar thickness, focusing on the results of simulations with SNR of at least -20 dB and co-registration errors less than or equal to 2 mm. Across simulated locations, thicker laminae exhibited systematically lower reconstruction error (Figure 4i), reflecting a greater spatial margin for accurate depth localization. This effect was most pronounced in deep and superficial laminae, where increased cortical thickness improved inference reliability. Middle laminae, in contrast, showed greater variability in error, consistent with the higher misclassification rates observed at moderate SNR and co-registration error levels. Cortical laminae thickness therefore impacts laminar reconstruction accuracy, with thicker laminae providing a greater margin for accurate depth localization. The greater variability observed for middle laminae likely reflects both their position with the cortical depth axis, where errors can occur in either direction, and regional variation in laminar thickness. Reconstruction accuracy is therefore likely shaped by both geometric constraints and regional differences in laminar architecture.

### Limitations of laminar source separation

Having established that a single source can be accurately localized in depth given sufficient SNR and co-registration accuracy, we next examined whether two simultaneous sources of equal strength at different layers could be differentiated. To test this, we simulated two sources at each of the 100 cortical locations: one fixed in the middle layer and the other varying from the most superficial to the deepest layer surface (SNR = -20 dB, co-registration error = 0 mm). Across all simulations, rather than producing two distinct peaks in relative model evidence, a single peak consistently emerged at an intermediate depth between the two sources, both in mean free energy (Figure 5a) and in the peak free energy model probability density (Figure 5b). This suggests that the presence of the middle-layer source systematically biased inference toward the middle-layer model compared to single-source simulations (Figure 5e,f). However, further analysis of the free energy distribution across models (Figure 5c) and across simulations (Figure 5d) suggests that distinct contributions from both sources may still be present. Notably, model evidence for the middle-layer surface was higher relative to single-source simulations (Figure 5g,h), indicating that while the current approach does not fully resolve two simultaneous sources, further refinement of the method may improve discriminability.

**Figure 5.**
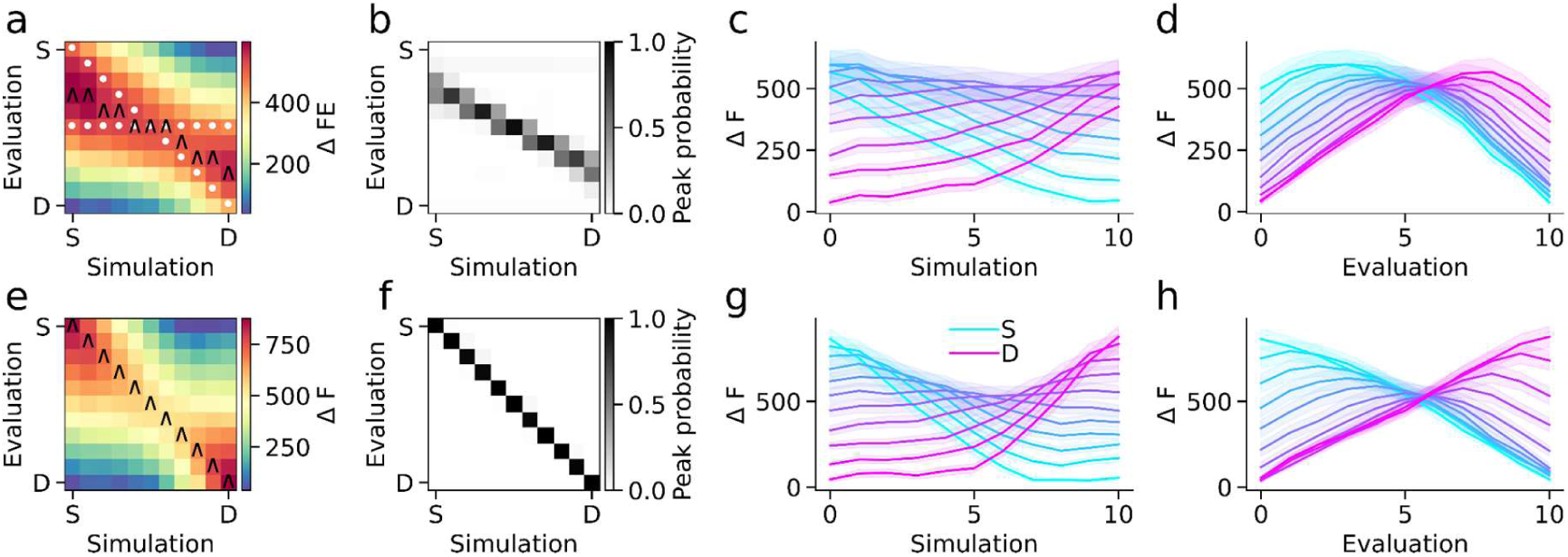
Two-source simulations at -20 dB SNR highlight the difficulty of separating simultaneous sources at different depths. **a,b** When one source was fixed at the middle layer and a second source was varied from superficial to deep, the mean free-energy (ΔF) matrices and peak-probability plots showed a single intermediate peak, indicating bias toward the middle-layer model (S = superficial surface layer, D = deep surface layer). **c,d** Despite this bias, plots of mean ΔF across simulated layers (c) and across forward-model evaluations (d) reveal that deeper and more superficial contributions still influence the model evidence (shaded regions denote SEM). **e-h** In contrast, single-source simulations at the same SNR showed stronger diagonality in ΔF (e,f) and clearer separation in layer-by-layer analyses (g,h), suggesting that additional refinements are needed to fully disentangle two concurrent laminar sources.

### Cortical column orientation and laminar inference

We assessed the influence of dipole orientation estimation on laminar source reconstruction by evaluating four methods^28^: surface normals from the original high-resolution mesh (OSN), surface normals from the downsampled mesh (DSN)^32^, cortical patch statistics (CPS)^33^, and link vectors (LV) computed between corresponding vertices on the pial and white matter surfaces^27^. For each method except link vectors, we included variants with orientations fixed across layers (“fixed”) and orientations independently computed per layer (link vectors have the same orientation at corresponding vertices across surfaces by definition). This resulted in seven variations in dipole orientation definition. Figure 6a illustrates these methods applied across cortical depth at representative vertices.

**Figure 6.**
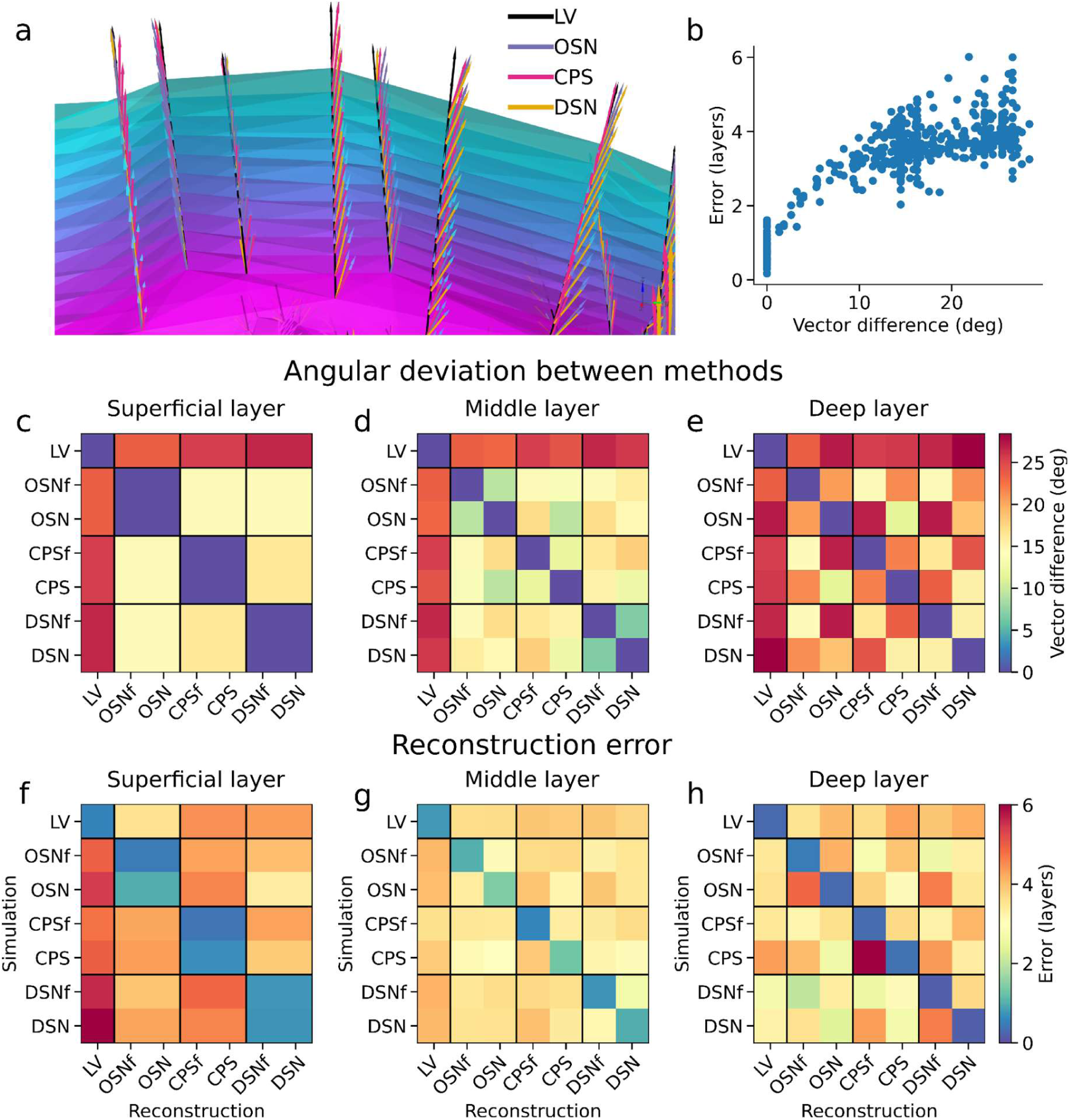
Impact of dipole orientation estimation on laminar source reconstruction accuracy. **a** Example dipole orientation vectors at a subset of cortical locations across layers for four methods: link vectors (LV), original surface normals (OSN), cortical patch statistics (CPS), and downsampled surface normals (DSN). **b** Relationship between orientation mismatch (angular difference in degrees) and laminar reconstruction error, over all simulation-reconstruction method pairs and cortical depths. **c-e** Angular deviation (median across vertices) between orientation estimation methods at three cortical depths: superficial (c, surface layer 1), middle (d, surface layer 6), and deep (e, surface layer 11). **f-h** Corresponding reconstruction error (in layers) for each simulation-reconstruction method pair at the same depths, averaged across 100 cortical locations. Errors are lowest along the diagonal, where simulation and reconstruction methods match, and largest when methods differ substantially (LV = link vectors; OSN = original surface normals; CPS = cortical patch statistics; DSN = downsampled surface normals. The suffix “f” indicates dipole orientations were fixed across layers).

To quantify differences between orientation vectors, we computed the angular deviation between each pair of methods at each cortical depth. These deviations were highest between LV and surface-normal-based approaches (Figure 6c-e). The fixed variants of the surface-normal-based approaches used the surface normals computed from the pial surface, and therefore the angular deviation between fixed and non-fixed variants of the same family increased with cortical depth (Figure 6c-e). We then tested how these orientation discrepancies affected reconstruction performance by simulating laminar sources using each method and reconstructing them with each of the others. Reconstruction error, defined as the deviation between the inferred and ground truth layer, was lowest when the same method was used for both simulation and reconstruction (Figure 6f-h), consistent with prior findings^28^. Errors were highest when simulation and reconstruction orientations were mismatched, particularly across different estimation families (e.g., LV vs DSN).

Across cortical depths, we observed a strong positive and nonlinear relationship between angular discrepancies in dipole orientation and errors in laminar source reconstruction (Figure 6b), indicating that even modest orientation misalignment can degrade depth precision. Importantly, we found no consistent advantage for fixed-orientation strategies over those allowing depth-varying orientations, suggesting that anatomical consistency across layers is less critical than aligning dipole orientations with the true cortical column geometry. These results underscore the importance of accurately estimating dipole orientation when performing laminar inference in real data, and caution against using anatomically implausible orientation models.

### Robustness to noise, model mismatch, and concurrent sources

To assess the extent to which the simulation results generalize beyond the idealized conditions of the primary analyses, we performed a series of robustness analyses that relaxed key assumptions regarding noise structure, forward-model accuracy, and source complexity (Supplementary Fig. 6-13). First, we repeated the SNR and co-registration simulations using temporally autocorrelated pink noise rather than white noise. Reconstruction accuracy, bias, and free-energy structure were largely unchanged, indicating that the principal findings do not depend on the assumption of spectrally flat sensor noise Supplementary Fig. 6 and 10). Second, we evaluated robustness to concurrent activity by introducing three additional interfering sources at random cortical locations. Each interfering source had an independent band-limited temporal profile and an amplitude equal to 10% of the primary source. Although reconstruction accuracy was modestly reduced, reliable laminar inference still emerged between approximately -35 and -20 dB SNR and deteriorated beyond approximately 2-3 mm co-registration error, closely matching the results of the primary simulations (Supplementary Fig. 7 and 11). Third, we assessed robustness to source-space mismatch using an alternative generative model in which simulated activity was generated on cortical surfaces containing 32,399 vertices per layer, whereas inversion was performed using the source model employed throughout the manuscript (29,130 vertices per layer). This mismatch resulted in modest reductions in reconstruction accuracy but preserved the overall dependence on SNR and co-registration precision (Supplementary Fig. 8 and 12). Finally, because laminar inference relies on subtle differences in lead fields across cortical depth, we evaluated the effect of forward-model mismatch by generating data using a BEM forward model while reconstructing sources using the Nolte single-shell model used throughout the main analyses. This manipulation produced the largest degradation in performance among the robustness analyses, reducing the range of SNRs and co-registration errors over which accurate laminar inference could be achieved. Nevertheless, the overall structure of the results remained similar to the primary simulations, with reconstruction accuracy improving monotonically with increasing SNR and decreasing co-registration error (Supplementary Fig. 9 and 13). To summarize these analyses, Figure 7 compares mean reconstruction error across all robustness conditions. Although performance was reduced under model mismatch and BEM-generated data, the qualitative conclusions regarding the requirements for reliable laminar inference remained unchanged. Collectively, these results indicate that the principal findings are robust to realistic deviations from the assumptions of the primary simulations, while highlighting the importance of accurate forward modeling for depth-resolved source reconstruction.

**Figure 7.**
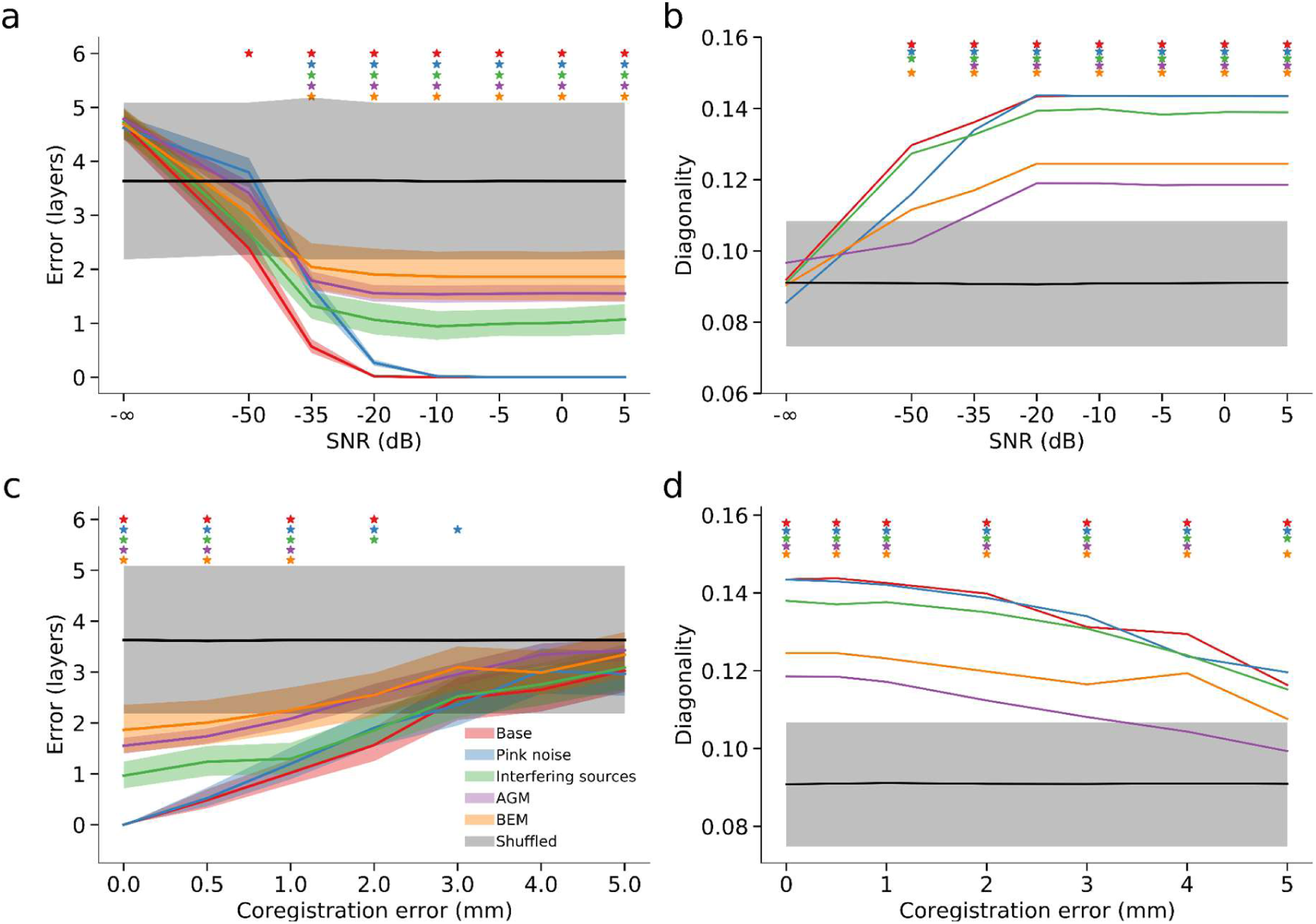
Robustness of laminar source reconstruction to violations of simulation assumptions. Comparison of reconstruction performance across the primary simulations (base) and four robustness analyses: pink noise, interfering sources, alternative generative model (AGM), and boundary element model (BEM) generative model. The AGM simulations used mismatched cortical source-space discretizations for simulation and inversion, whereas the BEM simulations used a BEM forward model for data generation and a Nolte single-shell model for source reconstruction. **a** Mean laminar reconstruction error (layers) for simulated versus shuffled data a function of SNR. **b** Diagonal dominance of the mean model-evidence matrices (trace/sum) for simulated versus shuffled data as a function of SNR. **c** Mean laminar reconstruction error (layers) for simulated versus shuffled data as a function of co-registration error. **d** Diagonal dominance of the mean model-evidence matrices (trace/sum) for simulated versus shuffled data as a function of co-registration error. Shaded regions indicate ± SEM across simulated cortical locations. Colored asterisks denote simulation conditions in which performance differed significantly from chance (p < 0.05). Across all robustness analyses, the qualitative dependence of laminar inference on SNR and co-registration precision was preserved. Pink-noise simulations produced results nearly identical to the primary simulations, whereas interfering sources caused only modest reductions in performance. The largest decreases in reconstruction accuracy and matrix diagonal dominance were observed for simulations incorporating forward-model mismatch (AGM and BEM).

### Anatomical features determine laminar inference accuracy

The previous analyses demonstrate that laminar inference accuracy critically depends on signal quality, co-registration precision, and accurate estimates of cortical column orientation. However, even with optimal signal-to-noise ratio (≥ -20 dB) and zero co-registration error, reconstruction accuracy is not uniform across the cortex^20^. This variability suggests that local anatomical properties may impose inherent constraints on laminar reconstruction. To evaluate how anatomical variability influences laminar source reconstruction, we simulated, one at a time, sources at every vertex of a multilayer cortical mesh (29,130 locations × 11 layers) under these idealized conditions. Layer depth was inferred using model evidence across all cortical layers. Reconstruction error, defined as the absolute difference between the simulated and inferred depth, was zero for 24,699 out of 29,130 vertices (84.79%), indicating that laminar inference is feasible across most of the cortex under these conditions. However, reconstruction accuracy varied regionally, with errors concentrated in anatomically constrained areas (Figure 8a).

**Figure 8.**
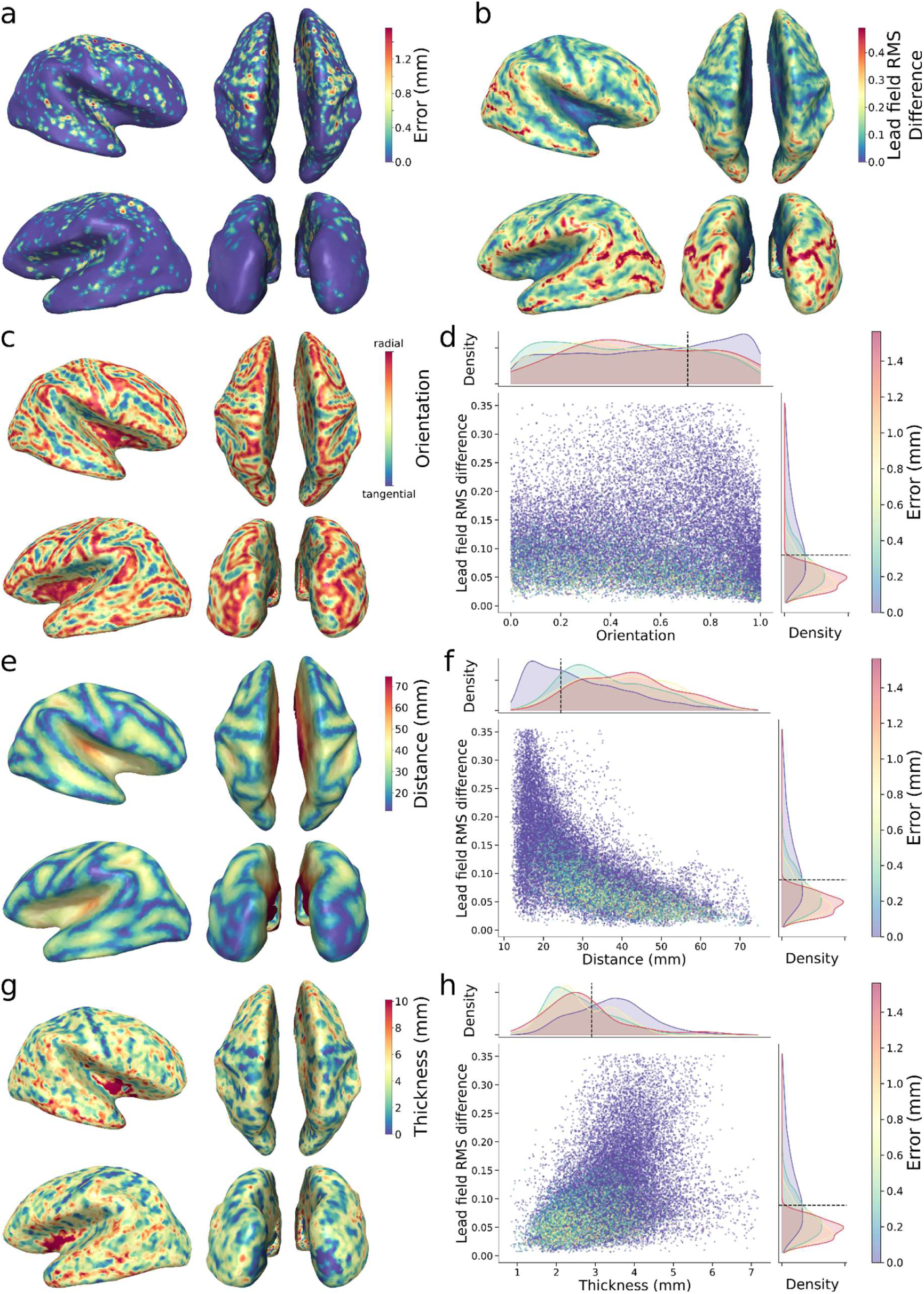
Anatomical factors influencing laminar reconstruction error. **a-b** Inflated-surface maps of laminar reconstruction error (a) and lead-field RMS difference (deep vs. superficial; b) **c-h** Inflated-surface maps and joint density and marginal histograms for each anatomical metric versus lead-field RMS difference: and dipole orientation (c,d), scalp-to-cortex distance (e,f), and cortical thickness (g,h). Joint density plots are color-coded by laminar reconstruction error (mm), and the marginal histograms group vertices into zero-error and three equally spaced bins spanning the range of non-zero errors. Lower lead-field differences, tangential orientations, greater distance to the scalp, and thinner cortex are associated with increased laminar-reconstruction error. Optimal thresholds in panels d, f, and h are indicated by black dashed lines and were defined using receiver operating characteristic (ROC) analysis (maximizing Youden’s J statistic) to discriminate zero versus non-zero reconstruction error.

To understand the anatomical basis for this regional variation, we examined four candidate factors: lead field separation between deep and superficial layers (Figure 8b), dipole orientation relative to the scalp (Figure 8c), scalp-to-cortex distance (Figure 8e), and cortical thickness (Figure 8g). Although none of these features perfectly separated vertices with zero error from those with nonzero error, all were significantly associated with reconstruction error. Specifically, vertices with low laminar discriminability exhibited smaller lead field differences between depths, more tangential orientations, greater distances from sensors, and thinner cortices.

Joint density plots and marginal histograms revealed that these relationships were nonlinear (Figure 8d,f,h). Among all features, the dissimilarity between deep and superficial lead fields emerged as the strongest predictor of error (optimal threshold = 0.09 a.u.; AUC = 0.756), suggesting that the ability to distinguish layer-specific contributions at the sensor level places a fundamental constraint on laminar inference. Cortical thickness (optimal threshold = 2.91 mm; AUC = 0.749) and scalp distance (optimal threshold = 24.43 mm; AUC = 0.689) also contributed, consistent with prior work on MEG sensitivity profiles, whereas dipole orientation had a weaker but still detectable effect (optimal threshold = 0.71 a.u.; AUC = 0.595). These findings suggest that even under favorable conditions, anatomical constraints shape the spatial distribution of inference precision.

To determine whether combinations of anatomical features better predict laminar inference accuracy than single features, we performed a multivariate logistic regression analysis. The logistic regression model significantly discriminated between error-free and erroneous reconstructions (mean cross-validated AUC = 0.810 ± 0.013), with the largest positive coefficient assigned to lead field difference (*β* = 10.64), followed by orientation relative to the scalp (*β* = 1.01) and cortical thickness (*β* = 0.76); scalp-to-cortex distance had minimal influence (*β* = -0.02). Adding pairwise interaction terms provided only a negligible improvement (mean cross-validated AUC = 0.812 ± 0.015). A random forest classifier corroborated these findings, highlighting the lead field difference as the strongest predictor (importance = 32%), followed by thickness (27%), distance (22%), and orientation (19%), though it did not enhance prediction accuracy beyond logistic regression (mean cross-validated AUC = 0.804 ± 0.013). Together, these results indicate that anatomical constraints on laminar inference accuracy primarily arise from additive contributions of distinct features rather than from complex interactions.

To determine how anatomical features influence laminar inference accuracy, we performed a mediation analysis with lead-field difference as a mediator. Cortical thickness showed a strong indirect effect via lead-field separability (indirect effect = 0.49, 95% CI [0.46, 0.52]), alongside a residual direct effect (*c′* = 0.67), indicating partial mediation. Scalp-to-cortex distance exhibited a smaller but significant indirect effect (indirect effect = -0.08, 95% CI [-0.08, -0.07]) with negligible direct effect (*c′* ≈ 0), consistent with near-complete mediation. In contrast, dipole orientation showed no significant indirect effect (indirect effect = 0.05, 95% CI [-0.01, 0.11]) but a strong direct effect (*c′* = 1.51), suggesting an additional contribution to signal detectability independent of depth separability. Cortical thickness and distance therefore influence laminar inference primarily through their effect on lead-field separability, whereas dipole orientation exerts a largely independent effect, reinforcing the central role of lead-field separability as the primary determinant of laminar inference accuracy.

### Empirical laminar inference in visual and sensorimotor cortex

Having established through simulations the conditions under which laminar inference is reliable, we next applied multilayer hpMEG source reconstruction to three empirical datasets: a visuospatial attention task, a visually cued button-press task, and a self-paced button-press task, using these insights to guide analysis choices. These datasets were chosen because they elicit well-characterized visual and sensorimotor ERFs with established cortical generators and expected laminar activation motifs.

Prior to laminar analysis, we quantified sensor-level signal-to-noise ratio (SNR) within the component-specific analysis windows and examined its relationship to trial count across participants. Across datasets, both trial count and sensor-level SNR varied substantially, with no simple linear relationship between the two (Supplementary Fig. 14). This likely reflects differences in component amplitude, noise structure, and preprocessing efficacy across paradigms and participants. Because laminar inference was performed on trial-averaged ERFs rather than single-trial activity, reconstruction performance depends primarily on the SNR of the averaged response rather than on trial count per se. Participants whose averaged responses exceeded an SNR threshold of -15 dB were retained for laminar analysis.

In the visuospatial attention task, participants were cued to attend to one visual hemifield, and for each trial we analyzed the VEF separately in the hemisphere contralateral versus ipsilateral to the attended stimulus location^46^. For each condition, the peak of the early visually evoked field (VEF) component was localized to primary visual cortex (V1), and sliding-window laminar inference was performed at a single anatomically and functionally optimal vertex. Vertex selection was based on independent functional localization and anatomical constraints identified in simulations (e.g., cortical thickness, dipole orientation, and lead field separability), which determine whether laminar inference is feasible at a given location (see Methods). This approach prioritizes regions where the inverse problem is sufficiently well-conditioned for laminar discrimination, rather than applying laminar inference uniformly across anatomically defined regions where identifiability may be poor.

The resulting laminar dynamics closely matched established feedforward models of early visual processing. At the peak of the VEF (80-100 ms; *M* = 90.6 ms), model evidence was dominated by superficial laminae (I-III) together with lamina IV (Figure 9; ipsilateral condition in Supplementary Fig. 15), consistent with thalamocortical input to lamina IV and subsequent activation of supragranular pyramidal populations^47,48^. To assess consistency across participants, we performed family-wise random-effects Bayesian model selection^42–44^, grouping laminae into supragranular (I-III), granular (IV), and infragranular (V-VI) families. Protected exceedance probabilities (PEPs) were computed at each time point to quantify the likelihood that one laminar family dominated over the others while accounting for chance-level equivalence^43,44^. This analysis confirmed a canonical feedforward sequence: early dominance of the granular family (around 60 ms), consistent with initial thalamic input to V1; and a subsequent supragranular dominance during the VEF peak (79-104 ms)^5,49–51^. An equivalent pattern was observed in the ipsilateral hemisphere (Supplementary Fig. 15). These results demonstrate that multilayer hpMEG recovers the expected laminar sequence of early visual processing in human V1, validating the empirical applicability of the multilayer framework and its ability to resolve feedforward laminar dynamics non-invasively.

**Figure 9.**
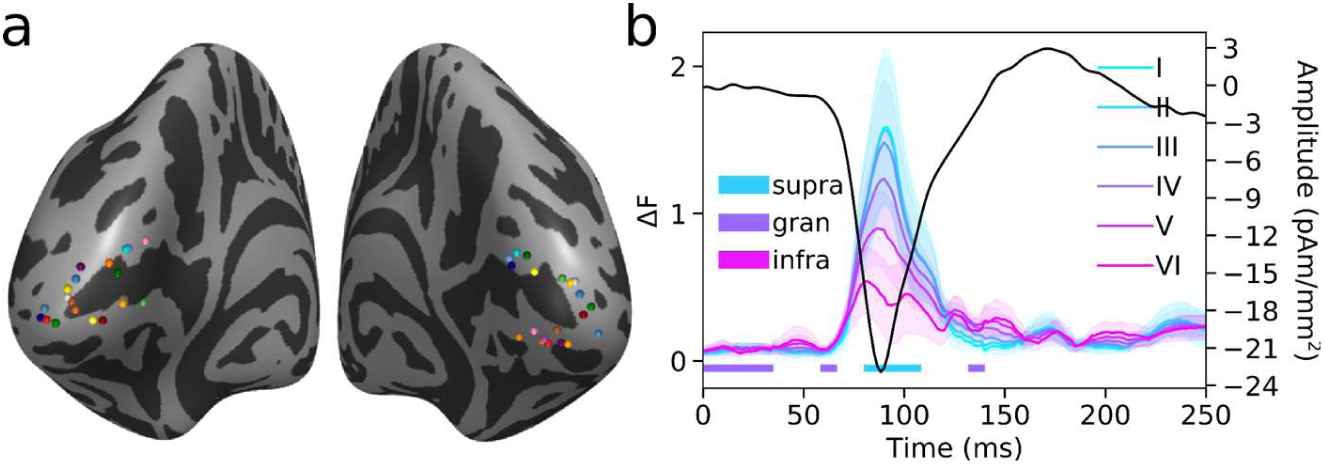
Feedforward laminar activation in primary visual cortex during visuospatial attention. **a** Locations of the selected VEF source vertices for each participant, projected onto an inflated fsaverage cortical surface; each colored sphere denotes one subject. **b** Sliding-window laminar inference applied to visually evoked fields (VEFs) localized to primary visual cortex (V1) during a visuospatial attention task, analyzing responses in the hemisphere contralateral to the attended stimulus location (see Supplementary Fig. 15 for the ipsilateral condition). Colored traces show lamina-specific relative model evidence (ΔF, relative to the worst model at each time point) averaged across participants and shaded regions denote SEM; the dark trace shows the corresponding middle-layer source time series. Horizontal color bars indicate, at each time point, the laminar family with the highest protected exceedance probability (PEP), grouping layers into supragranular (laminae I-III), granular (lamina IV), and infragranular (laminae V-VI) families. The VEF exhibits a canonical feedforward laminar sequence, with early dominance of the granular family consistent with thalamic input, followed by supragranular dominance at the VEF peak.

We next applied multilayer hpMEG source reconstruction to motor ERFs using two independent datasets: one involving visually cued button presses and another involving self-paced button presses. In each dataset, we localized the motor field (MF) component to the hand area of primary motor cortex (M1) and the motor-evoked field I (MEFI) component to the corresponding region of primary somatosensory cortex (S1), and performed sliding-window laminar inference at each site. Qualitative inspection revealed broadly similar laminar dynamics across the two datasets for both components (Supplementary Fig. 16), motivating their combination to enable group-level Bayesian model selection, which was not feasible within either dataset alone due to limited sample size.

Using the combined dataset, we performed family-wise Bayesian model selection for the MF in M1 and the MEFI in S1 (Figure 10). In M1, laminar inference revealed a reproducible sequence dominated by infragranular activity (laminae V-VI) beginning approximately 150 ms before movement onset, consistent with the contribution of deep-laminae pyramidal neurons projecting to the corticospinal tract to the generation of the MF component^52–54^. This deep-laminae dominance recurred in two additional transients, coinciding with the button press and shortly thereafter. A later emergence of supragranular activity (laminae I-III), approximately 300-350 ms after the button press, followed the deep-laminae transients. This temporal pattern suggests successive phases of motor preparation, corticospinal output, and post-movement reactivation, with the delayed superficial response potentially reflecting feedback or efference-copy-related processing within local cortical circuits.

**Figure 10.**
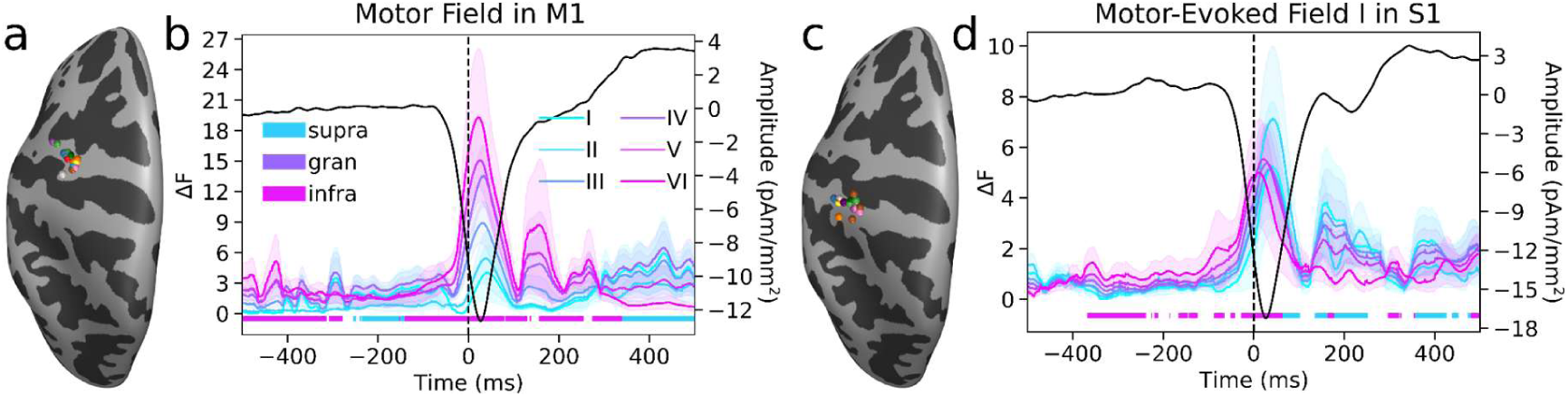
Laminar activation sequences of motor and somatosensory ERFs during movement. **a** Locations of the selected motor field (MF) source vertices for each participant in the hand area of primary motor cortex (M1), projected onto an inflated fsaverage cortical surface; each colored sphere denotes one subject. **b** Sliding-window laminar inference for the MF using the combined dataset of visually cued and self-paced button presses. **c** Locations of the selected motor-evoked field I (MEFI) source vertices for each participant in primary somatosensory cortex (S1). **d** Sliding-window laminar inference for the MEFI using the combined dataset. In b and d, colored traces show lamina-specific relative model evidence (ΔF, relative to the worst model at each time point) averaged across participants and shaded regions denote SEM; dark traces show the corresponding middle-layer source time series. Horizontal color bars indicate, at each time point, the laminar family with the highest protected exceedance probability (PEP), grouping layers into supragranular (laminae I-III), granular (lamina IV), and infragranular (laminae V-VI) families. MF responses in M1 are characterized by dominant infragranular activity preceding, during, and shortly after movement, followed by delayed supragranular activation, whereas MEFI responses in S1 show predominant infra- and then supra-granular activity at movement onset and subsequent deep and superficial transients.

Laminar inference of the MEFI in S1 showed a distinct profile, characterized by predominant infa- and then supra-granular activity at movement onset and additional deep and superficial transients at later latencies. This pattern is consistent with prior work identifying the MEFI as a short-latency (∼40 ms) response to proprioceptive and cutaneous reafference in the postcentral gyrus^52,55^. The qualitative correspondence between laminar dynamics observed in the two original datasets (Supplementary Fig. 16), together with their convergence in the combined analysis, supports the use of multilayer hpMEG to recover established sensorimotor activation motifs and extend them into the laminar domain.

## DISCUSSION

Building upon previous models of laminar inference in high-precision MEG (hpMEG) that compared deep versus superficial sources, we introduce a multilayer source reconstruction framework that aims to enable depth-resolved inference across the cortical column. Simulations establish feasibility limits for laminar inference, anatomical constraints guide vertex selection, and Bayesian model comparison provides statistical inference over cortical depth. In the simulations, this precision required a sufficiently high signal-to-noise ratio (approximately -35 dB for deep-versus-superficial distinctions and -20 dB for full laminar inference) and minimal co-registration error (< 2mm), conditions that can be met in practice through the use of individualized head-casts and optimized recording protocols^21,22^. While coarse deep-versus-superficial distinctions remain feasible at lower SNRs with two-layer source models, resolving finer laminar structure requires sufficiently high SNR, precise co-registration, and accurate forward modeling, and should therefore be interpreted as conditional on these factors. Importantly, supplementary robustness simulations demonstrated that these conclusions were largely preserved under more realistic conditions, including temporally autocorrelated noise, concurrent interfering sources, source-space mismatch, and forward-model mismatch, although the latter reduced the range of SNRs and co-registration errors over which accurate laminar inference could be achieved. Crucially, we confirm that correctly estimating cortical column orientation is essential for accurate laminar inference, as errors in dipole orientation estimation systematically degrade localization accuracy. Corroborating previously established results^20^, we show that anatomical features of the cortex such as thickness, dipole orientation, scalp-to-cortex distance, and lead field separation across depths nonlinearly shape the spatial distribution of inference accuracy, with lead field separability emerging as the strongest predictor.

Beyond simulations, we demonstrate the empirical plausibility of multilayer hpMEG by examining whether it recovers laminar activation patterns in human visual and sensorimotor cortex that are consistent with established physiological models. Notably, visual and motor ERFs exhibited complementary laminar sequences, with early granular and supragranular dominance in visual cortex and early infragranular dominance in motor cortex, reflecting their distinct circuit architectures and afferent/efferent nature, respectively. In primary visual cortex, visually evoked fields exhibited a feedforward laminar progression, with early granular dominance consistent with thalamocortical input, followed by supragranular activity at the response peak, with the same feedforward laminar sequence observed in both contralateral and ipsilateral hemispheres. In motor circuits, combining two independent datasets revealed consistent laminar trends across datasets: motor fields in primary motor cortex were dominated by infragranular activity beginning before movement onset and recurring around execution, consistent with corticospinal output, followed by delayed supragranular responses, while motor-evoked responses in primary somatosensory cortex showed predominant superficial activity at movement onset, consistent with somatosensory reafference. The convergence of these laminar patterns across cortical systems, task variants, and acquisition sites suggests that multilayer hpMEG can recover established physiological motifs at the level of laminar families under favorable conditions, providing evidence that non-invasive laminar inference can extend beyond deep-versus-superficial distinctions.

The empirical results demonstrate that multilayer hpMEG can recover physiologically plausible laminar activation sequences in both visual and sensorimotor cortices. At the same time, the simulations highlight important constraints on the interpretation of these findings. Laminar inference depends critically on the accuracy of the underlying forward model, is best interpreted as identifying the dominant laminar contribution within a localized cortical region and time window rather than fully resolving multiple simultaneous generators, and remains sensitive to methodological choices such as patch size, sliding-window duration, and preprocessing.

Accurate laminar inference ultimately depends on the fidelity of the forward model. A potential concern is that monotonic variations in peak model evidence across cortical depth could reflect depth-dependent biases in the forward model, signal-to-noise gradients, or regularization effects rather than true laminar specificity. However, our interpretation does not rely on static depth profiles at a single latency, but on the temporal evolution of laminar model evidence, which shows systematic shifts in dominant laminar contributions over time. Such dynamic transitions are not readily explained by fixed biases. Moreover, because the layer-specific source models are matched in complexity and orientation priors through vertex correspondence and link vector constraints, differences in model evidence primarily reflect depth-dependent differences in the forward model rather than differences in model dimensionality. Nonetheless, resolving activity from intermediate laminae remains challenging, particularly under lower SNR conditions. The simulations further demonstrated that lead field separability between superficial and deep cortical layers is the strongest anatomical predictor of laminar inference accuracy, emphasizing the importance of accurate forward modeling. To assess robustness to forward-model assumptions, we performed additional simulations incorporating source-space mismatch, forward-model mismatch, and more realistic noise conditions. While source-space mismatch produced only modest reductions in reconstruction accuracy, simulations generated using a BEM forward model and reconstructed using the single-shell model employed throughout this study showed larger performance reductions, particularly at lower SNRs and higher co-registration errors. Nevertheless, the qualitative dependence of reconstruction accuracy on SNR and co-registration precision remained unchanged across all robustness analyses. These findings suggest that the principal conclusions are reasonably robust to moderate model mismatch while also highlighting forward-model accuracy as a primary determinant of achievable laminar precision in empirical data. Future work could improve laminar inference precision by employing more sophisticated forward models, such as BEM or finite element models (FEMs). Targeted sensory paradigms that preferentially engage specific cortical laminae may also provide an important avenue for further validation of laminar specificity.

The sensitivity of laminar inference to orientation errors suggests that future improvements may come from more realistic models of cortical column geometry. In the present framework, corresponding vertices across layers share a common orientation, but cortical columns are not strictly straight^56^. Future models could therefore incorporate depth-varying orientation vectors derived from equivolumetric laminar surfaces^40^, as opposed to the equidistant surfaces used here. Estimating cortical column orientation using diffusion tensor imaging (DTI) is a promising alternative, as anisotropic diffusion within cortical gray matter can reflect the organization of cortical columns^57^. However, DTI-based estimates can be confounded by white matter fibers projecting into the cortex (the gyral bias effect), potentially biasing orientation measures. Advanced diffusion imaging techniques such as high angular resolution diffusion imaging (HARDI) and constrained spherical deconvolution, combined with surface-based analyses that sample diffusion metrics along radial cortical trajectories, can help isolate cortical diffusion signals from adjacent white matter influences^58,59^. Integrating these advanced imaging approaches into laminar MEG source models could yield more biologically plausible dipole orientation estimates and potentially improve depth localization, especially in highly folded cortex. The current framework provides a foundation for evaluating such models and for testing how column curvature influences laminar MEG source reconstructions.

Despite the significant improvements offered by our multilayer approach, a key limitation is the inability of current model-based inference methods to resolve two simultaneous sources of equal magnitude at different depths. Additional simulations including three weaker interfering sources at separate cortical locations produced only modest reductions in reconstruction accuracy, suggesting that the framework is reasonably robust to low-level concurrent activity. However, the inability to resolve two simultaneous sources of comparable magnitude at different depths indicates that the current approach should be interpreted as identifying the dominant laminar contribution within a localized cortical region and time window, rather than independently resolving all simultaneously active laminar generators. Although exact temporal alignment and perfectly matched source magnitudes are unlikely in biological systems, this limitation highlights the constraints of relying solely on Bayesian model comparison. One potential solution would be to use sensors capable of measuring higher spatial-frequency components of the magnetic fields, such as optically pumped magnetometers (OPMs), which could increase sensitivity to subtle spatial differences between closely spaced sources^60^. Future refinements should explore both sensor technology advances and alternative inference strategies, such as Bayesian model averaging (BMA), which assigns probabilistic weights to multiple competing models rather than selecting a single best-fit model^61^. These combined approaches may enhance the simultaneous resolution of multiple sources, overcoming a fundamental limitation of the current winner-takes-all approach.

A methodological consideration is the spatial extent of the underlying cortical generator. As shown in our simulations and previous hpMEG studies^19,20^, mismatches between the true spatial extent of cortical activity and the patch size assumed during source reconstruction can systematically bias laminar inference. Specifically, underestimation of source extent biases solutions toward deeper layers, whereas overestimation biases solutions toward superficial layers (Supplementary Fig. 17). These findings highlight a fundamental limitation of current non-invasive laminar inference approaches: cortical depth and spatial extent are partially confounded. Importantly, however, the principal empirical findings proved largely robust across patch sizes ranging from 2.5 to 10 mm (Supplementary Fig. 18). In the visual dataset, the dominant granular and supragranular responses remained evident across all tested patch sizes. The MF and MEF-I components exhibited highly similar laminar profiles despite minor differences in the significance of some supragranular effects. This suggests that the reported results are not driven by a particular choice of patch size. Nonetheless, an alternative interpretation of apparent changes in laminar dominance is that activity remains within the same laminar compartment while varying in spatial extent over time. Based on invasive recordings demonstrating layer-specific recruitment during visual processing and computational models predicting distinct laminar contributions to sensory and sensorimotor evoked responses^47,48,52–55^, we consider genuine changes in laminar engagement a plausible explanation for the observed dynamics. However, current non-invasive methods cannot fully dissociate changes in depth from changes in spatial extent, and the observed dynamics may reflect a combination of both processes.

Another methodological consideration concerns the temporal window over which model evidence is estimated. In supplementary simulations using different sliding window sizes (15, 25, 50, and 100 ms; Supplementary Fig. 1), we assessed the temporal sensitivity of laminar model evidence. Across all window lengths, model-based inference consistently recovered the true laminar depth, but the temporal profile of model evidence differed. Narrow windows (e.g., 15 ms) yielded sharper but noisier evidence profiles that collapsed earlier, while wider windows (e.g., 100 ms) integrated more information, resulting in broader and more stable evidence plateaus over time. Consistent with these simulations, empirical analyses repeated using 10, 25, and 50 ms windows (Supplementary Fig. 19) showed that short windows better resolved brief laminar transients, whereas longer windows produced smoother and more temporally extended evidence profiles. For example, the transient granular response in the visual ERF was most clearly detected using a 10 ms window, whereas the characteristic infra-/supragranular dynamics of the MF and MEF-I were more robustly detected using 25-50 ms windows. This trade-off reflects the balance between temporal resolution and sensitivity: short windows enable finer tracking of rapid changes but require high SNR, whereas longer windows are more robust to noise but may obscure transient laminar dynamics. Accordingly, window size should be selected according to the expected temporal scale of the neural process under investigation.

More generally, preprocessing choices may also influence the temporal and spatial characteristics of the reconstructed signals. For example, the button-press datasets were preprocessed using temporally extended signal space separation (tSSS)^62^, whereas the visual dataset was not. Although tSSS can alter the covariance structure of the sensor data, the broadly consistent physiological patterns recovered across these independently acquired datasets suggest that the principal laminar findings are not driven by this preprocessing step. Nevertheless, the influence of preprocessing approaches, including tSSS and related denoising methods, on laminar inference accuracy remains an important topic for future work.

While the present study focuses on time-resolved laminar inference of evoked responses, an important extension will be to characterize laminar dynamics of oscillatory activity. Previous work has examined laminar aspects of oscillatory activity using two-layer models, including frequency-resolved power analyses^24^ and burst-aligned averaging approaches^25^. However, these approaches either rely on steady-state power estimates or recast transient burst events as ERF-like signals, and therefore do not capture fully time-frequency-resolved laminar interactions. Extending the current multilayer framework to such non-phase-locked dynamics will require analyses that explicitly capture transient and frequency-specific activity, as well as inference methods capable of resolving interlaminar interactions. Developing such approaches represents an important direction for future work.

A complementary methodological direction is to move beyond model-comparison-based laminar inference altogether. A more direct, though equally technically challenging, approach would be to extract source signals directly from each layer of a multi-layer surface model across the cortex, rather than relying on model-based inference at specific vertices. Such an approach could be computationally tractable at the whole-cortex level and might resemble laminar local field potential (LFP) recordings, potentially enabling direct current source density (CSD) estimation and facilitating direct comparisons with laminar LFP data and laminar biophysical neural modeling^25^. However, even laminar LFP recordings are extracellular and thus remain subject to an inverse problem due to aggregation of diverse signal sources that is typically ignored in conventional CSD analyses^13^. Moreover, applying CSD-like methods to hpMEG presents significant challenges, including sign ambiguity and sensitivity to noise.

A practical consequence of the sensitivity to co-registration error identified in the simulations is the need to minimize dynamic head motion during acquisition. The most effective way to mitigate motion is through individualized head-casts, which substantially reduce head displacement and can realistically achieve < 2 mm co-registration error^21,22^. Alternatively, wearable customized arrays of OPMs could achieve similar precision by moving with the head, thus inherently reducing motion-induced errors. While head-casts or wearable OPM arrays do not eliminate movement entirely, residual movement artifacts can be addressed using additional correction strategies, including tSSS^62^, which was applied in the analysis of the button-press datasets presented here. Combining physical stabilization with signal-level correction may be necessary to reach the precision required for robust laminar inference in practice.

An important challenge for future applications of multilayer hpMEG is the development of subject-specific quality-control procedures. Although our simulations define general requirements for reliable laminar inference, the achievable accuracy in any individual dataset will depend on factors that are difficult to quantify directly, including forward-model accuracy, cortical column orientation estimates, co-registration precision, and local anatomical properties. One promising approach would be to incorporate dedicated localizer paradigms within the same recording session. For example, robust visual or sensorimotor evoked responses with well-characterized laminar activation sequences could be used to identify cortical locations with favorable anatomical and biophysical properties for laminar inference, quantify achievable signal-to-noise ratios, and constrain subsequent analyses of task-related activity. Such subject-specific reference measurements may provide a practical means of assessing the reliability of laminar inference in individual participants and of tailoring acquisition and analysis strategies to maximize reconstruction fidelity.

The ability to resolve sources with laminar precision necessitates a reevaluation of the theoretical limits of MEG resolution. The estimated number of pyramidal cells required to generate a detectable MEG signal ranges from 10,000 to 50,000, with the majority of the signal arising from laminae II/III and V^63^. Neuronal density varies substantially across the cortex, from approximately 20,000 cells/mm^2^ in frontal regions to around 45,000 in sensorimotor areas and up to 90,000 in the foveal region of primary visual cortex (V1)^64^. Factors such as cortical thickness, gyrification, and orientation of cortical columns relative to the sensors further constrain spatial resolution and depth precision. However, even with conventional MEG recording (without head-casts), spatial resolution as fine as ∼1 mm^2^ in V1^65^ and ∼3.5 mm^2^ in the sensorimotor cortex^66^ has been demonstrated. Given the enhancements offered by hpMEG, achieving laminar resolution is a realistic goal. If it is possible to resolve a source on the cortical surface with millimeter precision, the same principles should, in theory, allow for distinguishing sources at different cortical depths provided the right conditions are met.

Given these constraints, it is natural to ask how hpMEG compares to ultra-high field fMRI (7 - 11.7 Tesla) for laminar investigations. While fMRI provides spatially precise laminar profiles, it is fundamentally limited by its reliance on the hemodynamic response, which is subject to vascular drainage effects that can result in superficially biased laminar profiles. Efforts to mitigate this issue, such as deconvolution, vascular correction techniques, and contrast-based experimental designs, have improved laminar fMRI but do not fully address this limitation^18^. Another important fMRI limitation is its low temporal resolution (e.g. ∼12 s per whole-brain acquisition at 0.8 mm resolution^67^). As a result, many studies restrict laminar fMRI acquisition to a limited part of the brain in order to achieve better temporal resolution. hpMEG, in contrast, provides millisecond-resolution measurements of neuronal population activity with whole-head coverage and is unaffected by vascular artifacts, making it uniquely suited for studying activity with rapid laminar dynamics such as beta bursts^23^, induced oscillatory activity^24^, and event-related responses. However, because MEG requires anatomical imaging to define laminar boundaries, its accuracy depends on high-quality MRI-derived cortical reconstructions. Although recent attempts have been made to generate individualized laminar maps over the whole cortex using multimodal MRI and machine learning^68–70^, accurately achieving this for the whole cortex remains a major technical challenge. Successfully addressing this issue will require integrating advanced multimodal imaging approaches, ultra-high-field MRI, movement correction, and sophisticated computational methods. Current hpMEG analyses must therefore rely on laminar histological atlases based on a single, *ex vivo* brain such as the BigBrain atlas^40^. Developing methods for individualized laminar boundary estimation *in vivo* would represent a major step forward in both hpMEG and laminar fMRI research.

Emerging OPM technologies hold promise for further enhancing laminar inference by offering improved signal quality, allowing flexible sensor placement, and reducing operational costs^60^. While still in its early stages, simulation studies suggest that OPM-based MEG, if configured correctly, can achieve laminar resolution comparable to classic, SQUID-based MEG, at least for bilaminar models^34^. Future work should assess whether these benefits extend to true laminar inference. Both our results and those of Helbling^34^ suggest that incorporating individual anatomical features such as cortical thickness and curvature could optimize OPM sensor placement to capture higher spatial-frequency components of the magnetic fields or achieve higher effective SNRs, potentially improving laminar precision within targeted cortical regions. Simulations could help refine these configurations, eliminating the need for time-consuming trial-and-error adjustments. Such targeted precision may be particularly useful for pre-surgical mapping of eloquent cortical regions or for resolving lamina-specific computations in particular regions of interest.

In summary, we provide a framework for multilayer hpMEG and define the conditions under which depth-resolved laminar inference may be feasible. By systematically exploring the trade-offs between signal quality, anatomical constraints, and inference accuracy, we delineate best-case and marginal conditions for MEG-based laminar inference. While current implementations reliably support deep-versus-superficial distinctions and laminar-family-level interpretations, resolving individual laminae *in vivo* remains contingent on favorable conditions and requires further validation. These findings further suggest that there is unlikely to be a universal minimum trial count required for laminar inference. Instead, the achievable precision will depend on the interaction between trial count, component magnitude and variability, preprocessing quality, head stabilization, and recording noise characteristics, all of which determine the effective SNR of the evoked response. Future developments, including Bayesian model averaging, individualized laminar and orientation priors, more accurate forward models, and emerging sensor technologies, may further improve hpMEG’s ability to resolve cortical computations at the laminar level. These advances pave the way for a new era of laminar human neuroscience, bridging invasive electrophysiology with whole-brain, millisecond-resolution functional imaging.

## METHODS

### Empirical Data Acquisition

We validated multilaminar MEG inference using three independent human datasets comprising visual and motor paradigms acquired with high-precision MEG. Across datasets, healthy adult participants underwent structural MRI for individual cortical surface reconstruction and MEG recordings acquired with whole-head CTF systems at high sampling rates, with head position stabilized using participant-specific head-casts and continuously monitored using fiducial coils. The datasets span visually evoked responses in primary visual cortex, as well as self-paced and visually cued button presses engaging precentral and postcentral cortices, thereby sampling distinct cortical regions, task demands, and acquisition sites. Despite differences in experimental design, all datasets shared a common preprocessing and analysis framework, enabling consistent laminar source reconstruction and cross-dataset comparison of laminar activation patterns. Full details of participants, tasks, MRI and MEG acquisition protocols, preprocessing pipelines, and analysis parameters for each dataset are provided in the Supplementary Methods.

### Empirical Data Preprocessing

For all three empirical datasets, individual cortical surfaces were reconstructed from structural T1- and T2-weighted MRI using FreeSurfer *recon-all* (v6.0.1; see Supplementary Methods for dataset-specific MRI protocols)^26^. We then used FreeSurfer’s *mris_expand* function to generate 9 equidistant intermediate surfaces between the pial boundary surface and white matter boundary surface, yielding 11 cortical surfaces in total (Figure 1b). An arbitrary number of surfaces could be used, but we chose 11 for computational efficiency. These intermediate surfaces are not meant to represent the 6 cortical laminae, but rather to sample the cortical depth like a laminar electrode with multiple, evenly spaced contacts. Freesurfer operates on hemispheres independently, resulting in deep vertices and mesh faces cutting through subcortical structures. These vertices and associated faces were removed from each hemisphere of each layer mesh. The left and right hemisphere meshes of each intermediate surface were then combined by concatenation of their vertices and faces (left then right).

The resulting bihemispheric meshes each contained approximately 300,000 vertices, exceeding computational feasibility for source reconstruction. FreeSurfer generates the pial mesh by expanding the white matter boundary mesh until the pial boundary is reached. Thus, every pial mesh vertex has a corresponding white matter mesh vertex. The vectors linking these corresponding vertices, known as link vectors^27^, have been shown to provide the best estimate of cortical column orientation out of available methods^28^. One advantage of link vectors for constraining dipole orientation is that the vectors are equivalent across corresponding vertices in all intermediate surfaces. This ensures that model-based layer comparison is based solely on dipole location. However, most methods for mesh downsampling disrupt the correspondence between vertices when applied independently to each layer as they alter vertex coordinates rather than simply removing vertices. We therefore downsampled the pial boundary surface by a factor of 10, using the *vtkDecimatePro* function of the VTK library (v9.3.0)^29^ because it is purely subtractive (i.e. it only removes vertices), resulting in a mesh with approximately 30,000 vertices. We then downsampled the other 10 meshes by removing the same vertices that were removed from the pial surface, thus preserving vertex correspondence, and copying the pial mesh face structure (i.e. edges between vertices)^28^. This enabled a straightforward computation of link vectors (Figure 1c), which were used to constrain dipole orientations in all of the main simulations and analyses.

MEG data were preprocessed separately for each dataset, time-locked to the event of interest (visual grating onset or button press), and averaged over trials to obtain event-related fields used for laminar source reconstruction (details in Supplementary Methods).

### Simulations

We tested the efficacy of each analysis method using synthetic datasets based on the cortical surfaces and MEG data of a single participant from the visually cued button-press dataset (see Supplementary Methods). The MEG data was only used to define the sensor layout, sampling rate (600 Hz), number of trials (531), and number of samples (1201) for the simulations; the MEG sensor data itself was discarded. All simulations and analyses were implemented using the laMEG software package (v0.0.7; https://github.com/danclab/laMEG), built on a custom version of SPM^30^ compiled as a python library (https://github.com/danclab/spm), and are available at http://github.com/danclab/multilaminar_sim.

For each simulation, we specified a source centered at a vertex on one of the 11 cortical surfaces spanning the depth of the cortex from the pial to the white matter surface. We simulated a localized patch of current density with a Gaussian-shaped temporal profile over a 2 s time window with a peak at the center of the window and full width at half maximum (FWHM) of 400 ms (time course displayed in Supplementary Fig. 1). The spatial extent of the simulated source was set to 5 mm FWHM (corresponding to a mean patch size of 14.93 vertices). To generate synthetic MEG data, we used a single-shell forward model^31^ based on a surface composed of the 11 cortical layer meshes concatenated into a single mesh. The dipole moment for each simulation was set to 10 nAm unless otherwise specified. We selected 100 random cortical locations (vertices on the pial surface) and conducted separate simulations for different sensor-level signal-to-noise ratio (SNR) levels (−500, -50, -35, -20, - 10, -5, 0, and 5 dB) and co-registration error levels (0, 0.5, 1, 2, 3, 4, and 5 mm). White noise was added at the sensor level, scaled to achieve the desired per-trial SNR (computed as the ratio of signal power to noise power across all sensors)^20^.

To simulate MEG-MRI co-registration error, we applied a random rigid-body perturbation to the MRI-defined nasion (NAS), left preauricular (LPA), and right preauricular (RPA) fiducials. Let *fi* denote the three fiducial coordinates and 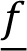 their barycenter; we first centered the fiducials by 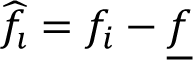, then drew (i) a random translation *t* with isotropic direction and normally distributed signed magnitude with scale equal to the target error level (mm), and (ii) a random rotation *R* about an isotropically distributed axis with rotation-vector magnitude drawn from a zero-mean normal distribution scaled by the target error level. We applied the transform about the barycenter, 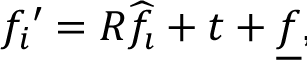, which preserves inter-fiducial distances (i.e., a rigid triangle) but yields different displacements at each fiducial due to rotation. Because the combined translation+rotation does not, in general, make every fiducial move by the same amount, we operationalized the “error level” as the maximum fiducial displacement, 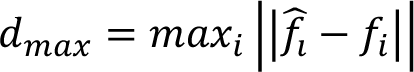, and used rejection sampling to redraw (*t*, *R*) until l*d*_max_ − *d*_target_l < 0.05 mm. The resulting perturbed fiducials (*NAS′*, *LPA′*, *RPA′*) were then used for MEG-MRI co-registration, thereby simulating misalignment between the subject’s head position in the MEG scanner and their anatomical MRI^20^. This operationalization provides a simple and well-controlled measure of co-registration error, but may overestimate the effective misalignment relative to more spatially distributed or temporally varying head movement. Because realistic head motion and its correction were not explicitly simulated, the reported error levels should be interpreted as conservative upper bounds.

To evaluate the accuracy of laminar source reconstruction in the presence of multiple simultaneous sources, we conducted a separate set of simulations in which two sources of equal strength were generated at different cortical depths. For each of the 100 selected cortical locations, we fixed one source in the middle layer while systematically varying the second source from the most superficial to the deepest cortical surface. The simulated sources followed the same Gaussian-shaped temporal profile as in the single-source simulations, with a dipole moment of 10 nAm and a spatial extent of 5 mm FWHM. Synthetic MEG data were generated using a single-shell forward model, with white noise added to achieve a sensor-level SNR of -20 dB.

To evaluate the impact of different cortical column orientation estimation methods on laminar source reconstruction accuracy, we systematically varied the approach used to define dipole orientations during both simulation and reconstruction. We tested seven distinct methods, including surface normal vectors computed from the original high-resolution mesh^28^ and a downsampled surface mesh^32^, cortical patch statistics^33^, and link vectors^27^, with additional conditions for whether dipole orientations were fixed across layers. For each method, we simulated sources at the 100 selected cortical locations, assigning simulated dipole orientations based on the selected approach and then reconstructing sources using each of the different orientation definitions. The downsampled surface normal method estimated dipole orientation at each vertex of the decimated cortical mesh by averaging the normal vectors of all adjacent triangular faces. This was implemented using the *spm_mesh_normals* function in SPM^30^. The original surface normal method followed the same approach but was applied to the full-resolution cortical surface before being mapped onto the decimated mesh for source localization^28^. The cortical patch statistics method calculated the mean normal vector at each vertex of the downsampled mesh using all neighboring vertices from the original high-resolution surface^33^. In contrast, the link vector method defined dipole orientation as the vector linking each pial surface vertex to its corresponding white matter vertex^27^, leveraging the one-to-one correspondence maintained during mesh decimation. Because different orientation estimation methods can yield subtly different vector fields, we assessed the angular discrepancies between these definitions and their effects on laminar inference accuracy. Dipole orientation vectors derived from each method were incorporated into the lead field matrix used for source inversion^28^ in order to evaluate how orientation differences impact model evidence and depth localization accuracy across simulations. For each of these methods, we tested versions where the orientation was computed independently for each layer and versions where it was computed on the pial surface and fixed across layers (except for the link vectors method which is equivalent across layers by definition).

To assess the extent to which the primary simulation results depended on simplifying assumptions, we performed four additional robustness analyses. First, the SNR and co-registration simulations were repeated using temporally autocorrelated pink noise (1/f amplitude spectrum) generated independently at each sensor and scaled to the same target RMS noise level as the original white-noise simulations. Second, to assess robustness to concurrent activity, three additional interfering sources were added at randomly selected cortical locations in the middle cortical layer. These sources had independent band-limited temporal profiles (10-30 Hz) and amplitudes equal to 10% of the primary simulated source. Third, to evaluate the effect of source-space mismatch, simulations were generated using an alternative generative model (AGM)^34^ based on multilayer cortical surfaces containing 32,399 vertices per layer, whereas source reconstruction was performed using the source model surfaces employed throughout the main analyses (29,130 vertices per layer), generated using a different surface downsampling factor (11% rather than 10%). This resulted in systematic differences between simulated dipole locations and the source locations assumed during inversion. Finally, to assess the effect of forward-model mismatch, simulated data were generated using lead fields computed with a boundary element model (BEM) forward model (computed from subject-specific inner-skull, outer-skull, and scalp surfaces using OpenMEEG v2.1.0^35^), while source reconstruction was performed using the Nolte single-shell model used throughout the study. For each robustness analysis, the same SNR and co-registration error manipulations as in the primary simulations were applied, and reconstruction accuracy, bias, and model-evidence structure were evaluated using identical procedures (see Simulation Analysis, below).

To examine how anatomical features influence the accuracy of laminar source reconstruction, we generated a large-scale set of simulations in which sources were placed at every vertex of a layered cortical mesh (one source per simulation). The mesh consisted of 29,130 locations at 11 cortical depths, resulting in a total of 320,430 simulated sources. Each simulation used a Gaussian-shaped temporal profile with a full width at half maximum (FWHM) of 400 ms and a dipole moment of 10 nAm (Supplementary Fig. 1 includes the time course of this simulated signal). Synthetic MEG data were generated using a single-shell forward model, and white noise was added to yield a sensor-level SNR of -20 dB. All of these simulations were conducted with zero co-registration error.

## Simulation Analysis

For each simulated dataset, we applied multiple sparse priors (MSP) inversion^39^ to each simulated dataset within a Hann windowed 50 ms analysis window centered around the peak of the simulated Gaussian temporal profile. These data were projected into 274 orthogonal spatial modes and 4 temporal modes, with a patch size of 5 mm (unless otherwise specified) and a single prior corresponding to the vertex the activity was simulated at. This was done to test the feasibility of a sliding time window approach previously applied to bilaminar MEG dynamics^25^. Because source reconstruction and model comparison is performed in each time window independently in this sliding time window framework, laminar inference was evaluated within a 50 ms window centered on the known peak of the simulated response, rather than across the full time course. We confirmed the robustness of this window size in supplementary simulations using sliding time windows of varying lengths (15, 25, 50, and 100 ms; Supplementary Fig. 1) across the whole epoch, which showed consistent reconstruction performance across a broad temporal range. Rather than performing source reconstruction on a single multilayer cortical mesh, we conducted separate MSP inversions for each of the 11 cortical depth surfaces, treating each as an independent forward model. For each simulation, we computed the model evidence (free energy) for each depth-specific source reconstruction and identified the cortical surface with the highest model evidence (Figure 1e). This allowed us to infer the most likely depth of the simulated source by comparing the relative free energy across reconstructions at different cortical depths (Figure 1f).

We quantified reconstruction accuracy by identifying, for each simulated cortical depth and vertex, the surface with the highest model evidence and comparing its depth to the known simulated source depth. Reconstruction error was defined as the absolute difference between the simulated and inferred depths (Figure 1g), measured both in number of surfaces and in millimeters by scaling surface indices using local cortical thickness. Reconstruction bias was defined as the signed difference between simulated and inferred depths (Figure 1h). These quantities were first computed per vertex and simulated depth, and then averaged across depths and cortical locations to obtain summary measures. Note that while mean model evidence matrices often exhibit a diagonal structure, indicating correct depth identification on average, reconstruction error and bias capture variability at the level of individual vertices and simulations, and are therefore sensitive to deviations that are not apparent in the averaged matrices.

To assess statistical significance, we performed permutation tests using 10,000 iterations, shuffling the free energy matrices for each simulated cortical depth and recomputing error and bias distributions under the null hypothesis of no depth sensitivity. P-values were calculated by comparing observed reconstruction errors and biases against these shuffled distributions, with statistical significance determined at *p* < 0.05. Effect sizes were defined as standardized differences relative to the shuffled null distribution, computed as the difference between the observed mean and the mean of the shuffled distribution divided by the standard deviation of the shuffled distribution. We computed 95% confidence intervals for reconstruction error and bias using bootstrap resampling (5,000 iterations). For each SNR or co-registration error level, error and bias were first averaged across simulated depths within each cortical location; cortical locations were then resampled with replacement and the mean error or bias was recomputed for each bootstrap sample.

We also computed a diagonal dominance score to quantify the extent to which model evidence matrices favored reconstructions at the correct depth. This score was defined as the proportion of total matrix mass accounted for by the diagonal values (i.e., the sum of diagonal values divided by the sum of all values in the free energy matrix). This metric was compared against a permuted distribution (*N* = 10,000) to assess whether observed diagonality exceeded chance levels.

We accounted for cortical laminar thickness variability by incorporating the BigBrain atlas^40^. Because cortical laminae differ in thickness across the brain (Figure 1d), mapping simulated and inferred sources to cytoarchitectonic laminae rather than fixed-depth surfaces improves biological interpretability. To achieve this, we mapped the BigBrain atlas to the subject’s anatomy via FreeSurfer’s surface-based registration through fsaverage space^41^. This allowed us to align the proportional cortical laminae boundaries from the atlas with the subject’s individual cortical geometry. For each simulated cortical location, we extracted the proportional laminae boundaries (six depth values defining the transitions between laminae) using precomputed laminar thickness estimates from the atlas. These boundaries were then scaled by the local cortical thickness to derive laminar depths in millimeters. The final mapping step assigned each of the 11 cortical surface layers used for source reconstruction to a corresponding cytoarchitectonic laminae. Reconstruction errors were then computed in laminar space rather than in fixed surface coordinates; specifically, an inferred source depth was only considered an error if it fell outside the lamina of the simulated source, regardless of its precise position in cortical depth. Significance was evaluated using the same permutation-based approach used for error in terms of layers and millimeters (*N* = 10,000).

To evaluate how local anatomy influences laminar inference, we quantified reconstruction error at each cortical location and related it to anatomical features. For each vertex, we computed absolute reconstruction error in millimeters. Given the highly skewed distribution of errors, with exact recovery (zero error) occurring in the majority of cortical locations, we additionally defined a binary error variable (zero vs. non-zero error) for classification-based analyses. This binary variable was used in receiver operating characteristic (ROC) analyses to identify anatomical features predictive of correct versus incorrect laminar inference. Continuous error values were retained for descriptive analyses.

We then analyzed the relationship between binary reconstruction error and several anatomical properties: cortical thickness (measured between the pial and white matter surfaces), distance to the scalp surface, orientation of the cortical column relative to the scalp, and variability in lead field magnitude across depth. Orientation was computed as the absolute dot product between the simulated dipole orientation and the scalp normal at each vertex. Lead field variability was quantified as the root mean square difference in lead field magnitude across surface layers, relative to the superficial (pial) surface. These features were then related to binary reconstruction error using kernel density estimation and receiver operating characteristic (ROC) analysis to identify anatomical thresholds predictive of accurate inference. Optimal thresholds were defined as the values that maximized Youden’s *J* statistic (sensitivity - specificity) on the ROC curve, after ensuring consistent feature polarity (AUC ≥ 0.5). To assess whether anatomical features jointly predict laminar inference accuracy better than single features, we performed multivariate logistic regression and random forest classification. We evaluated model performance using 5-fold cross-validated ROC analysis. Logistic regression models were tested both with and without pairwise interaction terms to quantify potential nonlinear feature interactions. Random forest hyperparameters were set to 500 trees, unlimited depth, and balanced class weights.

To further characterize the relationships between anatomical features and laminar inference accuracy, we performed a mediation analysis treating lead-field difference as a mediator. For each anatomical feature (dipole orientation, scalp-to-cortex distance, and cortical thickness), we estimated (i) a linear regression model relating the feature to lead-field difference, and (ii) a logistic regression model relating laminar inference accuracy (binary: zero vs. non-zero reconstruction error) to both the feature and lead-field difference. Indirect (mediated) effects were computed as the product of the regression coefficients linking the feature to the mediator and the mediator to the outcome. Confidence intervals for indirect effects were obtained using bootstrap resampling (5,000 iterations) over cortical vertices.

## Empirical Data Analysis

Laminar source reconstruction used the laMEG software package (v0.0.7; https://github.com/danclab/laMEG), built on a custom version of SPM^30^ compiled as a python library (https://github.com/danclab/spm).

For each participant and session, candidate sources were first localized by co-registering the averaged epochs to the combined 11-layer mesh (formed by concatenating all depth surfaces into a single source space), and then using an empirical Bayes beamformer (EBB) with a 5-mm patch size, four temporal modes, and a number of spatial modes equal to the rank of the preprocessed MEG data, within a task- and component-specific time window (see Supplementary Methods for dataset- and component-specific details). All laminar inference analyses were performed on trial-averaged ERFs, rather than on single-trial data. Sensor-level SNR was therefore used as the primary measure of data quality, reflecting the reliability of the averaged response (Supplementary Fig. 14). Sensor-level SNR was computed within the analysis window as the ratio in dB of the RMS of the trial-averaged signal across channels to the RMS of trial-by-trial deviations from that mean, yielding a single SNR time series per participant. Participants whose SNR within this window was < -15 dB were excluded (visually evoked response dataset: *N* = 6; visually cued button press dataset: *N* = 0; self-paced button press dataset: *N* = 0). This EBB step was used solely for spatial localization and vertex selection; all laminar inference and model comparison were performed using multiple sparse priors (MSP), as described below.

To select a single vertex for laminar inference within each region of interest (primary visual, motor, or somatosensory cortex depending on the ERF and component), we combined functional and anatomical criteria. Source time series were extracted from the middle cortical surface, and vertices were ranked by mean signal magnitude within the component-specific analysis window relative to baseline (defined per dataset and component; see Supplementary Methods). Among vertices in the anatomically defined region of interest, we retained the top 1% by signal magnitude and selected the vertex with the highest composite anatomical suitability score. This score was defined as the sum of z-scored cortical thickness, deep-superficial lead-field RMS difference, alignment between cortical column orientation and local scalp normal, and inverse distance to the scalp, thereby favoring locations with both strong signal and favorable anatomical properties for laminar discrimination.

At the selected vertex, sliding-window Bayesian model comparison was then performed across all 11 cortical depths. We applied multiple sparse priors (MSP) inversion to each dataset within overlapping windows (10 ms for the visually evoked response dataset, 25 ms for the button press datasets), with Hann windowing applied within each window, a number of spatial modes equal to the rank of the preprocessed data, 4 temporal modes, a patch size of 5 mm, and a single prior corresponding to the selected vertex. Separate sliding window MSP inversions were performed for each of the 11 cortical depth surfaces, treating each surface as an independent forward model. This yielded model evidence (free energy) time series for each depth-specific source reconstruction, for each subject and session.

In order to interpret the cortical depth surfaces in terms of laminae, we mapped the BigBrain atlas to each subject’s anatomy, computed proportional laminae boundaries from the atlas, scaled these boundaries by the local cortical thickness, and assigned each of the 11 cortical surface layers to a corresponding cytoarchitectonic lamina. Free energy values were then averaged within laminae. The resulting model evidence was then compared in two ways: i) by computing model evidence relative to the worst model (ΔF) at each time point, and ii) using Bayesian model selection to identify the most probable family of models at the group level at each time point. Because all cortical depth surfaces were generated by jointly downsampling the multilayer mesh while preserving one-to-one vertex correspondence across depth, each layer-specific source model contained the same number of dipoles and free parameters. Furthermore, dipole orientations were constrained using link vectors connecting corresponding vertices across surfaces, ensuring identical orientation priors across layer models. Consequently, differences in model evidence between layers reflect differences in source depth rather than differences in model complexity, dipole orientation, or source-space dimensionality.

Although differences in free energy (ΔF) greater than 3 are commonly interpreted as strong evidence in single-subject model comparisons, this heuristic does not extend to group-level inference. The ΔF > 3 criterion reflects within-subject evidence strength and cannot be used as a threshold for statistical significance across participants. In the present study, ΔF values were therefore used descriptively, and statistical inference was performed at the group level using a hierarchical Bayesian framework.

For Bayesian model selection we used the VBA (Variational Bayesian Analysis) toolbox^42^. The free energy measures of the six laminae were grouped in three families: supragranular (laminae I-III), granular (lamina IV) and infragranular (laminae V-VI) families. At each time point, family-wise random effects Bayesian model selection was applied to the subject-level free energy values. This approach treats model identity as a random variable across participants and estimates the population frequency of each laminar family, thereby accounting for inter-subject variability in model preference^43,44^. In the case when the posterior probability that all families are equally frequent in the population was below 0.5, we calculated protected exceedance probabilities (PEPs) to quantify the probability that one laminar family was more frequent in the population than the competing alternatives while accounting for chance-level differences in model frequency. Consequently, group-level inference was based on the consistency of model evidence across participants rather than on averaged free energy values.

## Data availability

The preprocessed MEG data, structural MRI scans, cortical surface reconstructions, and fiducial locations underlying the empirical analyses are available online (https://doi.org/10.5281/zenodo.20721185). Raw MEG and MRI datasets are available from the corresponding author upon reasonable request and subject to institutional ethical approvals and data-sharing agreements.

## Code availability

Experimental stimulus presentation and behavioral response collection were performed using task-specific software available at: https://github.com/jbonaiuto/cued_action_selection, https://github.com/danclab/auditory_laminar, and https://gitlab.com/TommyClausner/meglyon-experiment-runner. All custom data analysis code used in this study are available at https://github.com/danclab/multilaminar_sim/, https://github.com/danclab/laminar_motor_erf/, https://github.com/danclab/auditory_laminar, and https://github.com/cophyteam/laminar_visual_erf.

## Acknowledgements

We gratefully acknowledge support from the CNRS/IN2P3 Computing Center (Lyon - France) for providing computing and data-processing resources needed for this work. This work was supported by a Fondation pour l’Audition grant RD-2022-03 to J.J.B. and A.R.D., a European Research Council (ERC) consolidator grant 864550 to J.J.B., a French National Research Agency (ANR) grant HiFi (ANR-20-CE17-0023) to J.J.B and J.M.., a European Research Council (ERC) starting grant 716862 to M.B., a French National Research Agency (ANR) grant TheAlphaMEG (ANR-25-CE37-0908-01) to M.B, and the Discovery Research Platform for Naturalistic Neuroimaging funded by Wellcome [226793/Z/22/Z].

## Author contributions

J. J. B. conceived the study, supervised the work, acquired funding, developed the methodology and software, performed analyses, and wrote the manuscript. M. J. S., M. M., and D. Shelepenkov developed the methodology and software, performed analyses, generated visualizations, and contributed to writing the manuscript. I. A. contributed to software development, validation, formal analysis, visualization, and manuscript writing. Q. M. and S. G. contributed software and manuscript writing. J. J. B., M. F., C. F. P., Y. Z., F. L., and S. D. performed the experimental studies. M. B., J. M., S. D., F. L., and D. Schwartz. contributed to methodological development. A. R. D. and M. B. supervised the work. J. J. B., A. R. D., M. B., and J. M. acquired funding. All authors reviewed and edited the manuscript.

## References

1. Smith, C. U. M. A century of cortical architectonics. J. Hist. Neurosci. 1, 201–218 (1992).

2. Hirsch, J. A. & Martinez, L. M. Laminar processing in the visual cortical column. Curr. Opin. Neurobiol. 16, 377–384 (2006).

3. Harris, K. D. & Shepherd, G. M. G. The neocortical circuit: themes and variations. Nat. Neurosci. 18, 170–181 (2015).

4. Bastos, A. M., Loonis, R., Kornblith, S., Lundqvist, M. & Miller, E. K. Laminar recordings in frontal cortex suggest distinct layers for maintenance and control of working memory. Proc. Natl. Acad. Sci. 115, 1117–1122 (2018).

5. Bijanzadeh, M., Nurminen, L., Merlin, S., Clark, A. M. & Angelucci, A. Distinct laminar processing of local and global context in primate primary visual cortex. Neuron 100, 259–274 (2018).

6. Dykstra, A. R., et al. Testing circuit-level theories of consciousness in humans. Trends Cogn. Sci. In press, (2025).

7. Bastos, A. M. et al. Canonical Microcircuits for Predictive Coding. Neuron 76, 695–711 (2012).

8. Shipp, S., Adams, R. A. & Friston, K. J. Reflections on agranular architecture: predictive coding in the motor cortex. Trends Neurosci. 36, 706–16 (2013).

9. Borbély, S., Halasy, K., Somogyvári, Z., Détári, L. & Világi, I. Laminar analysis of initiation and spread of epileptiform discharges in three in vitro models. Brain Res. Bull. 69, 161–167 (2006).

10. Underwood, C. F. & Parr-Brownlie, L. C. Primary motor cortex in Parkinson’s disease: Functional changes and opportunities for neurostimulation. Neurobiol. Dis. 147, 105159 (2021).

11. Pigorini, A. et al. Simultaneous invasive and non-invasive recordings in humans: A novel Rosetta stone for deciphering brain activity. J. Neurosci. Methods 408, 110160 (2024).

12. Chung, J. E. et al. High-density single-unit human cortical recordings using the Neuropixels probe. Neuron 110, 2409–2421 (2022).

13. Gratiy, S. L., Devor, A., Einevoll, G. T. & Dale, A. M. On the estimation of population-specific synaptic currents from laminar multielectrode recordings. *Front*. Neuroinformatics 5, 32 (2011).

14. Caruso, L. et al. In Vivo Magnetic Recording of Neuronal Activity. Neuron 95, 1283–1291.e4 (2017).

15. Rosen, B. R. & Savoy, R. L. fMRI at 20: Has it changed the world? NeuroImage 62, 1316–1324 (2012).

16. Uhlhaas, P. J. et al. Magnetoencephalography as a Tool in Psychiatric Research: Current Status and Perspective. Biol. Psychiatry Cogn. Neurosci. Neuroimaging 2, 235–244 (2017).

17. Samuelsson, J. G., Peled, N., Mamashli, F., Ahveninen, J. & Hämäläinen, M. S. Spatial fidelity of MEG/EEG source estimates: A general evaluation approach. NeuroImage 224, 117430 (2021).

18. Lawrence, S. J. D., Formisano, E., Muckli, L. & De Lange, F. P. Laminar fMRI: Applications for cognitive neuroscience. NeuroImage 197, 785–791 (2019).

19. Troebinger, L., López, J. D., Lutti, A., Bestmann, S. & Barnes, G. Discrimination of cortical laminae using MEG. NeuroImage 102, 885–893 (2014).

20. Bonaiuto, J. et al. Non-invasive laminar inference with MEG: Comparison of methods and source inversion algorithms. NeuroImage 167, 372–383 (2018).

21. Troebinger, L. et al. High precision anatomy for MEG. NeuroImage 86, 583–591 (2014).

22. Meyer, S. S. et al. Flexible head-casts for high spatial precision MEG. J. Neurosci. Methods 276, 38–45 (2017).

23. Szul, M. J. et al. Diverse beta burst waveform motifs characterize movement-related cortical dynamics. Prog. Neurobiol. 228, 102490 (2023).

24. Bonaiuto, J. et al. Lamina-specific cortical dynamics in human visual and sensorimotor cortices. eLife 7, e33977 (2018).

25. Bonaiuto, J. J. et al. Laminar dynamics of high amplitude beta bursts in human motor cortex. NeuroImage 242, 118479 (2021).

26. Fischl, B. FreeSurfer. NeuroImage 62, 774–781 (2012).

27. Dale, A. M., Fischl, B. & Sereno, M. I. Cortical Surface-Based Analysis: I. Segmentation and Surface Reconstruction. NeuroImage 9, 179–194 (1999).

28. Bonaiuto, J. et al. Estimates of cortical column orientation improve MEG source inversion. NeuroImage 216, 116862 (2020).

29. Schroeder, W., Martin, K. M. & Lorensen, W. E. The Visualization Toolkit an Object-Oriented Approach to 3D Graphics. (Prentice-Hall, Inc., 1998).

30. Ashburner, J. et al. SPM12 manual. Wellcome Trust Cent. Neuroimaging Lond. UK 2464, (2014).

31. Nolte, G. The magnetic lead field theorem in the quasi-static approximation and its use for magnetoenchephalography forward calculation in realistic volume conductors. Phys. Med. Biol. 48, 3637–3652 (2003).

32. Hillebrand, A. & Barnes, G. R. The use of anatomical constraints with MEG beamformers. NeuroImage 20, 2302–13 (2003).

33. Lin, F.-H., Belliveau, J. W., Dale, A. M. & Hämäläinen, M. S. Distributed current estimates using cortical orientation constraints. Hum. Brain Mapp. 27, 1–13 (2006).

34. Helbling, S. Inferring laminar origins of MEG signals with optically pumped magnetometers (OPMs): A simulation study. Imaging Neurosci. 3, imag_a_00410 (2025).

35. Gramfort, A., Papadopoulo, T., Olivi, E. & Clerc, M. OpenMEEG: opensource software for quasistatic bioelectromagnetics. Biomed. Eng. OnLine 9, 45 (2010).

36. Brodmann, K. Vergleichende Lokalisationslehre Der Grosshirnrinde in Ihren Prinzipien Dargestellt Auf Grund Des Zellenbaues. (Barth, 1909).

37. von Economo, C. F. & Koskinas, G. N. Die Cytoarchitektonik Der Hirnrinde Des Erwachsenen Menschen. (J. Springer, 1925).

38. Wagstyl, K., Ronan, L., Goodyer, I. M. & Fletcher, P. C. Cortical thickness gradients in structural hierarchies. Neuroimage 111, 241–250 (2015).

39. Friston, K. et al. Multiple sparse priors for the M/EEG inverse problem. NeuroImage 39, 1104–1120 (2008).

40. Wagstyl, K. et al. Mapping Cortical Laminar Structure in the 3D BigBrain. Cereb. Cortex 28, 2551–2562 (2018).

41. Fischl, B., Sereno, M. I., Tootell, R. B. H. & Dale, A. M. High-resolution intersubject averaging and a coordinate system for the cortical surface. Hum. Brain Mapp. 8, 272–284 (1999).

42. Daunizeau, J., Adam, V. & Rigoux, L. VBA: a probabilistic treatment of nonlinear models for neurobiological and behavioural data. PLoS Comput. Biol. 10, e1003441 (2014).

43. Stephan, K. E., Penny, W. D., Daunizeau, J., Moran, R. J. & Friston, K. J. Bayesian model selection for group studies. NeuroImage 46, 1004–1017 (2009).

44. Rigoux, L., Stephan, K. E., Friston, K. J. & Daunizeau, J. Bayesian model selection for group studies — Revisited. NeuroImage 84, 971–985 (2014).

45. Belardinelli, P., Ortiz, E., Barnes, G. R., Noppeney, U. & Preissl, H. Source reconstruction accuracy of MEG and EEG Bayesian inversion approaches. PloS One 7, e51985 (2012).

46. Kinsbourne, M. Hemineglect and hemisphere rivalry. Adv Neurol 18, 41–49 (1977).

47. Yabuta, N. H. & Callaway, E. M. Functional streams and local connections of layer 4C neurons in primary visual cortex of the macaque monkey. J. Neurosci. 18, 9489–9499 (1998).

48. Yoshimura, Y., Dantzker, J. L. & Callaway, E. M. Excitatory cortical neurons form fine-scale functional networks. Nature 433, 868–873 (2005).

49. Callaway, E. M. Feedforward, feedback and inhibitory connections in primate visual cortex. Neural Netw. 17, 625–632 (2004).

50. Self, M. W., van Kerkoerle, T., Super, H. & Roelfsema, P. R. Distinct roles of the cortical layers of area V1 in figure-ground segregation. Curr. Biol. 23, 2121–2129 (2013).

51. Maier, A., Adams, G. K., Aura, C. & Leopold, D. A. Distinct superficial and deep laminar domains of activity in the visual cortex during rest and stimulation. Front. Syst. Neurosci. 4, 31 (2010).

52. Cheyne, D., Bakhtazad, L. & Gaetz, W. Spatiotemporal mapping of cortical activity accompanying voluntary movements using an event-related beamforming approach. Hum. Brain Mapp. 27, 213–229 (2006).

53. Cheyne, D. & Weinberg, H. Neuromagnetic fields accompanying unilateral finger movements: pre-movement and movement-evoked fields. Exp. Brain Res. 78, (1989).

54. Kristeva, R., Cheyne, D. & Deecke, L. Neuromagnetic fields accompanying unilateral and bilateral voluntary movements: topography and analysis of cortical sources. Electroencephalogr. Clin. Neurophysiol. Potentials Sect. 81, 284–298 (1991).

55. Oishi, M., Kameyama, S., Fukuda, M., Tsuchiya, K. & Kondo, T. Cortical activation in area 3b related to finger movement: an MEG study. Neuroreport 15, 57–62 (2004).

56. Bok, S. Der Einfluss in den Furchen und Windungen auftretenden Krümmungen der Grosshirnrinde auf die Rindenarchitektur. Z. Für Gesamte Neurol. Psychiatr. 121, 682–750 (1929).

57. Leuze, C. W. U. et al. Layer-Specific Intracortical Connectivity Revealed with Diffusion MRI. Cereb. Cortex 24, 328–339 (2014).

58. Gulban, O. F. et al. Cortical fibers orientation mapping using in-vivo whole brain 7 T diffusion MRI. Neuroimage 178, 104–118 (2018).

59. St-Onge, E., Daducci, A., Girard, G. & Descoteaux, M. Surface-enhanced tractography (SET). NeuroImage 169, 524–539 (2018).

60. Brookes, M. J. et al. Magnetoencephalography with optically pumped magnetometers (OPM-MEG): the next generation of functional neuroimaging. Trends Neurosci. 45, 621–634 (2022).

61. Hinne, M., Gronau, Q. F., van den Bergh, D. & Wagenmakers, E.-J. A Conceptual Introduction to Bayesian Model Averaging. Adv. Methods Pract. Psychol. Sci. 3, 200–215 (2020).

62. Taulu, S. & Simola, J. Spatiotemporal signal space separation method for rejecting nearby interference in MEG measurements. Phys. Med. Biol. 51, 1759 (2006).

63. Murakami, S. & Okada, Y. Contributions of principal neocortical neurons to magnetoencephalography and electroencephalography signals. J. Physiol. 575, 925–936 (2006).

64. Ribeiro, P. F. M. et al. The human cerebral cortex is neither one nor many: neuronal distribution reveals two quantitatively different zones in the gray matter, three in the white matter, and explains local variations in cortical folding. Front. Neuroanat. 7, (2013).

65. Nasiotis, K., Clavagnier, S., Baillet, S. & Pack, C. C. High-resolution retinotopic maps estimated with magnetoencephalography. NeuroImage 145, 107–117 (2017).

66. Barratt, E. L., Francis, S. T., Morris, P. G. & Brookes, M. J. Mapping the topological organisation of beta oscillations in motor cortex using MEG. NeuroImage 181, 831–844 (2018).

67. Chai, Y. et al. Unlocking near-whole-brain, layer-specific functional connectivity with 3D VAPER fMRI. Imaging Neurosci. 2, 1–20 (2024).

68. Trampel, R., Bazin, P.-L., Pine, K. & Weiskopf, N. In-vivo magnetic resonance imaging (MRI) of laminae in the human cortex. NeuroImage 197, 707–715 (2019).

69. Autio, J. A. et al. Charting cortical-layer specific area boundaries using Gibbs’ ringing attenuated T1w/T2w-FLAIR myelin MRI. bioRxiv 2024.09.27.615294 (2024) doi:10.1101/2024.09.27.615294.

70. Zeng, X. et al. Segmentation of supragranular and infragranular layers in ultra-high-resolution 7T *ex vivo* MRI of the human cerebral cortex. Cereb. Cortex 34, bhae362 (2024).

