## Supplementary Figures and Methods for "Multilayer MEG source modelling enables depth-resolved laminar inference in humans"

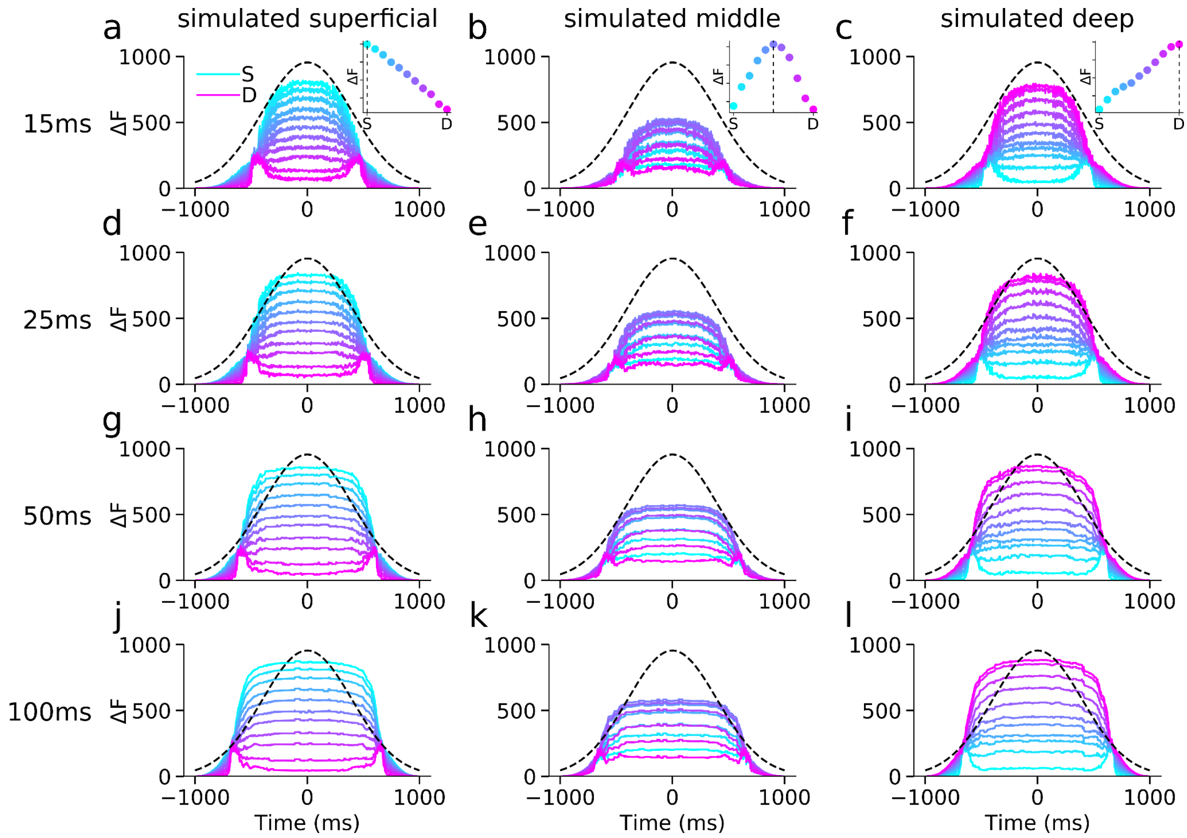


Supplementary Figure 1. **Effect of sliding window size on laminar inference accuracy.** Laminar model evidence over time (ΔF, relative to worst model at each time point) for simulated dipolar sources across 11 cortical depths, shown for sliding window sizes of 15 ms, 25 ms, 50 ms, and 100 ms (rows). Columns correspond to sources simulated at superficial, middle, and deep cortical depths. Each trace reflects the average ΔF across 100 simulated cortical locations at a fixed sensor-level SNR of -20 dB. Colors indicate cortical evaluation depth (S = superficial surface layer, D = deep surface layer). The black dashed line shows the simulated signal time course. Inset panels in the first row (a-c) show model evidence at t = 0ms, with peak ΔF values aligning with the simulated source depth (dashed vertical line). The depth-specific pattern of model evidence was consistent across window sizes for each simulated depth (columns). Results confirm that laminar model discrimination is consistent across a wide range of window sizes, with all tested windows successfully recovering the correct source depth.

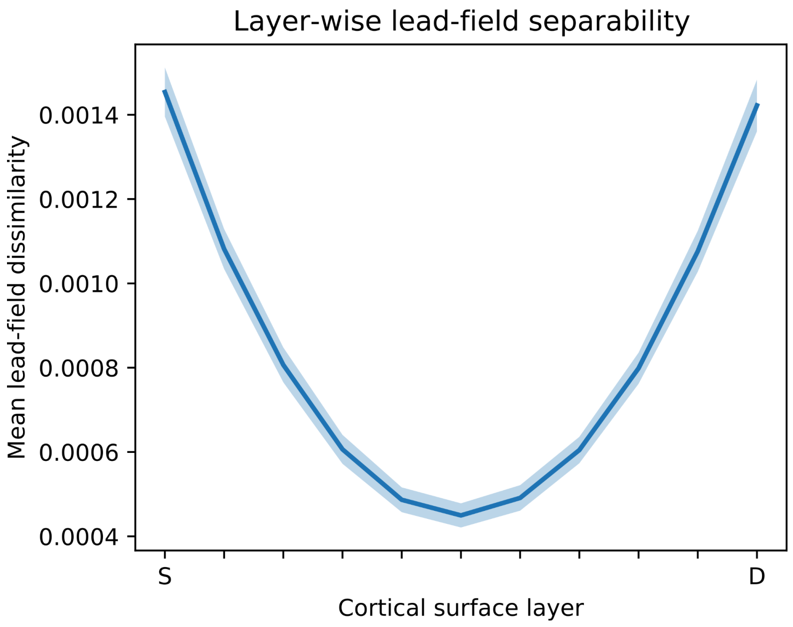


Supplementary Figure 2. **Lead-field separability across cortical depth.** Mean lead-field separability as a function of cortical surface layer, averaged across all simulated cortical locations. For each cortical vertex, lead fields were extracted for all cortical depth surfaces and normalized to unit length. Pairwise cosine dissimilarities (1 - |cosine similarity|) were then computed between the lead fields of each layer and all other layers. Lead-field separability for a given layer was defined as the mean cosine dissimilarity between that layer's lead field and the lead fields of all remaining cortical depths. Higher values therefore indicate that a layer produces more distinct sensor-level field patterns relative to other depths. The resulting profile exhibits a characteristic U-shape, with the most superficial (S) and deepest (D) cortical surfaces showing greater separability than intermediate depths. Shaded regions indicate ± SEM across cortical vertices.

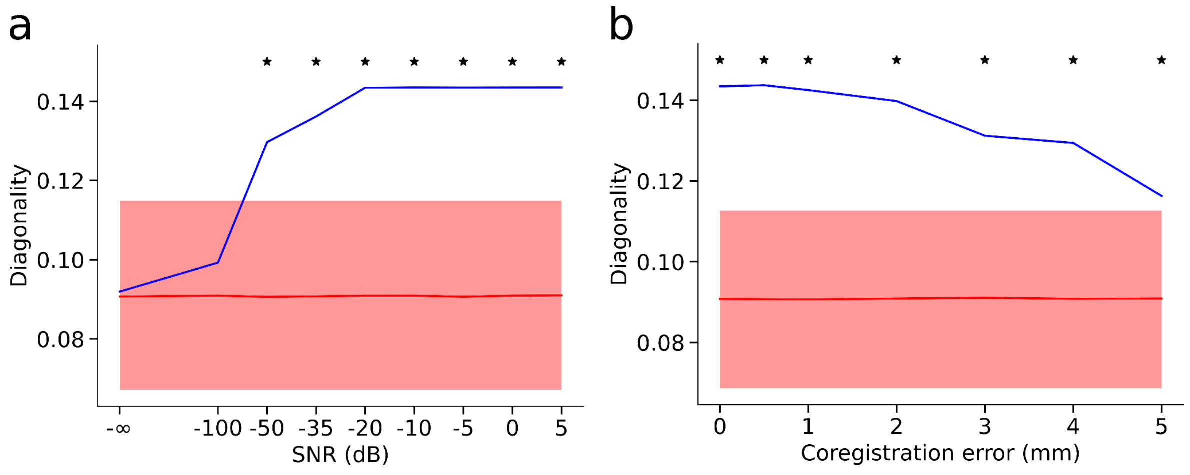


Supplementary Figure 3. **Diagonal-dominance scores for laminar model-evidence matrices across different SNR and co-registration error values. a** Varying sensor-level SNR, and **b** varying co-registration error. Blue curves show observed diagonal dominance (trace / sum of each matrix), and red curves show the mean from permuted matrices with shaded 95% CI. Black stars mark SNR or co-registration levels where observed diagonal dominance differed significantly from chance ($p < 0.05$).

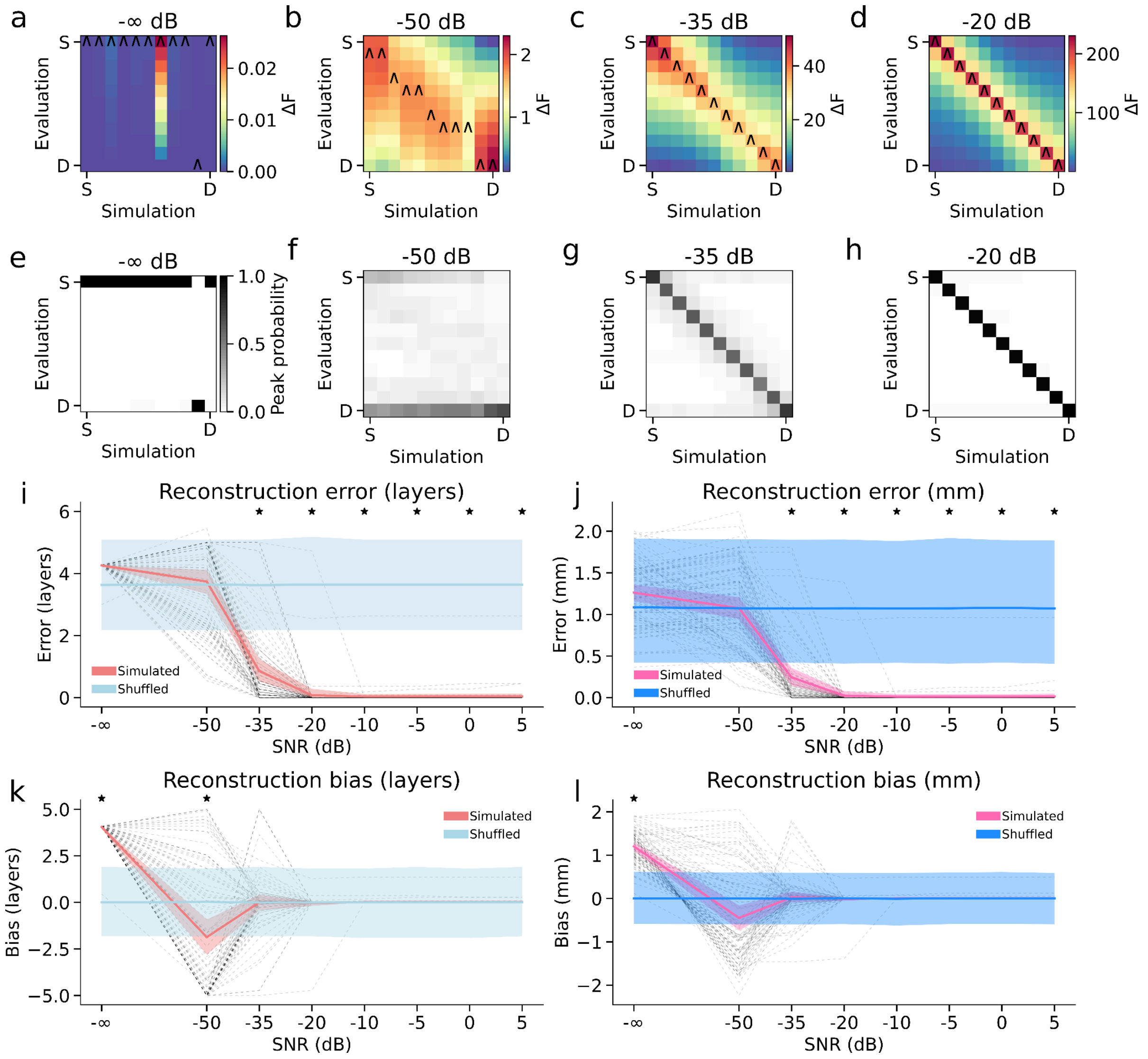


Supplementary Figure 4. **Laminar source-reconstruction accuracy with EBB as a function of SNR. a-d** Mean model evidence (free energy, ΔF) matrices (relative to the worst model; S = superficial surface layer, D = deep surface layer) for simulated signals placed on 11 cortical surfaces at four SNR levels (-∞ dB, -50 dB, -35 dB, -20 dB). Caret markers denote the forward model with peak free energy for each simulated depth. **e-h** Probability density of each forward model being the maximum-evidence solution, aggregated across cortical locations. **i, j** Laminar reconstruction error (in layers and mm). **k, l** Laminar reconstruction bias (in surface layers and millimeters), again comparing simulated and shuffled data. Shaded bands represent 95% confidence intervals, and asterisks mark SNR levels at which observed error or bias differed significantly from chance ($p < 0.05$).

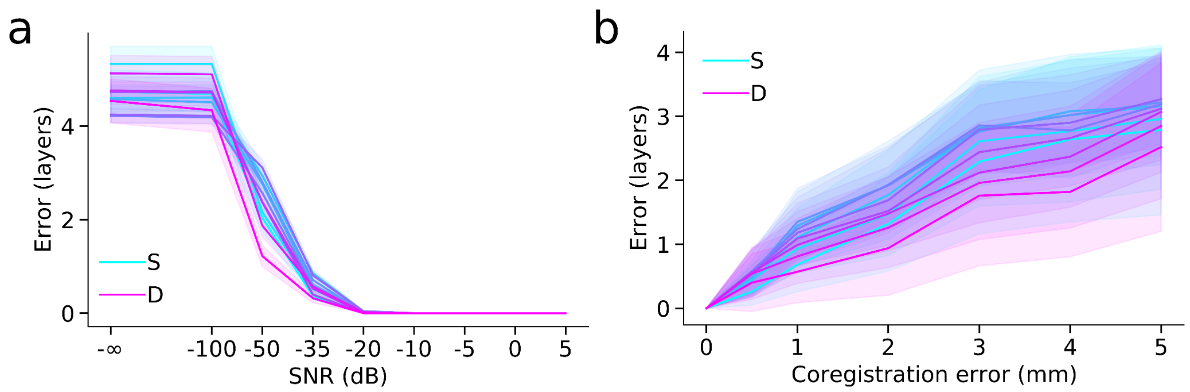


Supplementary Figure 5. **Layer-by-layer source-reconstruction error. a** Mean error as a function of SNR for each simulation surface layer, from superficial (S, light cyan) to deep (D, magenta). **b** Corresponding layer-by-layer error over increasing co-registration error.

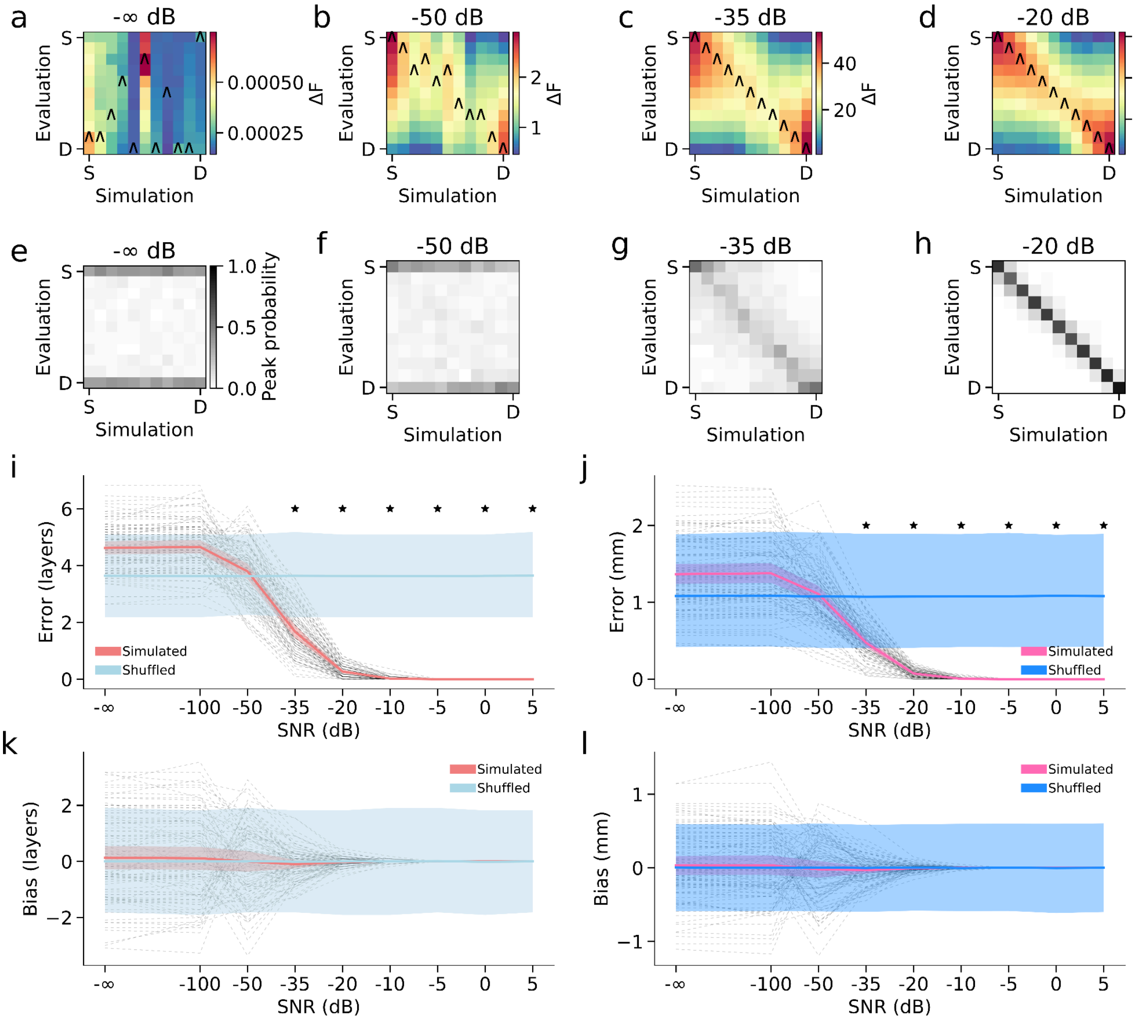


Supplementary Figure 6. **Laminar source-reconstruction accuracy as a function of SNR using pink noise. a-d** Mean model evidence (free energy, ΔF) matrices (relative to the worst model; S = superficial surface layer, D = deep surface layer) for simulated signals placed on 11 cortical surfaces at four SNR levels (-∞ dB, -50 dB, -35 dB, -20 dB). Caret markers denote the forward model with peak free energy for each simulated depth. **e-h** Probability density of each forward model being the maximum-evidence solution, aggregated across cortical locations. **i, j** Laminar reconstruction error (in layers and mm). **k, l** Laminar reconstruction bias (in surface layers and millimeters), again comparing simulated and shuffled data. Shaded bands represent 95% confidence intervals, and asterisks mark SNR levels at which observed error or bias differed significantly from chance ($p < 0.05$).

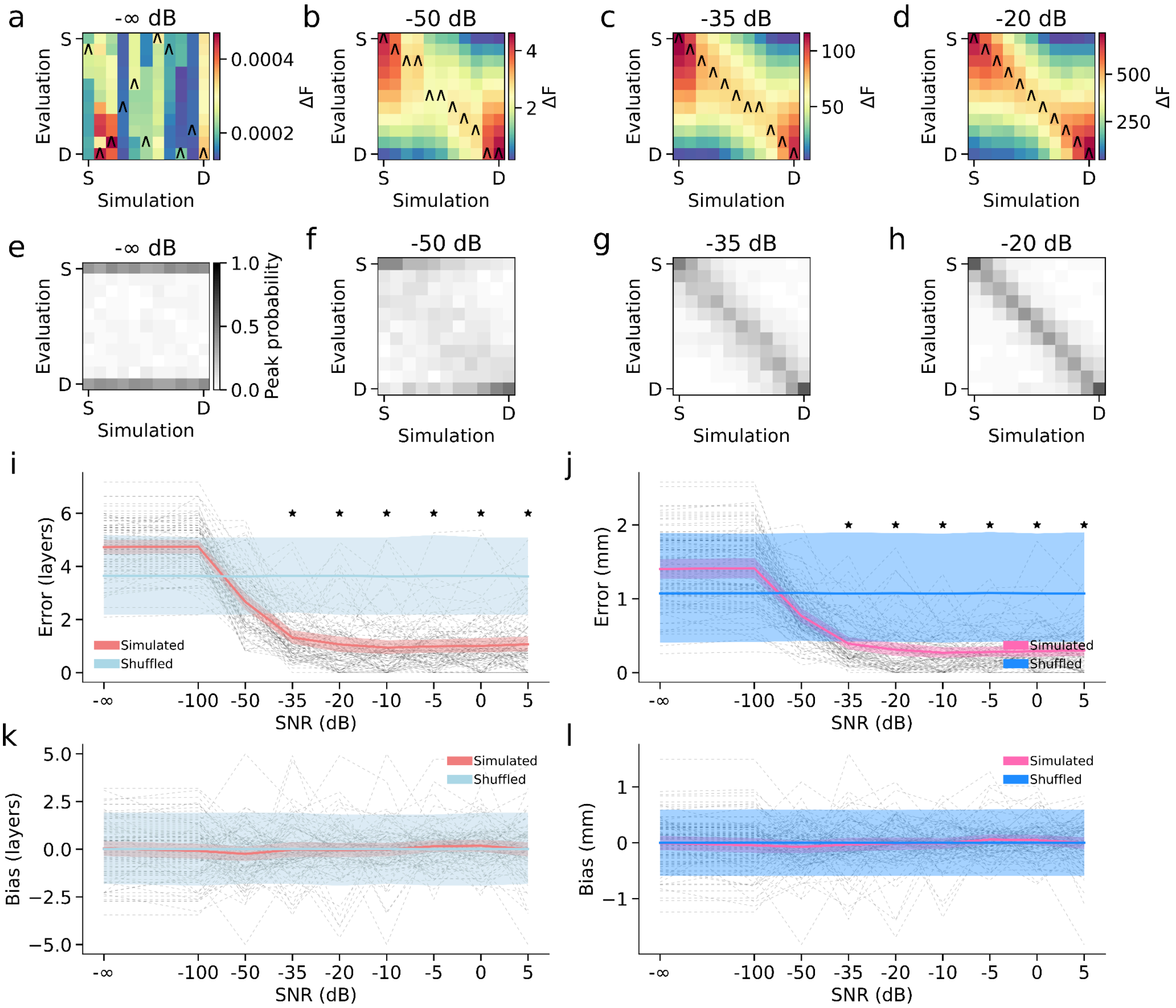


Supplementary Figure 7. **Laminar source-reconstruction accuracy as a function of SNR in the presence of additional interfering sources.** Simulations included three simultaneous distractor sources placed at random cortical locations in the middle cortical surface, each with 10% of the amplitude of the primary simulated source and band-limited (10-30 Hz) temporal profiles. **a-d** Mean model evidence (free energy, ΔF) matrices (relative to the worst model; S = superficial surface layer, D = deep surface layer) for simulated signals placed on 11 cortical surfaces at four SNR levels (-∞ dB, -50 dB, -35 dB, -20 dB). Caret markers denote the forward model with peak free energy for each simulated depth. **e-h** Probability density of each forward model being the maximum-evidence solution, aggregated across cortical locations. **i, j** Laminar reconstruction error (in layers and mm). **k, l** Laminar reconstruction bias (in surface layers and millimeters), again comparing simulated and shuffled data. Shaded bands represent 95% confidence intervals, and asterisks mark SNR levels at which observed error or bias differed significantly from chance ($p < 0.05$).

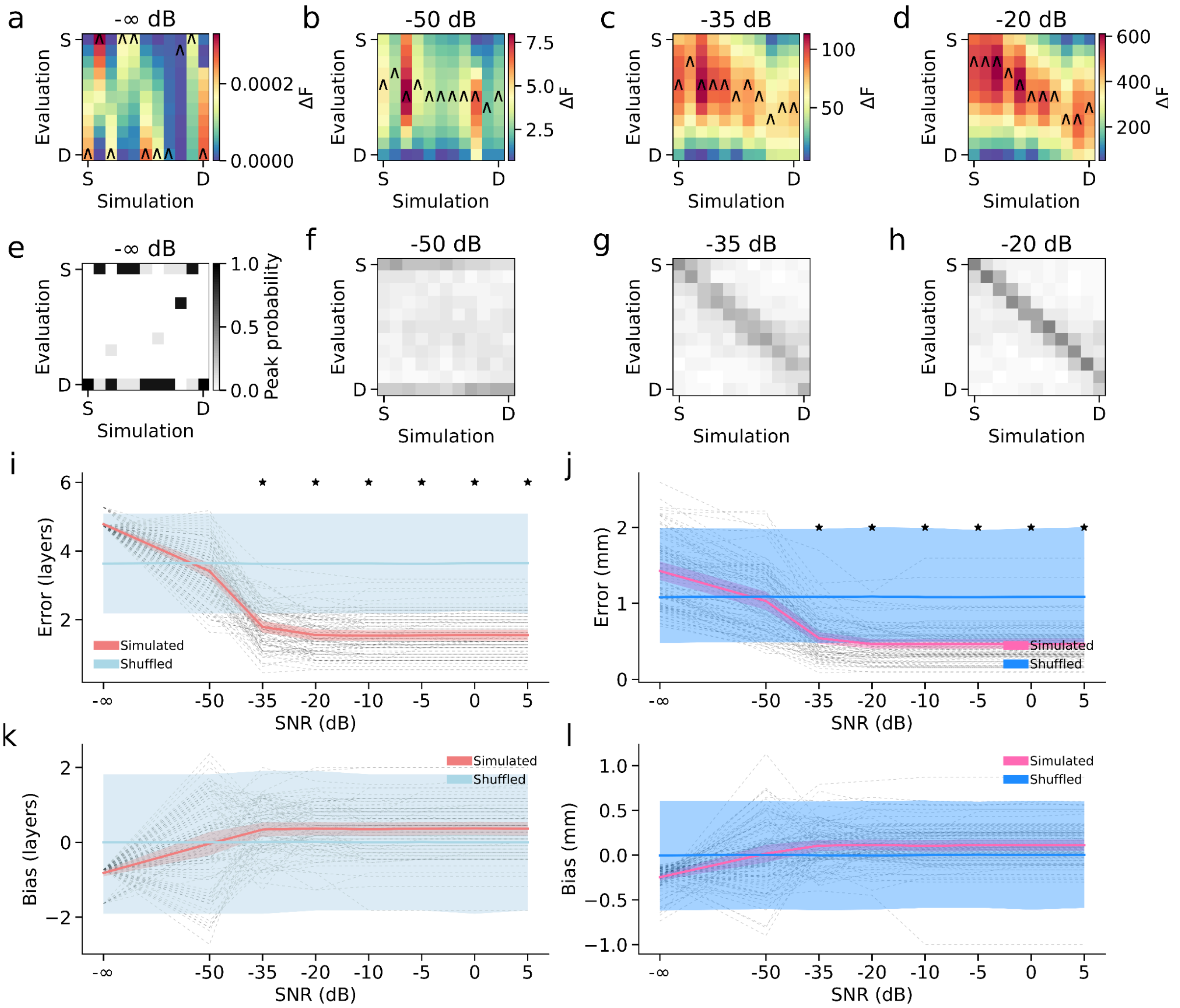


Supplementary Figure 8. **Laminar source-reconstruction accuracy as a function of SNR using mismatched cortical source spaces for simulation and inversion.** Simulated data were generated using multilayer cortical surfaces containing 32,399 vertices per layer, whereas source reconstruction was performed using the cortical source model employed in the main analyses (29,130 vertices per layer), introducing systematic mismatch between simulated and reconstructed dipole locations. **a-d** Mean model evidence (free energy, ΔF) matrices (relative to the worst model; S = superficial surface layer, D = deep surface layer) for simulated signals placed on 11 cortical surfaces at four SNR levels (-∞ dB, -50 dB, -35 dB, -20 dB). Caret markers denote the forward model with peak free energy for each simulated depth. **e-h** Probability density of each forward model being the maximum-evidence solution, aggregated across cortical locations. **i, j** Laminar reconstruction error (in layers and mm). **k, l** Laminar reconstruction bias (in surface layers and millimeters), again comparing simulated and shuffled data. Shaded bands represent 95% confidence intervals, and asterisks mark SNR levels at which observed error or bias differed significantly from chance ($p < 0.05$).

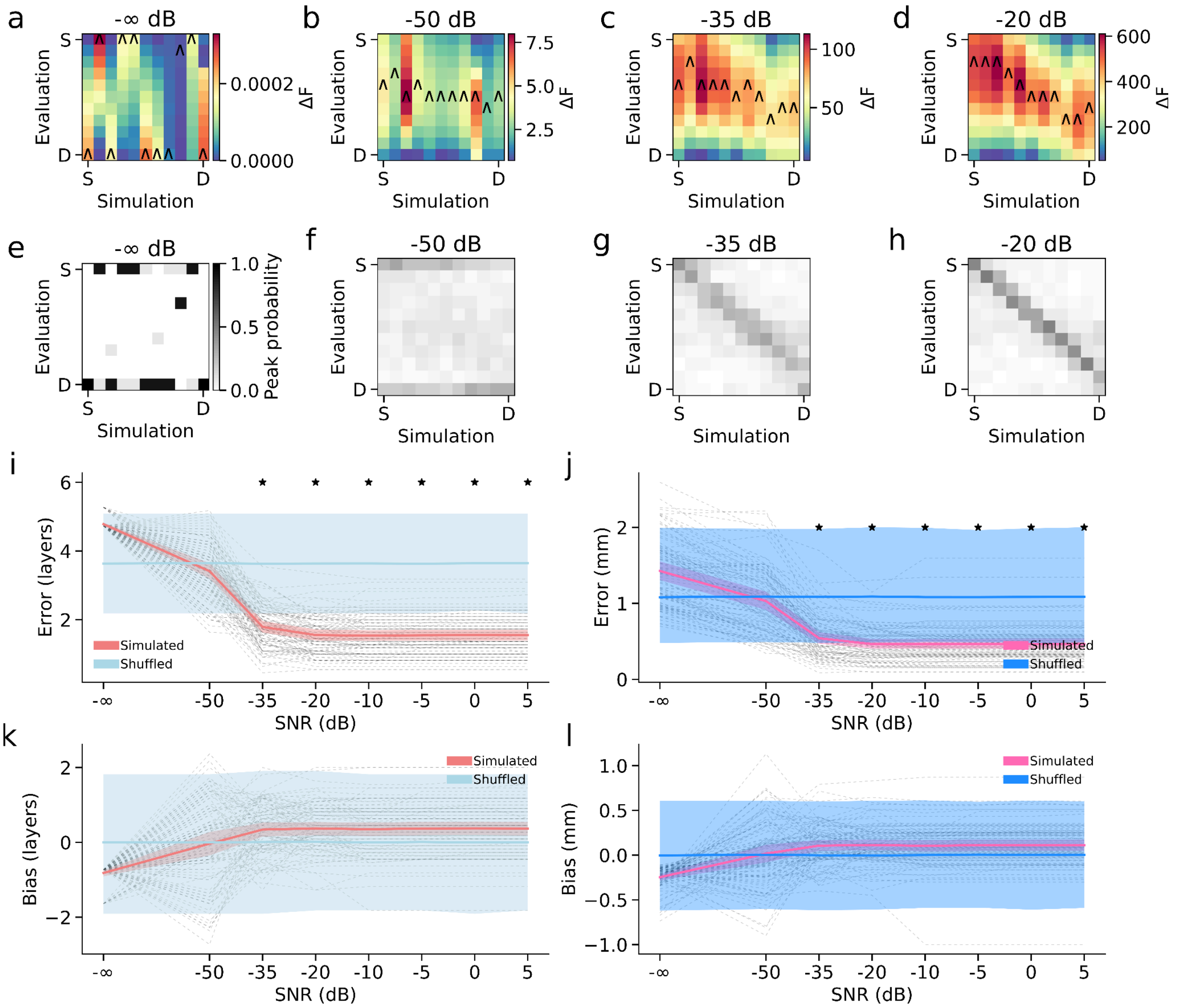


Supplementary Figure 9. **Laminar source-reconstruction accuracy as a function of SNR using mismatched forward models.** Simulated data were generated using lead fields from a boundary element model (BEM) forward model, whereas source reconstruction was performed using the same Nolte single shell forward model employed in the main analyses. **a-d** Mean model evidence (free energy, ΔF) matrices (relative to the worst model; S = superficial surface layer, D = deep surface layer) for simulated signals placed on 11 cortical surfaces at four SNR levels (-∞ dB, -50 dB, -35 dB, -20 dB). Caret markers denote the forward model with peak free energy for each simulated depth. **e-h** Probability density of each forward model being the maximum-evidence solution, aggregated across cortical locations. **i, j** Laminar reconstruction error (in layers and mm). **k, l** Laminar reconstruction bias (in surface layers and millimeters), again comparing simulated and shuffled data. Shaded bands represent 95% confidence intervals, and asterisks mark SNR levels at which observed error or bias differed significantly from chance ($p < 0.05$).

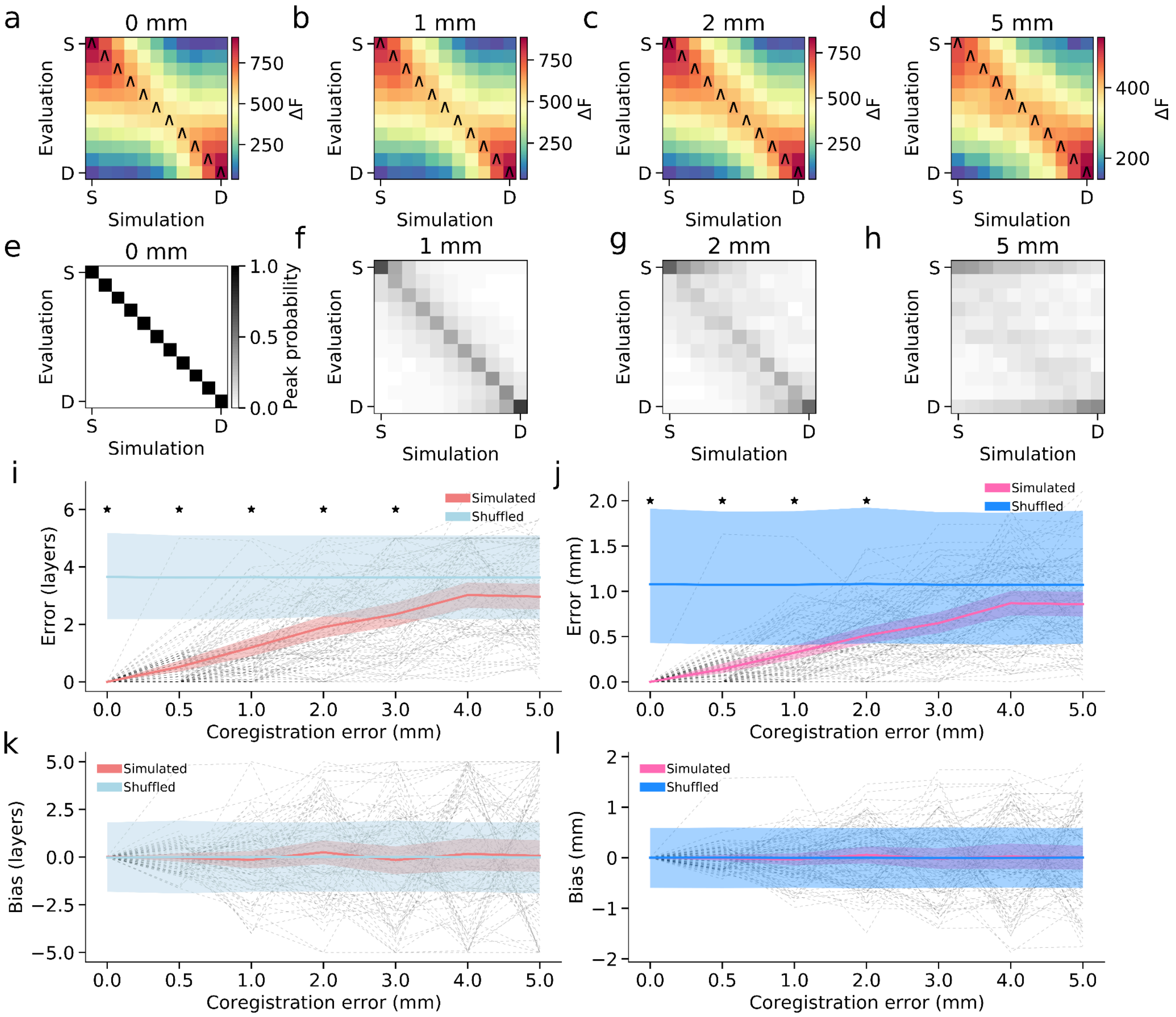


Supplementary Figure 10. **Laminar source-reconstruction accuracy as a function of co-registration error using pink noise. a-d** Mean model evidence (free energy, ΔF) matrices (relative to the worst model; S = superficial surface layer, D = deep surface layer) for simulated signals placed on 11 cortical surfaces 0, 1, 2, and 5 mm co-registration error. Caret markers denote the forward model with peak free energy for each simulated depth. **e-h** Probability density of each forward model being the maximum-evidence solution, aggregated across cortical locations. **i, j** Laminar reconstruction error (in layers and mm). **k, l** Laminar reconstruction bias (in surface layers and millimeters), again comparing simulated and shuffled data. Shaded bands represent 95% confidence intervals, and asterisks mark significance at $p < 0.05$.

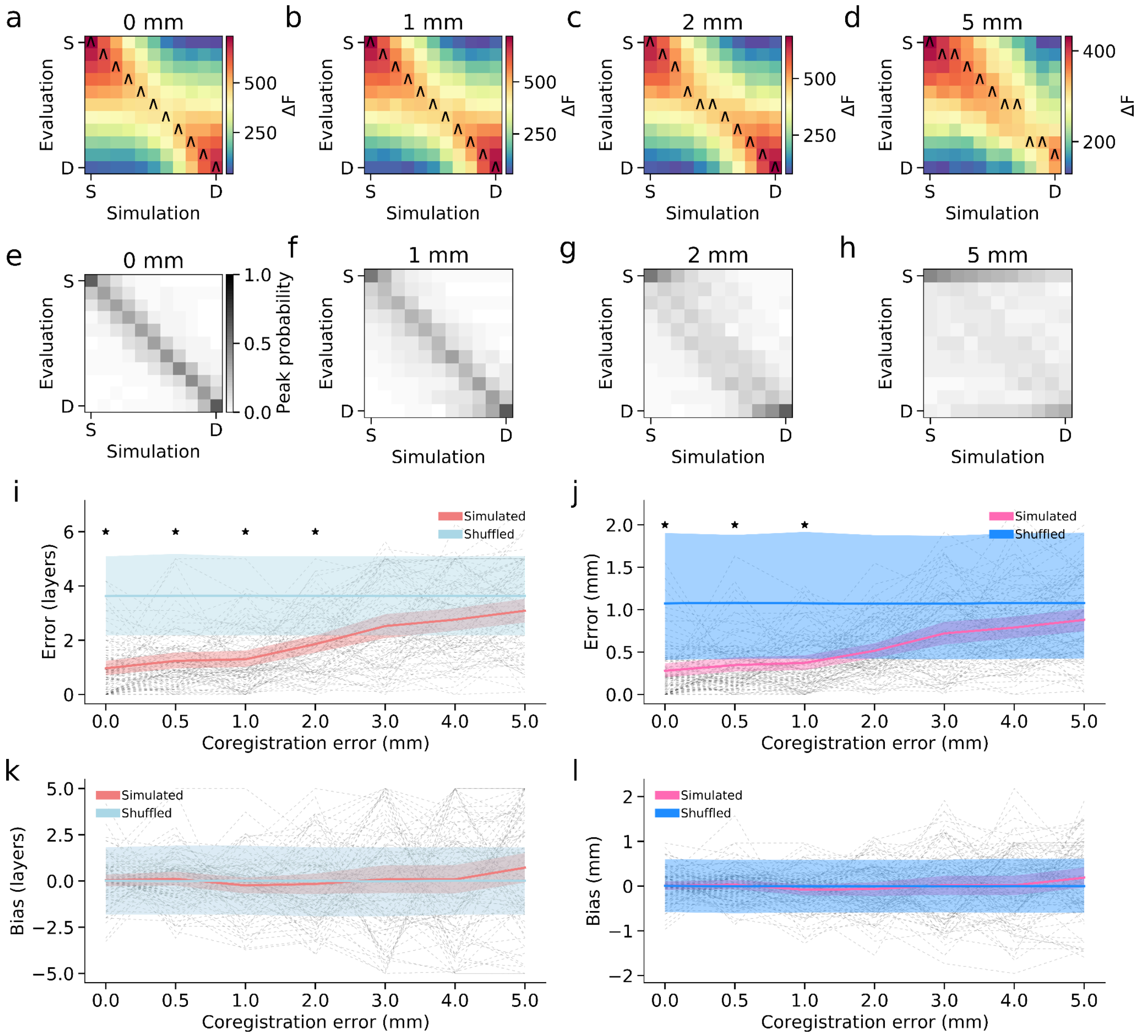


Supplementary Figure 11. **Laminar source-reconstruction accuracy as a function of co-registration error in the presence of additional interfering sources.** Simulations included three simultaneous distractor sources placed at random cortical locations in the middle cortical surface, each with 10% of the amplitude of the primary simulated source and band-limited (10-30 Hz) temporal profiles. **a-d** Mean model evidence (free energy, ΔF) matrices (relative to the worst model; S = superficial surface layer, D = deep surface layer) for simulated signals placed on 11 cortical surfaces 0, 1, 2, and 5 mm co-registration error. Caret markers denote the forward model with peak free energy for each simulated depth. **e-h** Probability density of each forward model being the maximum-evidence solution, aggregated across cortical locations. **i, j** Laminar reconstruction error (in layers and mm). **k, l** Laminar reconstruction bias (in surface layers and millimeters), again comparing simulated and shuffled data. Shaded bands represent 95% confidence intervals, and asterisks mark significance at $p < 0.05$.

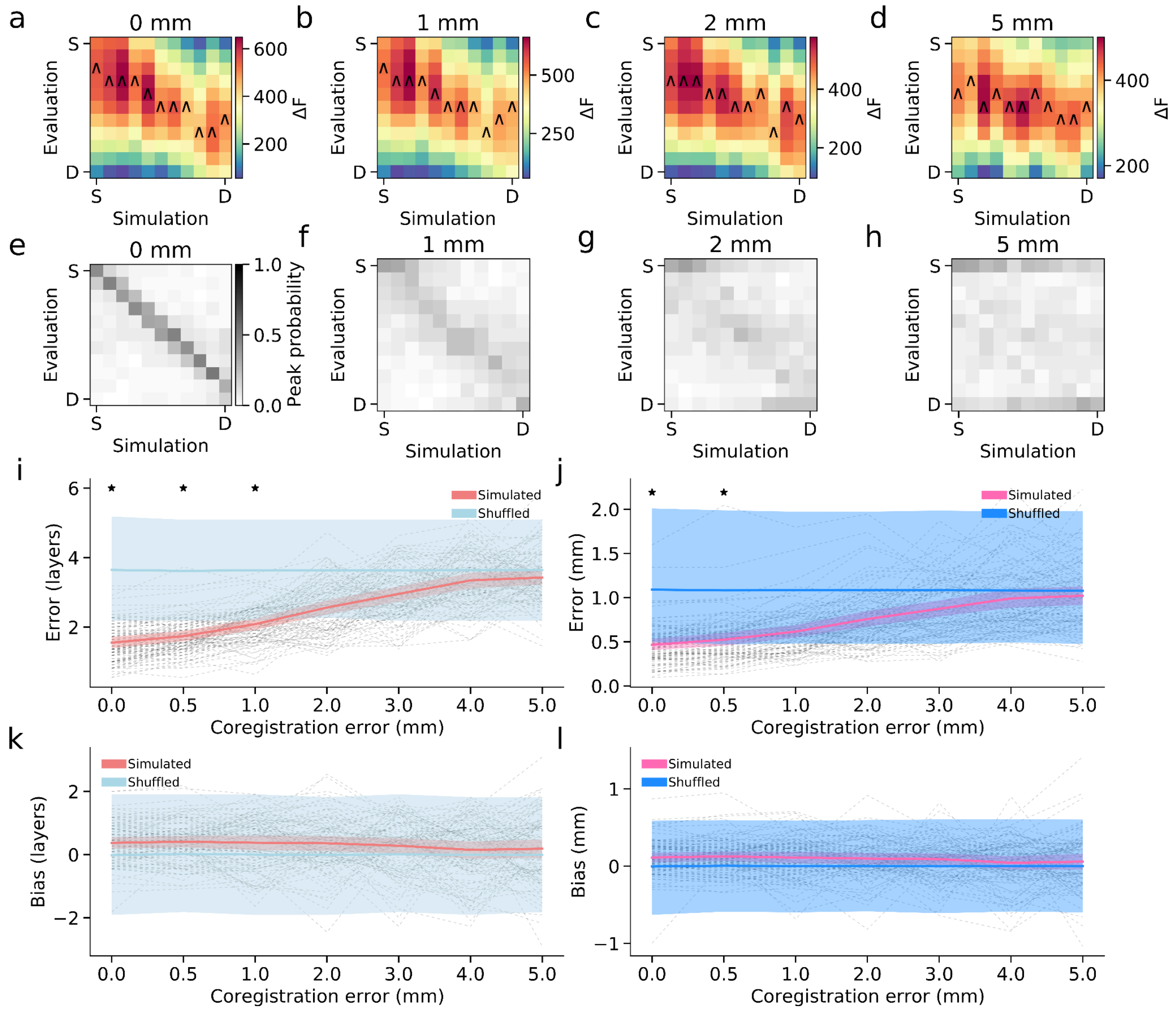


Supplementary Figure 12. **Laminar source-reconstruction accuracy as a function of co-registration error using mismatched cortical source spaces for simulation and inversion.** Simulated data were generated using multilayer cortical surfaces containing 32,399 vertices per layer, whereas source reconstruction was performed using the cortical source model employed in the main analyses (29,130 vertices per layer), introducing systematic mismatch between simulated and reconstructed dipole locations. **a-d** Mean model evidence (free energy, ΔF) matrices (relative to the worst model; S = superficial surface layer, D = deep surface layer) for simulated signals placed on 11 cortical surfaces 0, 1, 2, and 5 mm co-registration error. Caret markers denote the forward model with peak free energy for each simulated depth. **e-h** Probability density of each forward model being the maximum-evidence solution, aggregated across cortical locations. **i, j** Laminar reconstruction error (in layers and mm). **k, l** Laminar reconstruction bias (in surface layers and millimeters), again comparing simulated and shuffled data. Shaded bands represent 95% confidence intervals, and asterisks mark significance at $p < 0.05$.

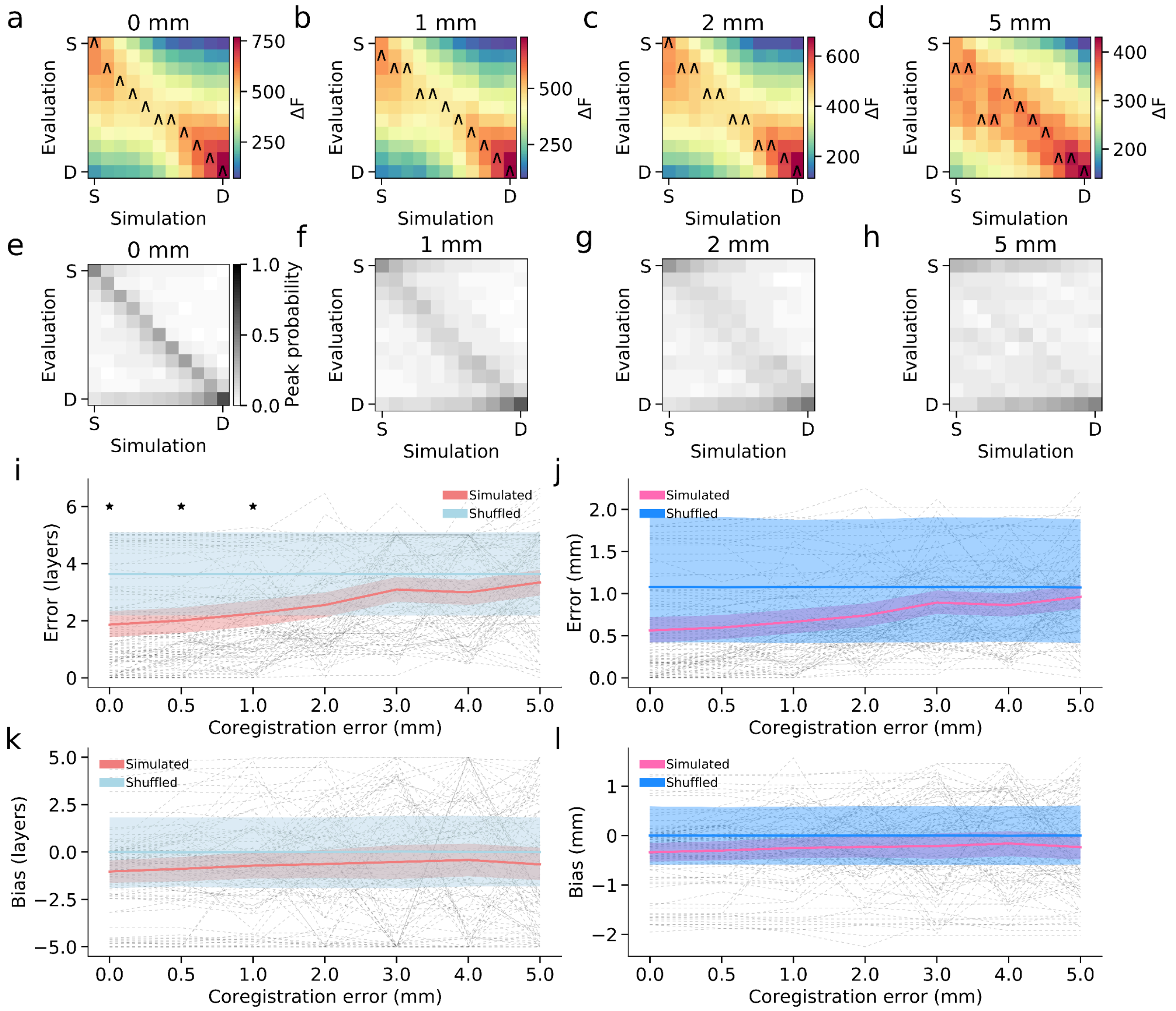


Supplementary Figure 13. **Laminar source-reconstruction accuracy as a function of co-registration error using mismatched forward models.** Simulated data were generated using lead fields from a boundary element model (BEM) forward model, whereas source reconstruction was performed using the same Nolte single shell forward model employed in the main analyses. **a-d** Mean model evidence (free energy, ΔF) matrices (relative to the worst model; S = superficial surface layer, D = deep surface layer) for simulated signals placed on 11 cortical surfaces 0, 1, 2, and 5 mm co-registration error. Caret markers denote the forward model with peak free energy for each simulated depth. **e-h** Probability density of each forward model being the maximum-evidence solution, aggregated across cortical locations. **i, j** Laminar reconstruction error (in layers and mm). **k, l** Laminar reconstruction bias (in surface layers and millimeters), again comparing simulated and shuffled data. Shaded bands represent 95% confidence intervals, and asterisks mark significance at $p < 0.05$.

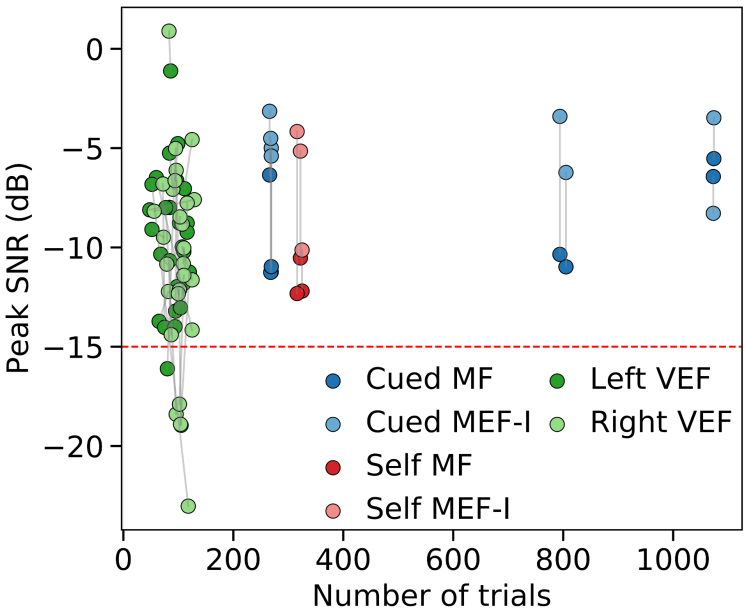


Supplementary Figure 14. **Relationship between trial count and signal-to-noise ratio (SNR) across datasets.** Sensor-level SNR (dB) within component-specific time windows plotted against the total number of trials for each participant and dataset. Data are shown separately for motor field (MF) and motor-evoked field I (MEF-I) components in the visually cued and self-paced button press datasets, and left and right stimulus presentation in the visually evoked response dataset. The gray lines connect datapoints from the same subject. SNR was computed as the ratio of the RMS of the trial-averaged signal to the RMS of trial-by-trial deviations. Considerable variability in SNR is observed across participants, datasets, and components with no simple linear relationship to trial count. The red dashed horizontal line indicates the SNR threshold (-15 dB) used for participant exclusion.

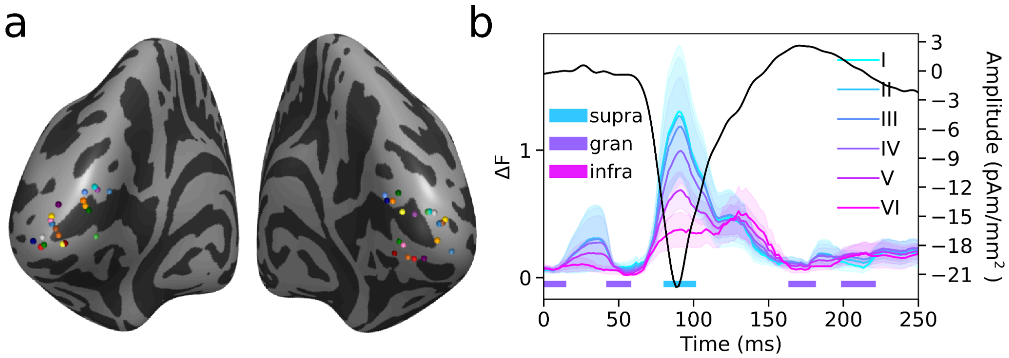


Supplementary Figure 15. **Feedforward laminar activation in ipsilateral primary visual cortex during visuospatial attention. a** Locations of the selected VEF source vertices for each participant, projected onto an inflated *fsaverage* cortical surface; each colored sphere denotes one subject. **b** Sliding-window laminar inference applied to visually evoked fields (VEFs) localized to primary visual cortex (V1) during a visuospatial attention task, analyzing responses in the hemisphere ipsilateral to the attended stimulus location. Colored traces show lamina-specific relative model evidence (ΔF, relative to the worst model at each time point) averaged across participants and shaded regions denote the SEM; the dark trace shows the corresponding middle-layer source time series. Horizontal color bars indicate, at each time point, the laminar family with the highest protected exceedance probability (PEP), grouping layers into supragranular (laminae I-III), granular (lamina IV), and infragranular (laminae V-VI) families. The ipsilateral VEF exhibits the same canonical feedforward laminar sequence observed contralaterally, with early granular dominance consistent with thalamic input, followed by supragranular dominance at the VEF peak and subsequent infragranular dominance.

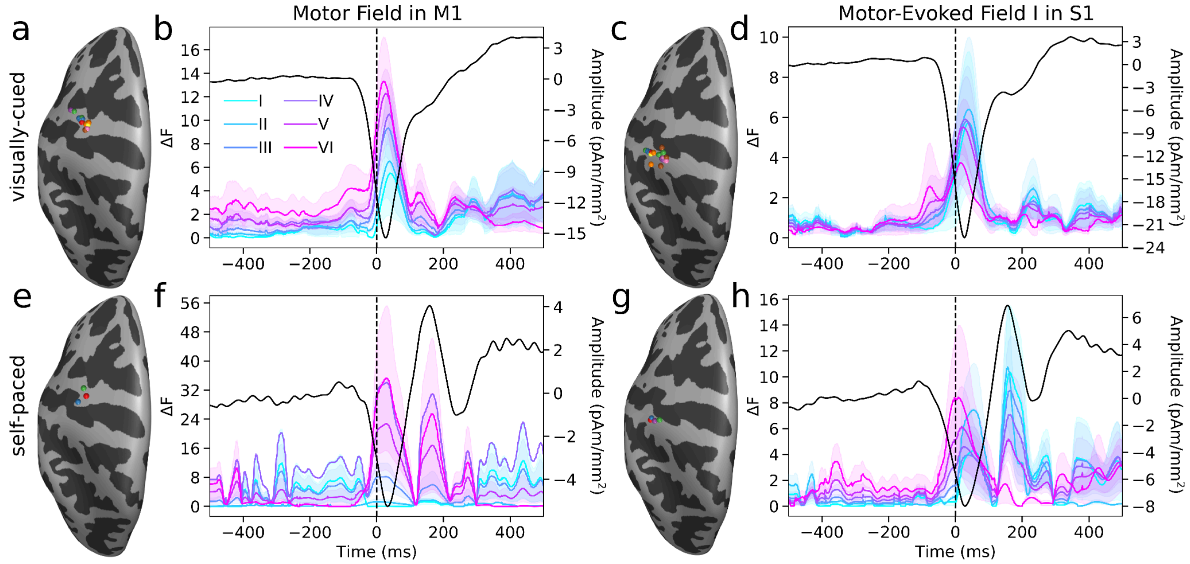


Supplementary Figure 16. **Qualitatively consistent laminar activation patterns in motor and somatosensory cortices across tasks. a** Locations of the selected motor field (MF) source vertices for each participant in the visually cued button-press task, projected onto an inflated fsaverage cortical surface; each colored sphere denotes one subject. **b** Sliding-window laminar inference for the MF in the visually cued button-press task. **c** Locations of the selected motor-evoked field I (MEFI) source vertices for each participant in the visually cued button-press task. **d** Sliding-window laminar inference for the MEFI in the visually cued button-press task. **e** Locations of the selected MF source vertices for each participant in the self-paced button-press task. **f** Sliding-window laminar inference for the MF in the self-paced button-press task. **g** Locations of the selected MEFI source vertices for each participant in the self-paced button-press task. **h** Sliding-window laminar inference for the MEFI in the self-paced button-press task. In b, d, f, and h, colored traces show lamina-specific relative model evidence (ΔF; shaded regions denote SEM) and black traces show the corresponding middle-layer source time series. Across both tasks and both cortical regions, qualitatively similar, though not identical, laminar dynamics are observed, with dominant infragranular activity surrounding movement onset in M1 followed by delayed supragranular activity, and predominantly infra- and then supra-granular responses in S1 at movement onset with additional superficial transients at later latencies.

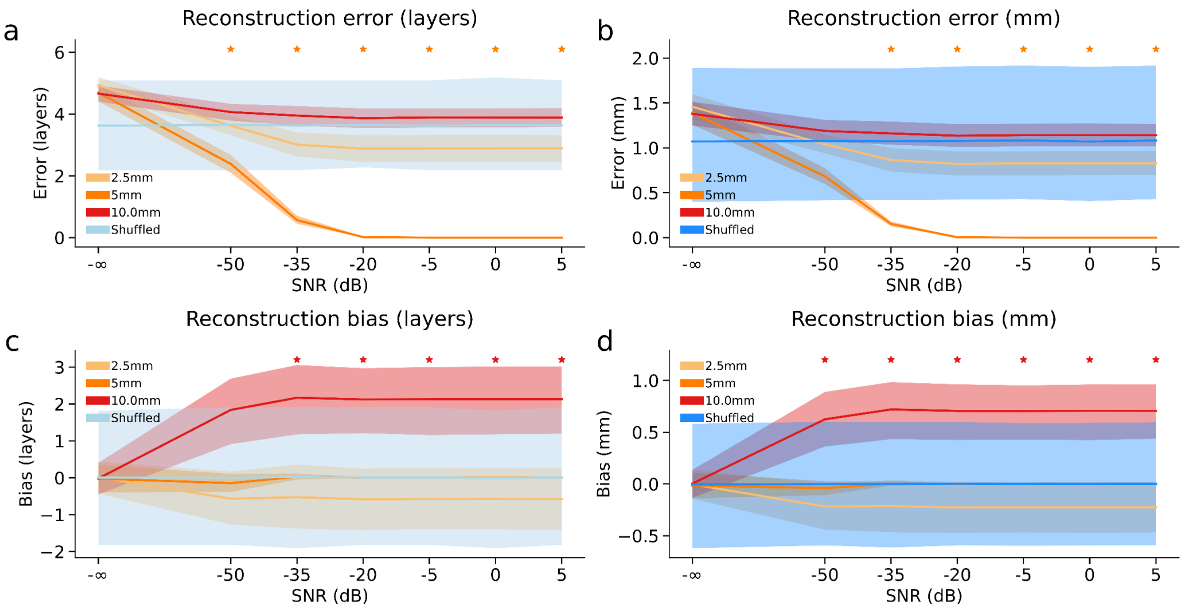


Supplementary Figure 17. **Effect of patch size mismatch on laminar source reconstruction accuracy across SNR levels. a,b** Laminar source reconstruction error (in layers and mm) falls below chance at higher SNR only when the patch size of the source reconstruction model matches that of the simulated data (5mm in these simulations). **c,d** Over-estimation of patch size biases laminar inference toward superficial layers, whereas under-estimation slightly biases toward deep layers. Shaded bands represent 95% confidence intervals, and asterisks mark SNR levels at which observed error or bias differed significantly from chance ($p < 0.05$).

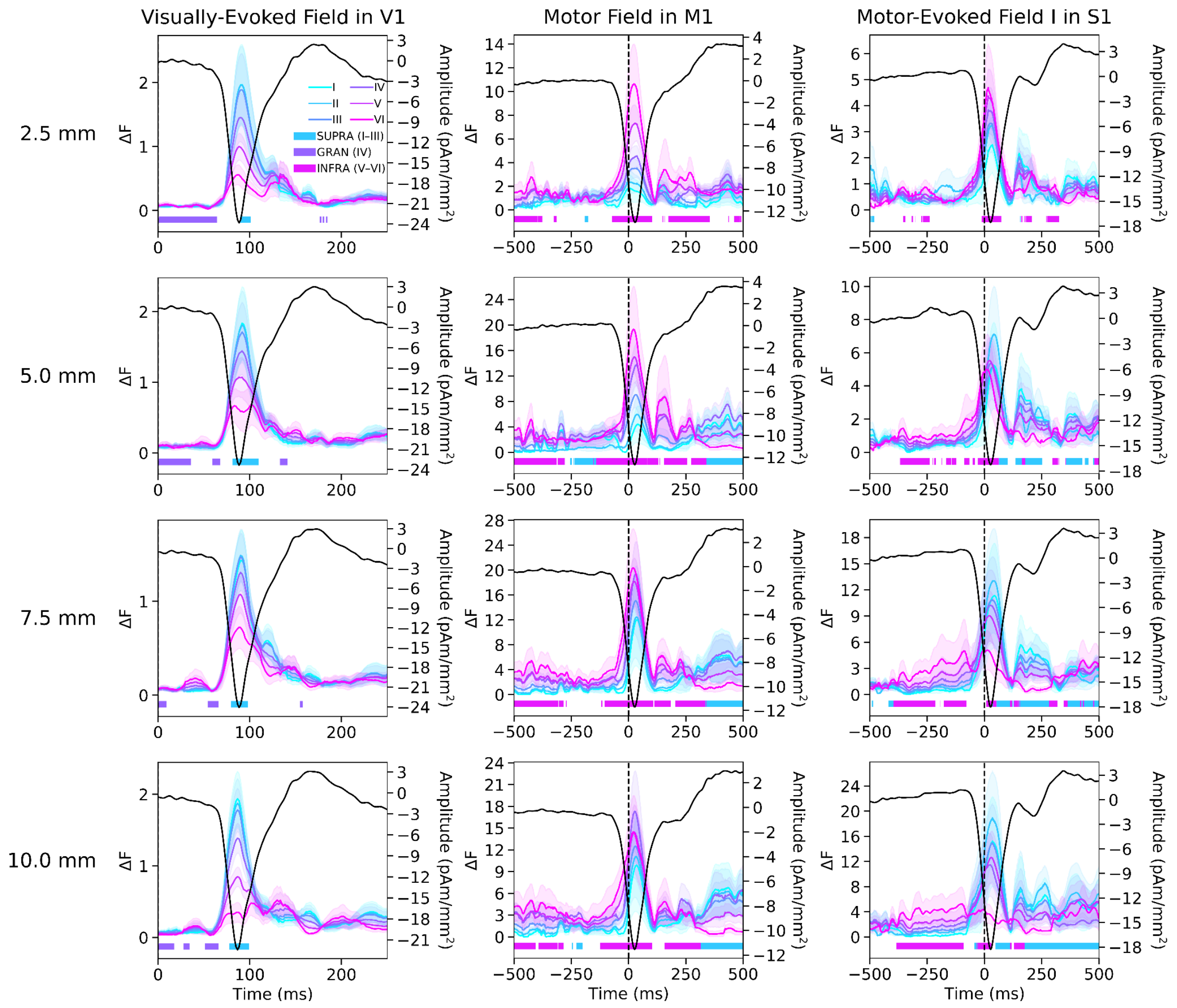


Supplementary Figure 18. **Sensitivity of empirical laminar inference to MSP patch size.** Sliding-window laminar inference results for the visually evoked field (VEF), motor field (MF), and motor-evoked field I (MEF-I; columns) using MSP patch sizes of 2.5 mm, 5 mm, 7.5 mm, and 10 mm (rows). Colored traces show lamina-specific relative model evidence (ΔF, relative to the worst model at each time point) averaged across participants, with shaded regions denoting SEM. Dark traces show the corresponding middle-layer source time series. Horizontal color bars indicate, at each time point, the laminar family with the highest protected exceedance probability (PEP), grouping layers into supragranular (laminae I-III), granular (lamina IV), and infragranular (laminae V-VI) families. Across all tested patch sizes, the principal laminar dynamics remained qualitatively stable. In the visual dataset, dominant granular and supragranular activity was preserved across patch sizes, whereas in the MF and MEF-I components the characteristic infra-/supragranular dynamics were largely unchanged, with only minor differences in the statistical significance and temporal extent of some supragranular effects. These results indicate that the empirical findings are robust to plausible variation in the assumed spatial extent of cortical sources.

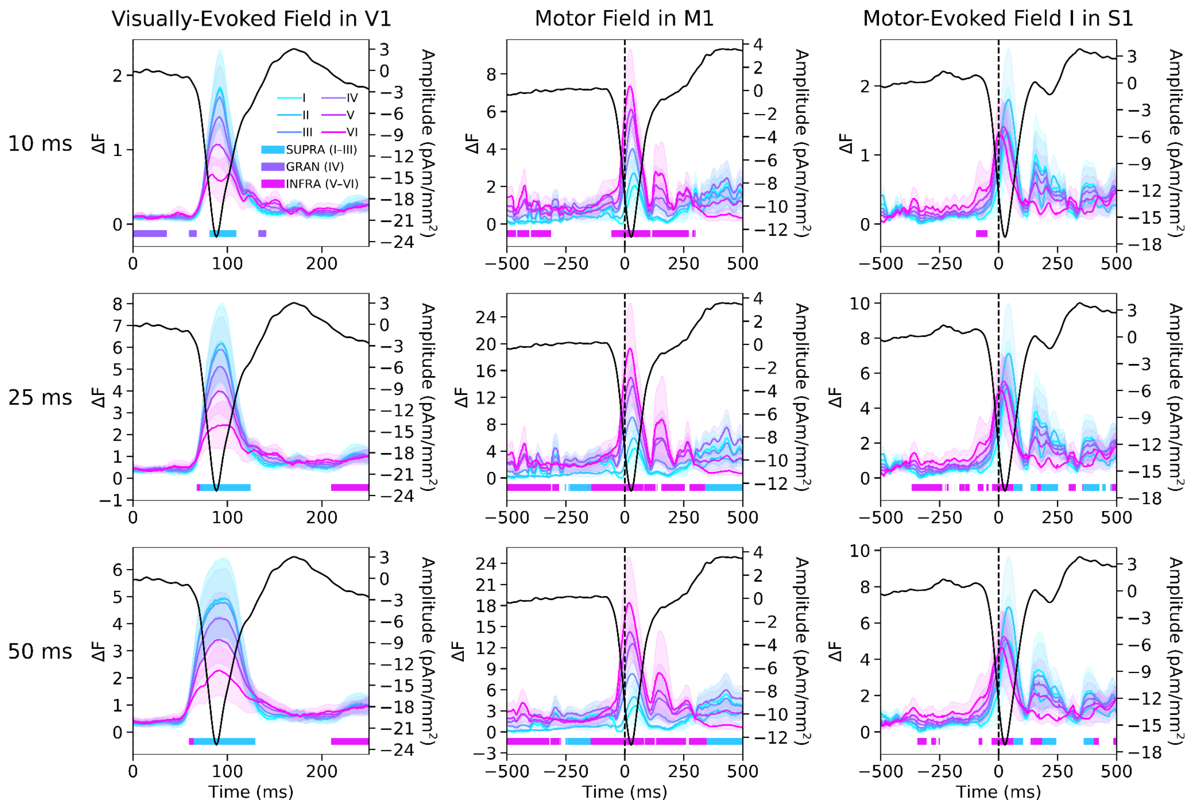


Supplementary Figure 19. **Sensitivity of empirical laminar inference to sliding-window duration.** Sliding-window laminar inference results for the visually evoked field (VEF), motor field (MF), and motor-evoked field I (MEF-I; columns) using window sizes of 10 ms, 25 ms, and 50 ms (rows). Colored traces show lamina-specific relative model evidence (ΔF, relative to the worst model at each time point) averaged across participants, with shaded regions denoting SEM. Dark traces show the corresponding middle-layer source time series. Horizontal color bars indicate, at each time point, the laminar family with the highest protected exceedance probability (PEP), grouping layers into supragranular (laminae I-III), granular (lamina IV), and infragranular (laminae V-VI) families. Short windows provide greater temporal specificity and better resolve brief laminar transients, whereas longer windows integrate information over time and produce smoother, more temporally extended evidence profiles. In the VEF, the transient granular response is most clearly resolved using a 10 ms window. In contrast, the characteristic infra-/supragranular dynamics of the MF and MEF-I are more robustly detected using 25-50 ms windows. These results demonstrate the trade-off between temporal resolution and sensitivity in sliding-window laminar inference.

### Supplementary Methods

#### *Visually evoked response dataset*

The visually evoked response dataset was collected at the CERMEP imaging platform (Lyon, France). A total of 28 healthy participants were included in the study (11 male, aged 25.7 ± 3.86 years). All participants were right-handed, as assessed by the Edinburgh Handedness Inventory[1], and reported having either normal or corrected-to-normal vision. The study protocol was in accordance with the Declaration of Helsinki, and all participants gave written informed consent which was approved by the regional ethics committee for human research (CPP Est IV - 2019-A01604–53). For each participant, structural MRI, MEG recordings, and eye-tracking data were acquired.

MRI data were acquired on a 3T Siemens Prisma scanner (CERMEP, Lyon, France) using three protocols. The first protocol acquired a T1-weighted MPRAGE sequence (TR = 2100 ms, TE = 3.33 ms, TI = 900 ms, flip angle = 8°, 1mm isotropic resolution, 256 × 256 mm field of view, 192 slices) and these images were used to create participant-specific head-casts to reduce between-session co-registration error and within-session movement during MEG recordings. Head-casts were created using a 3D scalp surface extracted from the T1-weighted volume using FreeSurfer (v6.0.0)[2], 3D-printed, placed in a dewar replica, and filled with polyurethane foam. Recesses for nasion and preauricular (LPA/RPA) fiducial coils were incorporated to enable coil locations to be recovered in the participant's native MRI space from the head-cast CAD model[3]. The second protocol acquired a sagittal 3D T2-weighted SPACE sequence (TR = 3200 ms, TE = 299 ms, flip angle = 120°, 0.8 mm isotropic resolution, 280 × 320 mm field of view). The third protocol used a quantitative multi-parameter mapping (MPM) protocol optimized for cortical surface reconstruction[4], comprising multi-echo 3D FLASH acquisitions with proton density (PD) weighting (flip angle = 6°, TE = 2.3-18.4 ms in eight echoes), T1 weighting (flip angle = 21°, TE = 2.3-18.4 ms in eight echoes), and magnetization transfer (MT) weighting (flip angle = 6°, TE = 2.3-13.8 ms in six echoes), each acquired with TR = 25 ms at 0.8 mm isotropic resolution (280 × 320 mm field of view) using parallel imaging (GRAPPA, factor 2). Calibration data included B1 mapping and gradient-echo field mapping to enable correction of transmit-field inhomogeneity and geometric distortions. Quantitative maps (PD, R1, MT, and R2*) were computed using the hMRI toolbox[5]. These maps, as well as the T2-weighted images, were aligned to the T1 used to create the head-cast, and a custom FreeSurfer-based (v6.0.0)[2] reconstruction procedure[6] was used to generate pial and white/grey matter boundary surfaces from the MPM-derived PD and T1 contrasts together with the T2-weighted volume.

MEG data was collected using a 275-channel Canadian Thin Films (CTF) MEG system in a magnetically shielded room at a sampling rate of 1200 Hz. Eye tracking data was recorded using an EyeLink 1000+ at a sampling rate of 1000 Hz. Participants performed a selective visual orientation-discrimination task while in the supine position. On each trial, participants were simultaneously presented with two vertical grating stimuli (diameter 3.5°, 12 cycles across) located laterally at an eccentricity of 3.7° of visual angle from the screen center. The task was to determine the orientation of the target grating relative to an implicit 45° tilted reference diagonal (not displayed on screen), while ignoring the distractor on the other side of the screen. Stimuli were projected on a screen positioned at approximately 50 cm from the subject. Participants responded using a two button button-box using either the index or middle finger of their right hand. The experiment was structured into four blocks of 300 trials each, with the side of the target (left/right) balanced across blocks. A central cue (a leftward or rightward pointing arrow) was presented at the beginning of each block, indicating the side of the target stimulus for that block.

MEG data was preprocessed using MNE Python (v.1.9)[7]. Raw data was band-pass filtered between 0.1 and 80 Hz, line noise was removed by a zero-phase overlap-add FIR filter at 50 Hz, and the data were then downsampled to 600 Hz and epoched around stimulus onset (-0.5 s to 0.35 s). To ensure high data quality, epochs were excluded if the participant's gaze deviated by more than 1.5° of visual angle from the fixation cross prior to stimulus onset, if they contained blinks or other pronounced ocular artifacts, or if they were associated with incorrect behavioral responses or premature responses (reaction time < 250 ms). Following these automated procedures, all remaining epochs were visually inspected, and additional trials contaminated by noise or movement-related artifacts were manually removed. Trials were analyzed separately for the left- and right-attention conditions. After preprocessing, this left an average of 187.1 trials per subject (SD = 52.5), including 89.7 trials (SD = 27.7) for the left-attention condition and 97.4 trials (SD = 28.5) for the right-attention condition.

Localization was performed in each hemisphere within primary visual cortex (Brodmann area 17), defined using each participant's FreeSurfer Brodmann Areas ex vivo atlas (BA_exvivo)[8], over 50 to 150 ms relative to the onset of the visual grating, in line with the expected visual response timing[9]. Sliding time window model comparison was performed with overlapping windows of 10 ms and 1 temporal mode. All preprocessing and laminar analysis code for this dataset can be found at https://github.com/cophyteam/laminar_visual_erf.

***Visually cued button-press dataset***

The visually cued button-press dataset was collected at the University College London Wellcome Centre for Human Neuroimaging. A total of eight healthy participants were included in the study (six male, aged 28.5 ± 8.52). The study protocol was in accordance with the Declaration of Helsinki, and all participants gave written informed consent which was approved by the UCL Research Ethics Committee (reference number 5833/001). For each participant, structural MRI, MEG recordings, and eye-tracking data were acquired.

MRI data were acquired on a 3T Siemens Magnetom TIM Trio system using the body coil for RF transmission and a 32-channel head coil for reception in two separate protocols. The first protocol generated a precise scalp image for head-cast construction[3] using a T1-weighted 3D FLASH sequence (1 mm isotropic resolution, field of view = 256 × 256 × 192 mm [A-P, H-F, R-L], TR = 7.96 ms, flip angle = 12°, single echo, bandwidth = 425 Hz/pixel, acquisition time = 3 min 42 s). A spacious 12-channel head coil was used without padding or headphones to avoid skin displacement. Participant-specific head-casts were created from a 3D scalp model extracted from the T1-weighted images using SPM[3,10]; the model was 3D-printed, placed in a dewar replica, and filled with polyurethane foam. Recesses for nasion and preauricular (LPA/RPA) fiducial coils were incorporated so that coil locations could be determined directly in native MRI space from the head-cast CAD model[3].

The second protocol used a quantitative multi-parameter mapping (MPM) protocol optimized for cortical surface reconstruction[4]. Three RF- and gradient-spoiled multi-echo 3D FLASH acquisitions with predominantly PD, T1, and MT weighting were acquired at 800 μm isotropic resolution (field of view = 256 mm H-F, 224 mm A-P, 179 mm R-L). Flip angles were 6° (PD), 21° (T1), and 6° (MT). Gradient echoes were acquired with alternating readout polarity at eight equidistant echo times ranging from 2.34 to 18.44 ms (ΔTE = 2.30 ms; bandwidth = 488 Hz/pixel); six echoes were acquired for the MT-weighted volume to maintain TR = 25 ms across contrasts. MT weighting was achieved using a 4 ms Gaussian RF pulse applied 2 kHz off-resonance with a nominal flip angle of 220° prior to excitation. Parallel imaging (GRAPPA, acceleration factor 2 in both A-P and R-L directions, 40 integrated reference lines) accelerated acquisition. Additional calibration sequences mapped transmit field inhomogeneities and corrected geometric distortions due to B0 inhomogeneity. Total acquisition time was <30 min[11,12]. Quantitative maps (PD, R1, MT, and R2*) were computed according to established procedures[4]. These maps were aligned to the T1 used to create the head-cast, and a custom FreeSurfer-based (v6.0.1)[2] reconstruction procedure was used to generate pial and white/grey matter boundary surfaces from the PD and T1 volumes of the MPMs[6].

MEG was recorded at 1200 Hz on a 275-channel Canadian Thin Films (CTF) MEG system in a magnetically shielded room. Participants performed a visually cued action-selection task (random-dot kinematogram, 2 s; 500 ms delay; left/right arrow instruction; speeded left/right button press) while seated upright and wearing a participant-specific head-cast to stabilize head position[3]. Subjects completed 1-6 sessions on different days; each session comprised three 15-min blocks of 180 trials for 540-2700 trials per participant (M = 1822.5). Two sessions each were excluded from three participants due to technical issues during data acquisition, and a total of six sessions across three participants were excluded from the motor field (MF) component analysis due to having components that localized outside of the primary motor cortex. Visual stimuli were rear-projected at ~50 cm, and responses were made on a button box with the right index (left button) or middle (right button) finger. All acquisition and task materials have been described previously[11].

Because this dataset was acquired in an upright seated position across multiple sessions, preprocessing differed slightly from that used for the visually evoked response dataset. In particular, small between-block and between-session head-position shifts required explicit movement compensation and cross-session realignment, which were not necessary in the visually evoked dataset acquired in a single supine session. In addition, line-noise removal and artifact correction procedures were adapted for the longer continuous recordings and motor-related artifacts characteristic of the button-press task.

Raw MEG was therefore movement-compensated with Maxwell filtering (temporal Signal Space Separation, tSSS)[13,14] using continuously tracked head-position indicator (cHPI) coils. For each run, cHPI coil locations were extracted and transformed from CTF device coordinates to MNE's device frame; time-resolved head position was then estimated, and we generated quality-control plots of coil trajectories, inter-coil distances, and head-position traces/fields. The device→head transform from the first run of each session was saved and used as the common destination so that subsequent runs were realigned to the same head coordinate frame (three blocks referenced to the first). SSS was applied in head coordinates with temporal extension (st_duration = 10 s) and a fixed spherical origin at [0, 0, 0.04] m (head frame). This procedure yields a single movement-corrected raw file per block that is co-registered across blocks within a session.

Data were then downsampled to 600 Hz and line noise was removed using an iterative ZapLine procedure applied until residual 50 Hz power was eliminated[15]. Independent component analysis (ICA; Infomax, 25 components) was fit on a copy of the data bandpass-filtered between 1 and 60 Hz. Components reflecting ocular, cardiac, or muscle artifacts were manually identified and the corresponding unmixing matrix was then applied to the original, unfiltered data to remove them. The cleaned data were then bandpass-filtered between 0.1 and 30 Hz for event-related field analysis and epoched around button press events (-2 s to 2 s). Trials with missing responses were excluded. Epochs from each block were then concatenated within session, and the concatenated epochs were averaged to obtain motor ERFs for laminar inference (yielding one session-level average per subject).

For the motor field (MF) component of the motor ERF, localization was performed in the left precentral gyrus, defined using each participant's FreeSurfer Desikan-Killiany cortical parcellation, over -75 to -25 ms relative to the button press. For the motor-evoked field I (MEFI) component, localization was performed within the left postcentral gyrus, defined analogously, using a post-movement window of 15 to 55 ms. All preprocessing and laminar analysis code for this dataset can be found at https://github.com/danclab/laminar_motor_erf.

***Self-paced button-press dataset***

The self-paced button-press dataset was collected at the CERMEP imaging platform (Lyon, France). Data were collected from five neurologically healthy adults (one male; 32.4 ± 5.9 y). The study protocol was in accordance with the Declaration of Helsinki, and all participants gave written informed consent which was approved by the regional ethics committee for human research (CPP Sud-Méditerranée V - 2021-A00042-39). For each participant, structural MRI and MEG data were acquired.

MRI data were acquired using either a 3T Siemens Prisma system (CERMEP, Lyon, France) or a 3T Siemens Magnetom Vida-XT-64 system (Numaris/X VA20A-04T4; University of Miami, Coral Gables, FL, USA). The structural protocol comprised: (i) a sagittal 3D T1-weighted MPRAGE (TR = 2530 ms, TI = 1100 ms, flip angle = 9°, 0.8 mm isotropic, GRAPPA acceleration factor 2, TE = 3.41 ms in Lyon and 3.42 ms in Miami); (ii) a sagittal 3D T2-weighted SPACE sequence (TR = 3200 ms, TE = 409 ms, 0.8 mm isotropic, variable flip angle mode “T2 var”, acceleration implemented using compressed sensing [factor 3.0] in Miami and GRAPPA/CAIPIRINHA in Lyon to achieve comparable scan duration); and (iii) two sagittal multi-echo 3D FLASH sequences (TR = 20 ms, eight echoes, TE = 1.87-14.59 ms) acquired at flip angles of 5° and 30° with 1 mm isotropic resolution (GRAPPA factor 2). The FLASH images were processed in FreeSurfer to synthesize a 5° FLASH volume, and the resulting high-SNR synthetic image was used to derive the scalp surface for alignment of 3D head scans. FreeSurfer (v6.0.1)[2] was used to reconstruct the cortical pial and white matter boundary surfaces from the T1- and T2-weighted volumes.

MEG data were recorded using the same CTF system used to record the visually evoked response dataset at 1200 Hz. In a single session, participants completed two blocks of an auditory oddball task followed by two blocks of a multi-tone masker task, described in detail elsewhere[16]. Between trials, participants initiated the next trial by pressing a button with their right index finger on a response box. Analyses in the present study focused exclusively on these self-paced button-press events, which occurred independently of auditory stimulation.

Participants wore individualized head-casts to stabilize head position during MEG acquisition[3] while in the supine position. Unlike the visually cued button-press dataset, the fiducial coils (nasion, LPA, RPA) were taped directly to the participant’s skin to track head motion relative to the cast. To recover the coil positions in native MRI space, each participant’s head and fiducials were digitized using an Artec Eva handheld 3D scanner. The resulting 3D mesh was cropped to the facial region and co-registered to the MRI-derived scalp surface (from the synthetic FLASH image) using a combination of fiducial-based alignment and iterative closest point (ICP) registration implemented in MNE-Python. The final fiducial coordinates were used for MEG-MRI co-registration in subsequent laminar source reconstruction analyses. Each session comprised 316-332 button presses per participant (M = 328.8).

Preprocessing procedures were exactly the same as those used for the visually cued button-press dataset, with the exception that because the self-paced responses exhibited greater trial-to-trial variability in movement timing and associated artifacts, *autoreject*[17] was applied to the session epochs to automatically detect and repair residual artifacts using cross-validated consensus thresholds (consensus = 0-1, n_interpolate = [1, 4, 32]) before averaging over trials.

Localization of the MF and MEFI components of the motor ERF used exactly the same anatomical ROIs (left precentral and postcentral gyri) and time windows used for the visually cued button-press dataset. Two participants were excluded from the MF analysis and one from the MEFI analysis due to having components that localized outside of the hand area of the primary motor or somatosensory cortex, respectively. All preprocessing and laminar analysis code for this dataset can be found at https://github.com/danclab/auditory_laminar.

***Supplementary References***


1. Oldfield, R. C. The assessment and analysis of handedness: The Edinburgh inventory. *Neuropsychologia* **9**, 97–113 (1971).
2. Fischl, B. FreeSurfer. *NeuroImage* **62**, 774–781 (2012).
3. Meyer, S. S. *et al.* Flexible head-casts for high spatial precision MEG. *J. Neurosci. Methods* **276**, 38–45 (2017).
4. Weiskopf, N. *et al.* Quantitative multi-parameter mapping of R1, PD(*), MT, and R2(*) at 3T: a multi-center validation. *Front. Neurosci.* **7**, 95 (2013).
5. Tabelow, K. *et al.* hMRI–A toolbox for quantitative MRI in neuroscience and clinical research. *Neuroimage* **194**, 191–210 (2019).
6. Carey, D. *et al.* Quantitative MRI Provides Markers Of Intra-, Inter-Regional, And Age-Related Differences In Young Adult Cortical Microstructure. *bioRxiv* https://doi.org/10.1101/139568 (2017) doi:https://doi.org/10.1101/139568.
7. Gramfort, A. *et al.* MEG and EEG data analysis with MNE-Python. *Front. Neuroinformatics* **7**, 267 (2013).
8. Fischl, B. Estimating the Location of Brodmann Areas from Cortical Folding Patterns Using Histology and Ex Vivo MRI. in *Microstructural Parcellation of the Human Cerebral Cortex* (eds Geyer, S. & Turner, R.) 129–156 (Springer Berlin Heidelberg, Berlin, Heidelberg, 2013). doi:10.1007/978-3-642-37824-9_4.
9. Bijanzadeh, M., Nurminen, L., Merlin, S., Clark, A. M. & Angelucci, A. Distinct laminar processing of local and global context in primate primary visual cortex. *Neuron* **100**, 259–274 (2018).

10. Troebinger, L. *et al.* High precision anatomy for MEG. *NeuroImage* **86**, 583–591 (2014).

11. Bonaiuto, J. *et al.* Lamina-specific cortical dynamics in human visual and sensorimotor cortices. *eLife* **7**, e33977 (2018).

12. Bonaiuto, J. *et al.* Non-invasive laminar inference with MEG: Comparison of methods and source inversion algorithms. *NeuroImage* **167**, 372–383 (2018).

13. Taulu, S. & Simola, J. Spatiotemporal signal space separation method for rejecting nearby interference in MEG measurements. *Phys. Med. Biol.* **51**, 1759 (2006).

14. Taulu, S., Kajola, M. & Simola, J. Suppression of Interference and Artifacts by the Signal Space Separation Method. *Brain Topogr.* **16**, 269–275 (2004).

15. de Cheveigné, A. ZapLine: A simple and effective method to remove power line artifacts. *NeuroImage* **207**, 116356 (2020).

16. Dykstra, A. R. & Gutschalk, A. Does the mismatch negativity operate on a consciously accessible memory trace? *Sci. Adv.* **1**, e1500677 (2015).

17. Jas, M., Engemann, D. A., Bekhti, Y., Raimondo, F. & Gramfort, A. Autoreject: Automated artifact rejection for MEG and EEG data. *NeuroImage* **159**, 417–429 (2017).
